# When AI encounters natural history: Morphological OTUs reshape our understanding of Earth’s life

**DOI:** 10.64898/2026.04.28.721370

**Authors:** Zhihong Zhan, Maolin Ye, Michael C. Orr, Weiqiang Chen, Xue Liu, Ling Yue, Xin Sun, Feng Zhang

## Abstract

Biodiversity can be quantified only after organisms are assigned to reproducible units, yet most individuals encountered in nature lack reliable species-level identifications. Molecular operational taxonomic units can organize unnamed diversity, whereas image-based approaches generally depend on predefined species levels. Here, we show that operational biodiversity units can be derived directly from phenotypes. We developed morphOTU, a framework combining self-supervised representation learning, operational metric supervision, and adaptive hierarchical clustering to organize specimen images in continuous phenotypic space. Across five benchmark datasets comprising flowers, wood anatomy, and beetle habitus, morphOTUs recovered coherent species-level structure and produced α-diversity estimates close to those obtained from expert identifications. This structure remained informative when species were excluded from representation learning, when labeled data were sparse, and per-species sampling was limited. In a heterogeneous field-survey dataset of 4,717 insects representing 269 species across 12 orders, fine-tuning on only 28 common species produced diversity estimates close to expert labels (Shannon index, 3.75 versus 3.53). Visual explanations localized variation to biologically meaningful structures, including body outliers, surface sculpture, floral symmetry, and wood vessels. Morphological units therefore provide an operational layer for organizing and quantifying biodiversity before, alongside, or in the absence of formal species names.

## Introduction

Biodiversity becomes quantitatively tractable only when individual organisms can be assigned to reproducible biological units. Although biodiversity encompasses variation from genes to ecosystems, named species remain the principal units through which organisms are counted, compared, monitored, and prioritized in most ecological and conservation analyses (*1*, *2*).

Taxonomy, therefore, provides essential infrastructure for biodiversity science through the hierarchical organization established by the Linnaean system, and the nomenclatural stability promoted by the ICZN, ICN, ICNP, and ICVCN allows biological observations to be connected, retrieved, and communicated as structured scientific knowledge (*3–7*). Yet a large proportion of the organisms encountered in biodiversity surveys, natural history collections, and environmental samples cannot be identified reliably to species, particularly within hyperdiverse invertebrate groups and other poorly featured lineages (*8–10*).

This limitation reflects the long-recognized taxonomic impediment, including extremely high species richness, dependence on specialized expertise, slow and labor-intensive workflows, and uneven taxonomic capacity across organismal groups and geographic regions (*8*, *9*, *11*). The approximately 2.2 million cataloged species represent only a fraction of the diversity that may exist (*10*, *12–14*), while formal description and expert identification cannot keep pace with the rate at which biodiversity observations are being generated. Uneven taxonomic capacity and taxonomic biases further leave many species-rich groups poorly represented in biodiversity knowledge systems (*15–17*). Unidentified specimens are eventually often excluded from quantitative analyses. Without reproducible operational units, observable biodiversity remains difficult to count, track, and incorporate into standardized ecological assessment.

Molecular operational taxonomic units (OTUs) partially overcome such limitations by organizing organisms according to genetic similarity without requiring formal species names (*18*). DNA barcoding and metabarcoding have enabled high-throughput biodiversity assessment from individuals, bulk samples, and environmental DNA (*19–22*), while denoising methods such as DADA2 and UNOISE infer reproducible sequence variants without requiring taxonomic assignment, although incomplete reference databases may subsequently limit their annotation (*23*, *24*).

However, molecular workflows require biological materials, laboratory and sequencing infrastructure, and appropriate preservation and specialized markers. They can also be affected by amplification bias and incomplete reference libraries (*25–28*), leaving many detected lineages without reliable taxonomic assignments and contributing to the accumulation of unnamed “dark taxa” in molecular datasets (*29*, *30*). Moreover, molecular OTUs are not always readily connected to the visible phenotypes represented in field observations, specimen images, and natural history collections.

Computer vision offers a complementary opportunity by extracting taxonomic and phenotypic information from rapidly expanding image repositories (*31*). Deep-learning models can recognize diagnostic morphological features and achieve high species-identification accuracy for well-represented taxa (*32–34*). Nevertheless, most image-based systems operate under closed-set supervision: they assign images to a predefined catalogue of labeled species but do not construct reproducible units for taxa absent from training or not yet described. They may also focus on background noise or imaging artifacts rather than organismal phenotypes (*35*, *36*). Zero-shot and few-shot methods can extend recognition beyond densely labeled training sets (*37–40*), but they generally predict predefined taxonomic categories rather than delineating de novo, abundance-based operational units from unlabeled assemblages.

An alternative is to treat phenotype itself as the basis of an operational biodiversity unit. Rather than assigning images exclusively to named species, organisms could be positioned in a continuous, high-dimensional phenotypic space and partitioned into reproducible morphological groups that can be counted and compared among samples. Such units would parallel molecular OTUs while remaining non-destructive, image-traceable, and accessible to expert inspection.

Distinct evolutionary lineages may remain morphologically cryptic (*41*). Moreover, image-derived distances may also be influenced by specimen orientation, resolution, illumination, preservation, background, and anatomical view. A practical framework must learn organism-centered features, accommodate lineage-specific distance structures, distinguish biological variation from imaging noise, and remain informative when many species lack labeled representatives. It must also generate abundance-based units that preserve ecological quantities, rather than merely producing visually plausible clusters, while retaining interpretable evidence for expert evaluation. To our knowledge, no general framework currently integrates these requirements to derive operational biodiversity units directly from organismal images.

Here, we introduce morphOTU, an image-based framework for organizing and quantifying biodiversity in phenotypic space. MorphOTU combines OTU-Former, a two-stage architecture based on self-supervised representation learning and optional supervised metric fine-tuning, with hierarchical clustering and dynamic threshold scanning. It transforms specimen images into phenotypic embeddings and partitions them using lineage-specific thresholds to generate morphological operational taxonomic units (morphOTUs).

We evaluated the framework across five benchmark datasets spanning flowers, wood transverse anatomy, and beetle dorsal habitus, and tested its robustness to unseen species, sparse taxonomic labels, limited per-species sampling, cross-lineage transfer, and image degradation. We further applied it to a heterogeneous field-survey dataset containing 4,717 insect images representing 269 species across 12 orders. We asked whether image-derived units recover reference species-level structure, remain informative when species labels are unavailable or when species are absent from representation learning, reproduce species-based α-diversity estimates, and correspond to biologically meaningful morphological traits.

Our results show that morphOTUs provide an operational layer between raw images and formal taxonomy, allowing unidentified organisms to be provisionally organized, counted, and compared while retaining a direct connection to observable phenotype. They are not substitutes for formal taxonomy or integrative species delimitation, but reproducible phenotypic units that can support biodiversity assessment before, alongside, or in the absence of complete species-level identification.

## Results

### Overview of the MorphOTU workflow

Across five standardized benchmark datasets spanning flowers, wood transverse sections, and beetle dorsal habitus (2k–63k images; 7–291 taxa), the Transformer-based encoder transformed each specimen image into a fixed-dimensional “morphological barcode”—here used as a metaphor for an image-derived embedding rather than a homologous sequence marker. Using lineage-specific static or dynamic thresholds together with hierarchical clustering, these embeddings were organized into morphOTUs, producing abundance-based diversity data. In parallel, class activation maps (CAMs) (*42*) highlighted diagnostic morphological regions, including floral symmetry elements, wood vessel architecture, beetle body outlines, sculpturing, and surface textures. These visual explanations allow users to inspect and validate phenotypic differences among morphOTUs, linking cluster boundaries to biologically interpretable traits and accelerating morphological assessment and diversity correction. The three beetle datasets also differed substantially in image acquisition and standardization, including variation in resolution, specimen scale positioning, and background uniformity, allowing us to evaluate morphOTU performance across heterogeneous imaging conditions and specimen-preparation protocols. Together, these results establish a unified, high-throughput workflow for converting digital images into abundance-based morphological OTUs that bridge traditional morphology with modern representation learning (Fig. 1).

**Figure 1.**
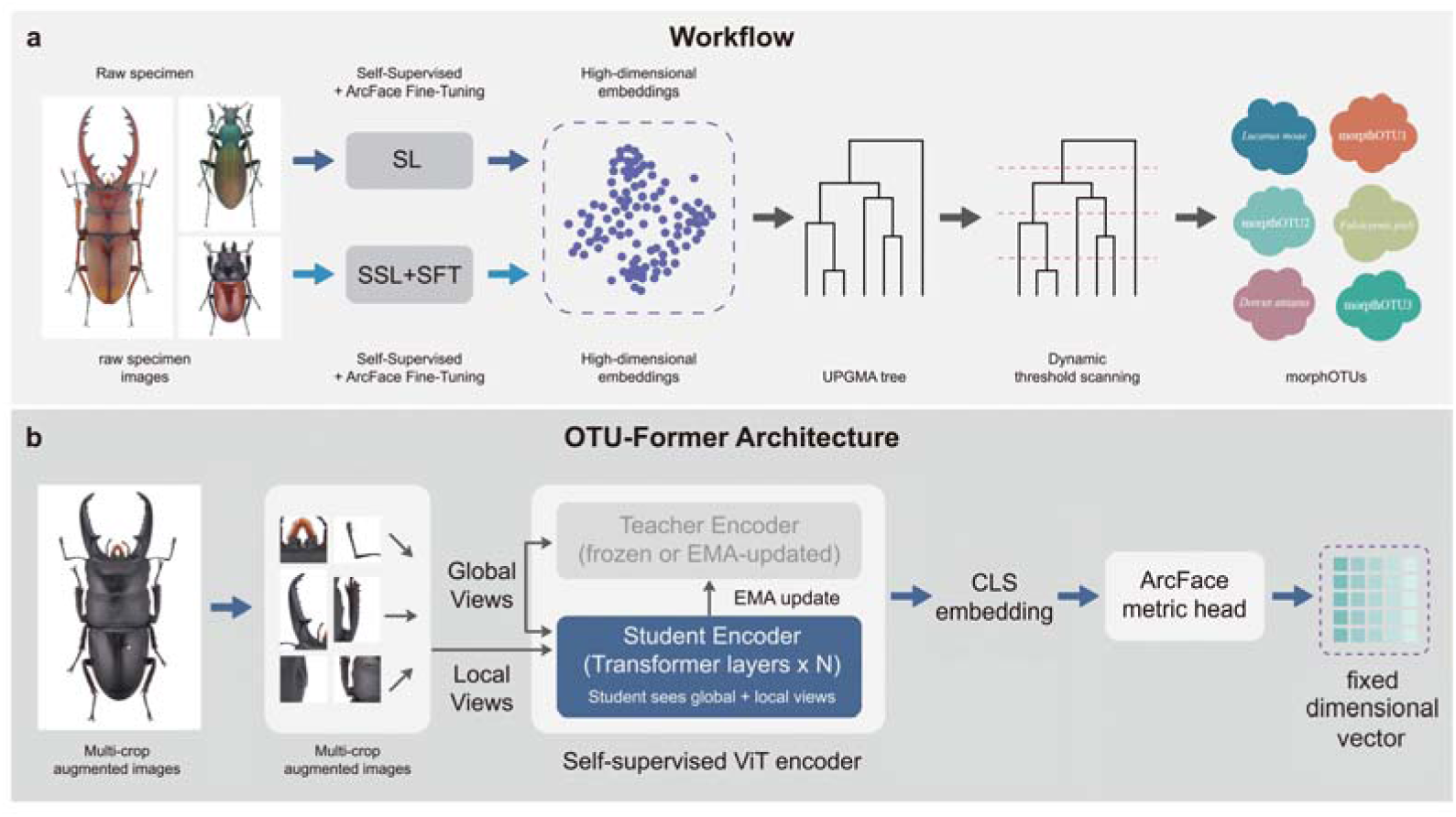
Overview of the morphOTU framework and OTU-Former architecture. **a,** Workflow illustrating the construction of image-based morphological operational taxonomic units (morphOTUs). Raw specimen images are first encoded using two embedding-learning workflows: 1) supervised learning (SL) and 2) the proposed self-supervised plus metric-learning approach (SSL+SFT), to generate high-dimensional morphological embeddings. Hierarchical clustering and dynamic threshold scanning partition the embedding space into stable, abundance-based morphOTUs suitable for biodiversity assessment. **b,** Schematic of the SSL+SFT architecture used in OTU-Former. OTU-Former adopts a stabilized self-supervised teacher–student ViT framework with multi-crop augmented views. The student learns organism-centred morphological features using the teacher—updated via EMA—as a stable self-distillation target.

Global/local contrastive losses and masked-token prediction yield robust CLS embeddings. A subsequent ArcFace fine-tuning stage sharpens class boundaries, producing fixed-dimensional phenotypic vectors for open-set, image-based OTU delineation.

### Benchmarking MorphOTUs on closed-set datasets

Across all five closed-set datasets, we systematically evaluated four embedding-learning paradigms: (1) pretrained backbone models, (2) supervised fine-tuned backbones (SL), (3) self-supervised learning (SSL), and (4) supervised fine-tuning applied to SSL features (SSL+SFT). Our SSL encoder is a lightweight architecture specifically designed for small- to medium-scale datasets (from several hundred to tens of thousands of images), and, when combined with ArcFace fine-tuning (*43*), forms the OTU-Former module (Fig. 1b). To assess model-family differences independent of training strategy, we tested a diverse panel of CNN and Transformer backbones: CNNs (ResNet-50, ConvNeXt-v2, RegNetY) (*44–46*), and Vision Transformers (ViT, EVA02, DINOv3) (*47–49*). Except for OTU-Former, which uses ViT exclusively, we trained each architecture under all four learning paradigms. This design allowed us to compare the relative contributions of backbone choice and training objective to embedding quality.

Across all five closed-set datasets, we evaluated embedding quality using three representative metrics: classification transferability (LP accuracy), retrieval performance (mAP), and clustering quality (ARI). Embeddings extracted from pretrained backbone models already performed well on Flowers-102, likely reflecting the strong overlap between this dataset and the large-scale image corpora used during pretraining. A summary of these performance improvements across the four embedding-learning paradigms is shown in Fig. 2a.

**Figure 2.**
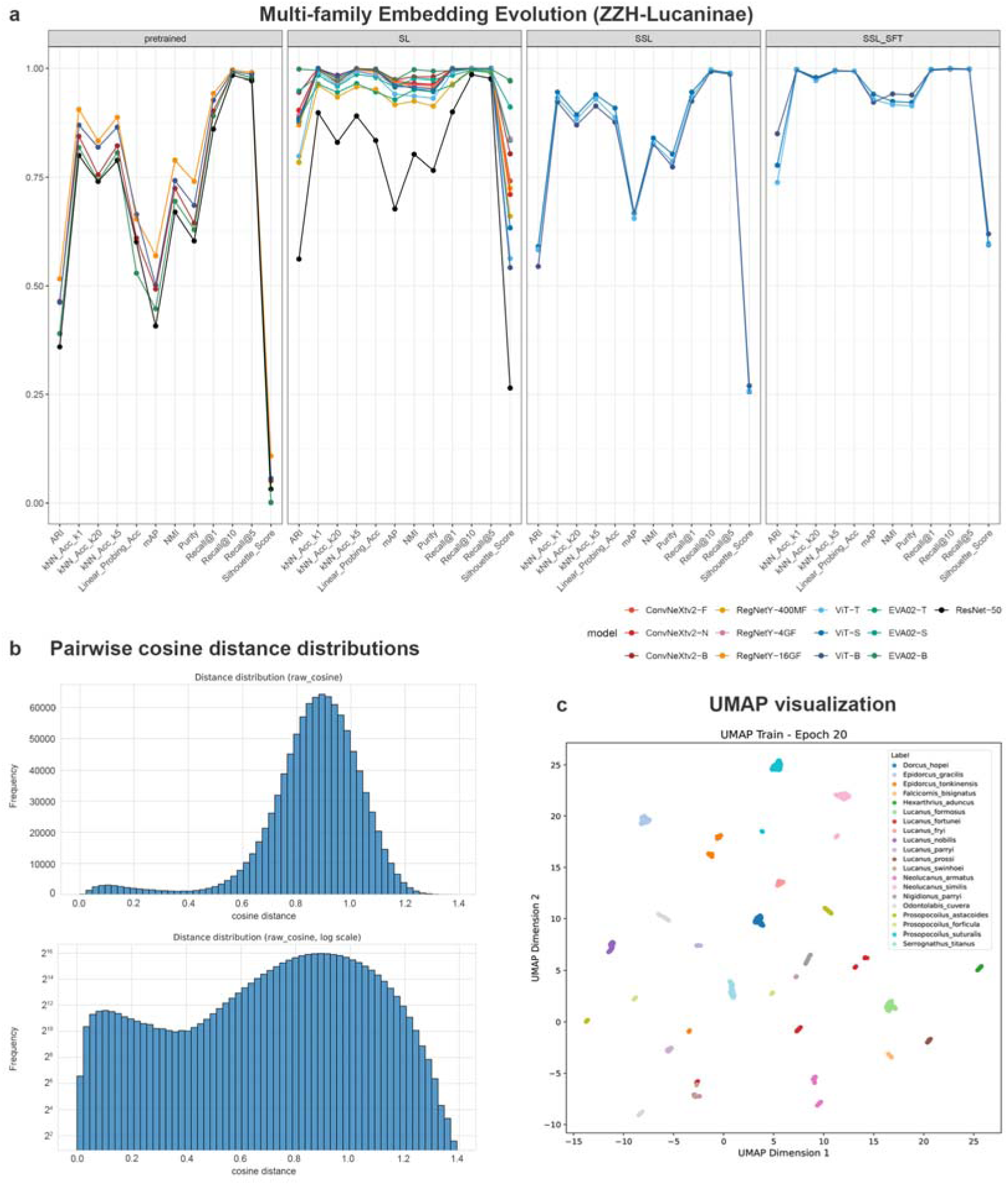
Embedding quality, pairwise distances, and visual clustering under closed-set conditions. **a,** Progressive improvement in embedding structure for ZZH-Lucaninae across four embedding-learning regimes: pretrained backbones, supervised learning (SL), self-supervised learning (SSL), and SSL with supervised fine-tuning (SSL+SFT). **b,** Pairwise cosine distance distributions from SSL+SFT embeddings for ZZH-Lucaninae. The broad, partially structured distribution reflects overlapping intra- and interspecific variation rather than a discrete barcoding gap. **c,** UMAP visualization of SSL+SFT embeddings for ZZH-Lucaninae after supervised fine-tuning shows well-defined species-level structure. Together, these panels demonstrate that morphOTU generates biologically coherent phenotypic organization under ideal closed-set conditions.

For Flowers-102, further fine-tuning offered limited additional benefit in this case, and we therefore did not conduct SSL experiments. In contrast, embeddings from WOOD and ZZH-Lucaninae were moderate in quality, whereas Rove-Tree-11 and NHM-Carabids showed substantially lower performance (e.g., LP accuracy < 0.5, mAP < 0.3, ARI < 0.35; Fig. S1). As expected, ResNet-50 consistently lagged behind the other backbones, while the remaining pretrained CNN and Transformer models showed no decisive winner.

Supervised fine-tuning using AutoGluon substantially improved embedding quality across nearly all backbones and datasets. Except for RegNetY-400MF, all models achieved LP accuracy > 0.95, mAP > 0.85, and ARI > 0.7 across the five datasets (Fig. S2). ConvNeXt-v2 and EVA02 exhibited the strongest performance overall; remarkably, even their smallest variants (∼5M parameters; ConvNeXt-v2-F and EVA02-T) maintained most metrics above 0.95.

Self-supervised learning (SSL) provided a notable improvement over pretrained backbones, with LP accuracy increases exceeding 0.2 on several datasets and approaching 1.0 on WOOD. As dataset difficulty increased, the relative gains achieved by SSL diminished, indicating that datasets of varying complexity offer different capacities for effective feature learning (Fig. S3). After ArcFace fine-tuning (SSL+SFT), embedding quality approached that of fully supervised training across all datasets (Fig. S4), including the most challenging NHM-Carabids dataset, where LP accuracy exceeded 0.97, mAP surpassed 0.9, and ARI exceeded 0.7. Within the two-stage OTU-Former pipeline, increasing ViT model size (T/S/B) yielded limited gains, indicating that moderate-capacity encoders are sufficient for small- to medium-scale biodiversity datasets. Together, these embedding evaluations show that supervised fine-tuning (SL, SSL+SFT) and SSL alone produce high-quality, clusterable morphological barcodes under closed-set conditions.

To characterize embedding geometry, we computed pairwise cosine distances between all images. Distance ranges were largely determined by model family: RegNetY and ResNet-50 produced narrower ranges (0–1), substantially lower than other models (0–1.5; Fig. S5). SSL distances were generally slightly smaller than SL or SSL+SFT (Fig. S6), likely because supervised fine-tuning explicitly enlarges interclass margins (*50*, *51*). Unlike molecular genetic distances, which often display a well-defined “barcoding gap”, cosine-distance distributions from image embeddings exhibited no single, discrete separation (Fig. 2b; Figs. S7–S16). Instead, they showed at most a shallow trough, consistent with lineage-specific variability in morphological divergence. UMAP projections of the same embeddings nevertheless showed well-defined species-level structure under SSL+SFT (Fig. 2c). Intra-class distances remained tightly constrained, with mean values typically within approximately 0.3 (Fig. S17). These patterns jointly indicate that although fixed global thresholds are inappropriate, morphology-based OTU formation remains feasible when adaptive, lineage-specific thresholds are used.

To examine this process, we applied UPGMA trees to illustrate how dynamic thresholds determine morphOTU boundaries. As shown in Fig. 3a, no single cutoff yields perfect partitions: some thresholds oversplit species (P1–3), whereas others lump several species into one OTU (P2–4). Ideally, final decisions would incorporate expert evaluation or CAM-based guidance, using multiple partitioning schemes and UPGMA trees (with jackknife support) to verify ambiguous groupings. To streamline this procedure and assess practicality, we compared morphOTUs generated under a single (often suboptimal) threshold against reference species labels. Using a two-stage dynamic threshold scanning procedure, we identified lineage-appropriate interspecific distance ranges (typically spanning 0.2–0.4), selected the best-scoring cutoff according to BCubed-F score (*52*), and calculated richness and Shannon diversity indices. The BCubed-F optimum was used only for retrospective benchmarking against reference species labels. In label-free applications, candidate thresholds would instead be selected using distance-profile stability, jackknife-supported clusters, abundance filtering, and expert inspection of CAMs. MorphOTU-derived diversity estimates closely matched true species-level values, particularly when abundance-based noise filtering was applied. Shannon indices were less sensitive than richness, but all three metrics showed high concordance with reference species labels (Fig. S18).

**Figure 3.**
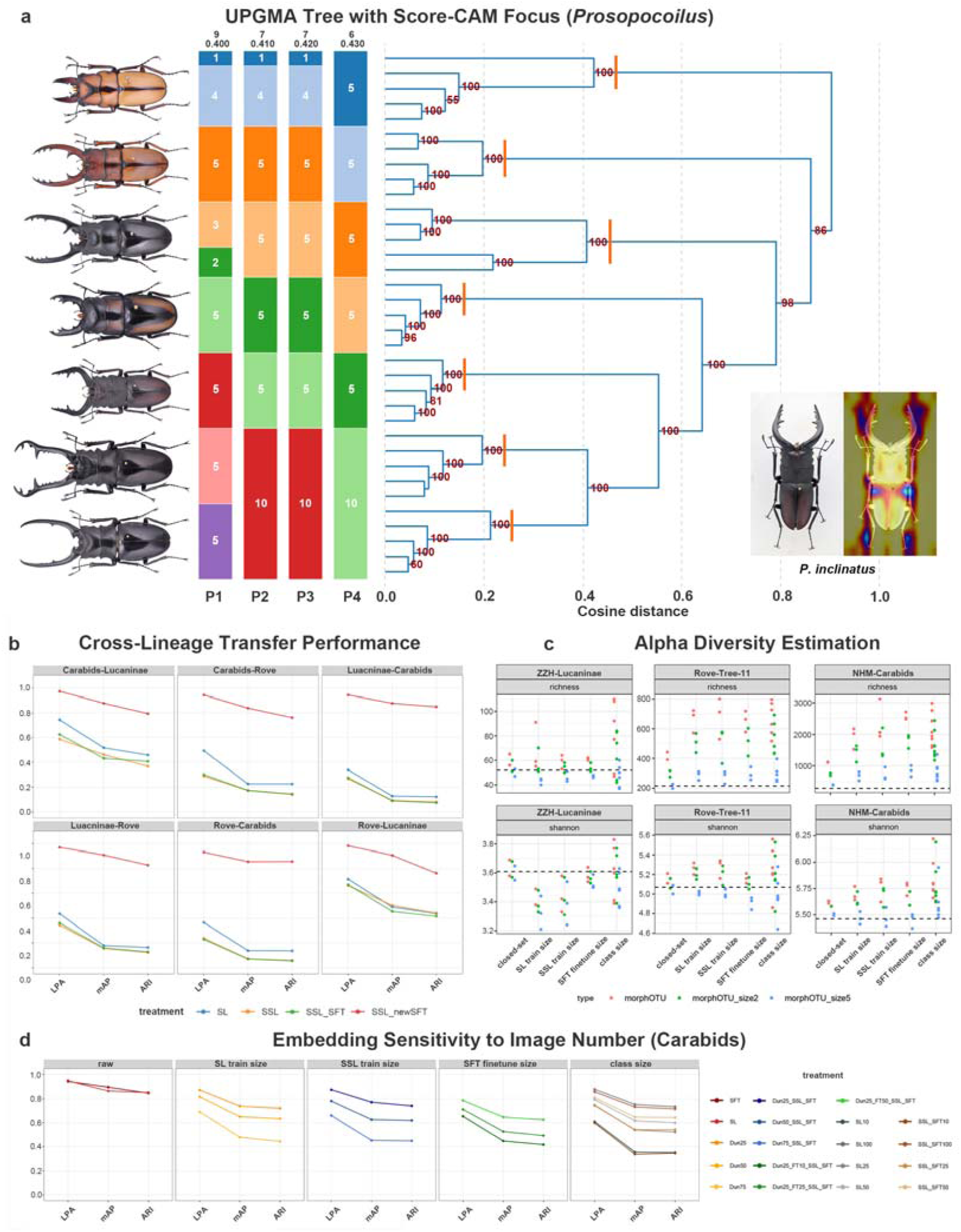
Dynamic thresholding, interpretability, cross-lineage generalization, and sampling robustness. **a,** Dynamic threshold scanning on a UPGMA tree for *Prosopocoilus*. Stable morphOTU boundaries are revealed across cosine-distance cutoffs. Orange vertical bars indicate reference species labels, and the four partitioning schemes (P1–P4) illustrate how clusters split or merge as thresholds increase. Score-CAM overlays (right) highlight the diagnostic morphological regions driving these separations, linking phenotypic evidence directly to tree structure. **b,** Cross-lineage generalization across beetle datasets. Models trained on one lineage show reduced performance—Linear Probe Accuracy (LPA), mAP, and ARI—when transferred to another due to deep morphological divergence. Fine-tuning the SSL encoder on labeled data from the receiving lineage (SSL_newSFT) restores cluster separability and retrieval performance, demonstrating the portability of OTU-Former features across lineages. **c,** α-diversity estimation under real-world constraints. Richness and Shannon diversity remain stable across closed-set and progressively more challenging open-set scenarios. SSL+SFT maintains consistent diversity estimates, particularly after low-abundance morphOTU filtering, underscoring the robustness of morphOTUs under open-set noise and sparse taxonomic labeling. **d,** Sensitivity of embedding structure to per-species image number (Carabids). Panels compare how different data-limitation regimes affect embedding separability (LPA, mAP, ARI). Five training-data conditions are shown: raw: Full SSL+SFT trained on all images from closed-set species; SL train size / SSL train size / SFT finetune size: Three subsampling regimes in which each species is limited to 100, 50, 25, or 10 images, applied respectively to supervised training, SSL pretraining, or ArcFace fine-tuning (SSL pretraining uses full unlabeled data unless subsampling is explicitly applied); class size: Closed-set species filtered by a minimum-image threshold (species ≥105), altering training-set composition while retaining each species’ full sample size.

Overall, these results demonstrate that morphological barcodes produced by both SL and SSL-based training regimes are stable and intrinsically clusterable. When paired with dynamic, lineage-specific distance thresholds—and optionally guided by expert inspection—the resulting morphOTUs approximate reference species-level groupings with high fidelity. Even when using a single operational cutoff within each dataset, morphOTU-based diversity indices remain usable and closely match the expert-assigned species labels, confirming the feasibility of morphology-driven OTU construction on closed-set datasets.

### Robustness under real-world biodiversity constraints

#### Tests for case scenarios under realistic limitations

In real-world applications, available training data, particularly specimens that can be confidently identified to species (i.e., labeled data), are often limited due to well-known scaling constraints, and operational settings frequently involve open-set conditions. To further estimate the practical scope, strengths, and weaknesses of the morphOTU framework under such realistic constraints, we designed a series of subsampling experiments across the three beetle datasets. All tests employed a unified backbone architecture (ViT-T) to control for model capacity. Evaluation criteria remained consistent with the closed-set benchmarks, including embedding quality, variation in pairwise distance distributions, and the accuracy of diversity estimates.

#### Open-set with “unseen” species

Real-world biodiversity surveys frequently involve open-set conditions in which many encountered species are absent from the training data: a scenario especially common in invertebrate groups (*10*). To evaluate how morphOTU behaves under such conditions, we examined the generalization performance of both SL and SSL+SFT embeddings as the number of open-set species increased, thereby characterizing the framework’s scaling behavior. We constructed four gradients in which unseen species constituted 0%, 25%, 50%, and 75% of all images. SL and SSL+SFT performed comparably across these settings.

As expected, embedding quality decreased as the proportion of open-set data increased. When unseen species increased to 75% of all images, leaving only 25% represented in training, LP accuracy dropped to 0.66–0.81, mAP to 0.45–0.56, and ARI to 0.44–0.46 (Figs. S19, S20). Pairwise cosine distances and intra-class distances also became slightly more variable as unseen species increased (Figs. S21, S22), indicating a loosening of embedding compactness. Nevertheless, the overall embedding geometry remained structured, suggesting that meaningful clusters can still be recovered even when the majority of species are not present during training.

When morphOTUs were generated using a single distance threshold, OTU counts increased relative to baseline models trained under closed-set conditions. After abundance filtering, ZZH-Lucaninae and Rove-Tree-11 retained richness estimates similar to their closed-set baselines, whereas NHM-Carabids exhibited more than a two-fold increase in estimated OTU richness. Shannon diversity derived from morphOTUs showed errors within 0.4, with abundance-filtered estimates performing better in larger datasets (Fig. 3c; Fig. S18).

Our experiments demonstrate that scaling laws strongly influence embedding quality, yet morphOTUs retain coherent structure even when the available training data are reduced to only 25% of the full dataset. This robustness is crucial for practical applications such as discovering novel species and quantifying diversity in large volumes of unidentified material under realistic open-set ecological conditions.

#### Sparse labeled supervision

In the scaling-law tests above, labeled data and training data were identical. However, in practical self-supervised learning workflows, the amount of labeled material is typically far smaller than the amount of imagery available for training. This corresponds to a common biodiversity scenario in which large numbers of specimens can be incorporated into pretraining (without labels), but only a limited fraction can be taxonomically identified. To assess the impact of labeled data availability on the ArcFace SFT stage, we varied the size of the annotated subset while keeping SSL pretraining fixed. Experiments were conducted on the Dun25 setting, with 75% of images used for SSL pretraining and the remaining 25% unseen species held out as open-set test data. We then fine-tuned ArcFace using 75% (as in the pretraining), 50%, 25%, or 10% of the labeled portion.

Even with only 10% labeled data for SFT, embeddings remained stable, achieving LP accuracy > 0.65, mAP > 0.44, and ARI > 0.41. Pairwise cosine distances and intra-class distances also remained consistent across label fractions (Figs. S19, S20). Diversity estimates behaved similarly to the open-set results above: richness and Shannon indices derived from morphOTUs remained close to expert-assigned species labels (Fig. S18).

These results show that, given strong SSL pretraining, extensive high-quality taxonomic annotation is not essential. Moderate labeled fractions (10%–25% in this study) are sufficient to preserve good generalization performance, in line with contemporary paradigms of “large-scale pretraining followed by small-scale finetuning” (*53*, *54*). Open-set data can be fully leveraged during SSL pretraining to further enhance representation quality. In the most extreme cases, such as WOOD and ZZH-Lucaninae, SSL pretraining alone, without any supervised finetuning, was sufficient to recover reliable morphOTUs (Fig. S3).

Across these regimes, SSL embeddings remain more robust than SL, and SSL+SFT achieves the highest stability. Performance declines sharply only when species have ≤10 images, emphasizing the minimum sampling required for reliable morphology-based OTU construction.

#### Limited images per species

We next examined how reductions in per-species sample size affect generalization by reducing the observed intra-class diversity. Using fixed training and test species, we created four sampling regimes with 100 images per species (NHM-Carabids only), 50, 25, and 10 images per species. Test sets always included the remaining closed-set material as well as open-set species. Embedding quality declined with decreasing sample size, but performance remained acceptable at ≥25 images per species (LP accuracy > 0.75, mAP > 0.52, ARI > 0.49), and cosine distance and intra-class distance distributions remained stable. At 10 images per species, however, performance degraded sharply (Fig. 3d; Figs. S19−S22).

SSL encoders were consistently more robust than supervised models for ZZH-Lucaninae and Rove-Tree-11, whereas performance on NHM-Carabids was weaker, potentially owing to its greater variation in image resolution, specimen scale, and background uniformity, which may hinder the learning of generalizable features. Importantly, limited labeled sampling per species does not necessarily constrain the scale of SSL pretraining, because large external collections of unlabeled biodiversity images could be incorporated to improve representation quality without additional species-level annotations. Richness and Shannon indices remained accurate after abundance filtering, although errors at n = 10 per species exceeded those from all prior testing scenarios (Fig. S18).

#### Cross-lineage transfer

We also evaluated an extreme setting in which a model trained on one beetle dataset was tested entirely on the other two (i.e., test sets consisted exclusively of open-set species). Embedding clusterability improved with larger training sample sizes, confirming that representation learning remains sensitive to sample density. SL embeddings performed slightly better than SSL and SSL+SFT in this fully open-set setting, but overall performance remained limited (Fig. 3b). After ArcFace fine-tuning on the receiving lineage (SSL_newSFT), however, both embedding quality (LP accuracy > 0.93; mAP > 0.83; ARI > 0.76) and diversity estimates (Fig. S18) reached levels comparable to the closed-set baselines.

#### Accelerating morphological diagnosis through interpretable feature representations

Beyond providing diversity estimates, morphOTUs derived from image embeddings also enable transparent visualization of the morphological cues underlying OTU separation through class-activation mapping (*42*). If the model successfully learns object-focused rather than background-driven features, the resulting heatmaps should reveal meaningful morphological differences among individuals and species. To evaluate this, we constructed three genus-level subsets for each beetle dataset (Table 2), with sample sizes ranging 230−13,265 images, and examined whether SL and OTU-Former architectures consistently attend to the beetle body and whether the highlighted regions correspond to biologically interpretable traits. Here, we define “biologically interpretable” as regions that: (i) correspond to diagnostic characters used in taxonomic keys or formal species descriptions (e.g., body outline, mandible shape, elytral sculpture, punctation patterns), and (ii) were independently confirmed by expert entomologists as taxonomically meaningful.

These criteria allow us to evaluate interpretability using both formal morphological character systems and expert assessment. Consequently, OTU-Former completed its full pretraining pipeline on all subsets, and its two-stage embedding metrics remained comparable to the closed-set benchmarks, indicating that the framework trains reliably even on very small (around 200 images) datasets.

Our datasets contain relatively simple backgrounds, and both SL and OTU-Former models generally focus on the beetle body rather than clustering on spurious background cues (Fig. 4a). Nonetheless, OTU-Former consistently produced cleaner and more object-focused Score-CAM heatmaps (*55*), with substantially larger attention concentrated on the beetle body compared to supervised baselines (shown as paired SL vs SFT examples for each species). This indicates that OTU-Former learns morphological features more effectively and is less susceptible to background artifacts or label-related noise. Expert evaluation of these Score-CAM maps further showed that OTU-Former preferentially attends to biologically meaningful structures, such as body outlines, cuticular texture, and other continuous phenotypic traits that are difficult to quantify using traditional morphological character systems. In this sense, morphOTUs function as “OTUs in continuous phenotype space,” complementing discrete, character-based taxonomy and providing an interpretable and image-traceable representation of morphological variation.

**Figure 4.**
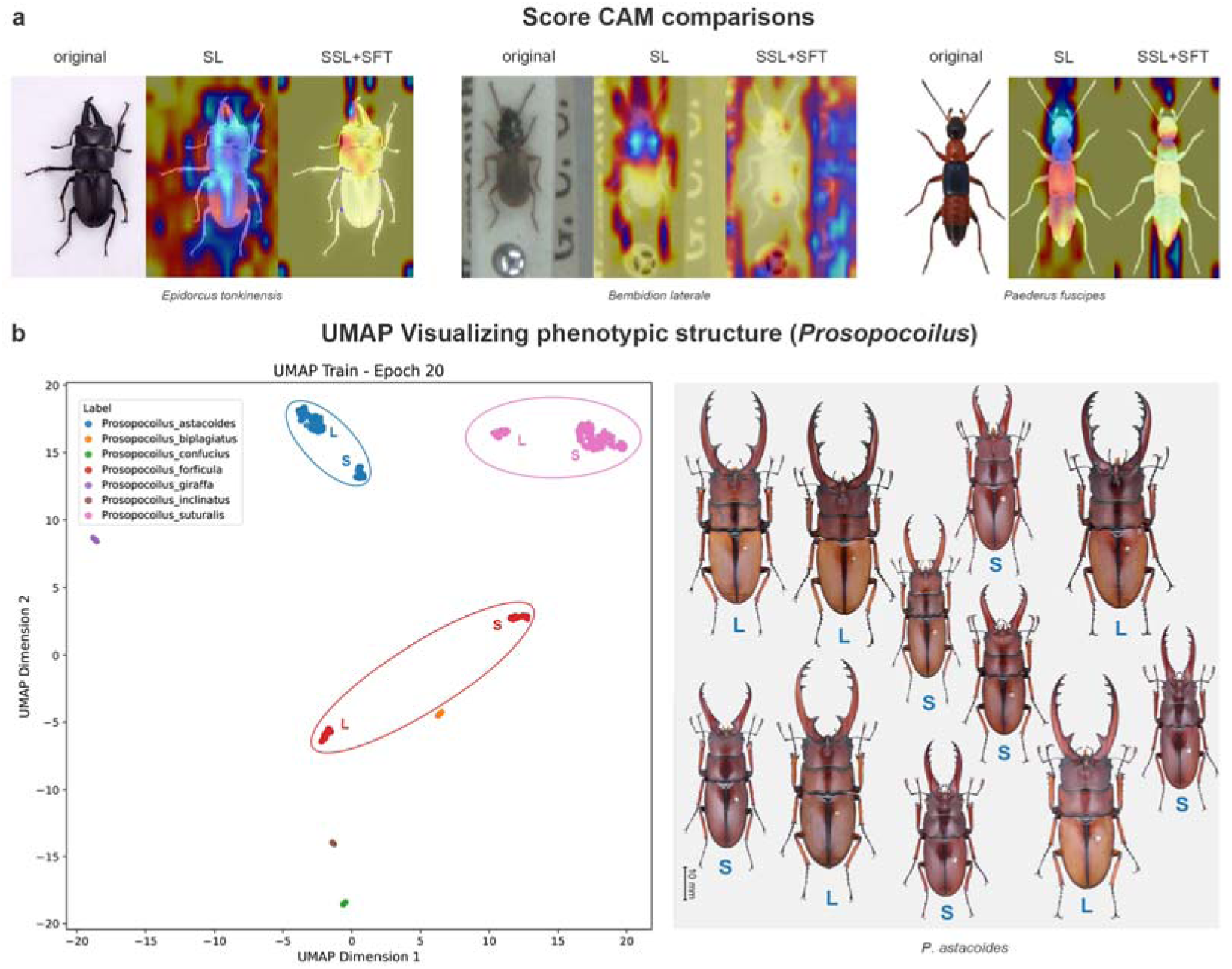
Morphological interpretability and taxonomic insight from morphOTUs. **a,** Score-CAM comparisons between supervised learning (SL) and SSL+SFT across beetle genera. OTU-Former focuses more precisely on biologically informative structures, yielding cleaner and more coherent activation maps that align with expert-known diagnostic traits. **b,** Fine-scale phenotypic structure in *Prosopocoilus* revealed by SSL+SFT embeddings. Left: UMAP visualization showing clear separation of large-bodied (L) and small-bodied (S) individuals across different species, highlighting subtle intraspecific morphological structuring captured by morphOTUs. Right: Representative *P. astacoides* specimens (equal scale; scale bar = 10 mm) illustrating the corresponding morphological differences between large-bodied (L) and small-bodied (S) individuals. These examples demonstrate how image-based morphOTUs capture continuous phenotypic gradients not evident from coarse categorical traits alone.

The nine genus-level subsets represent groups of closely related species that are often challenging in practical morphology-based classification (Fig. S28). Detailed examination of morphOTUs within the three ZZH-Lucaninae genera revealed that OTU-Former can resolve fine-scale phenotypic differences, including distinct body-form variants within the same species (Fig. 4b), as well as clearer separations among historically problematic taxa. These results demonstrate that morphOTUs provide a consistent and interpretable representation of fine-scale morphological structure, supporting rapid diagnosis, character exploration, and expert knowledge transfer within closely related species groups.

#### Impact of low-quality images on training performance

To evaluate how image quality influences representation learning, we conducted a controlled degradation experiment on the high-quality ZZH-Lucaninae dataset. Images were downsampled to short-edge resolutions of 512, 256, 128, 64, and 32 pixels, thereby simulating increasing levels of quality loss (Fig. S27). All models (SL, SSL, and SSL+SFT) were trained exclusively on degraded images, and embedding quality was evaluated on the original 512-pixel test images. This design evaluates the effect of low-quality training data under a standardized high-quality evaluation set. It does not isolate the performance expected when both training and test images are degraded.

Supervised learning (SL) and supervised fine-tuning after self-supervised pretraining (SSL+SFT) exhibited minimal performance loss at 512, 256, and 128 pixels, but showed pronounced declines at 64 and 32 pixels (Fig. 5a). In contrast, purely self-supervised models (SSL) began to degrade substantially at resolutions below 128 pixels, indicating greater sensitivity to image degradation that disrupts semantic consistency and induces distributional shift (*56*, *57*). This sensitivity also partly explains the weaker SSL performance observed on the NHM-Carabids dataset, where beetles often occupy only a small portion of the frame and many specimens are blurred or lack clear structural details.

**Figure 5.**
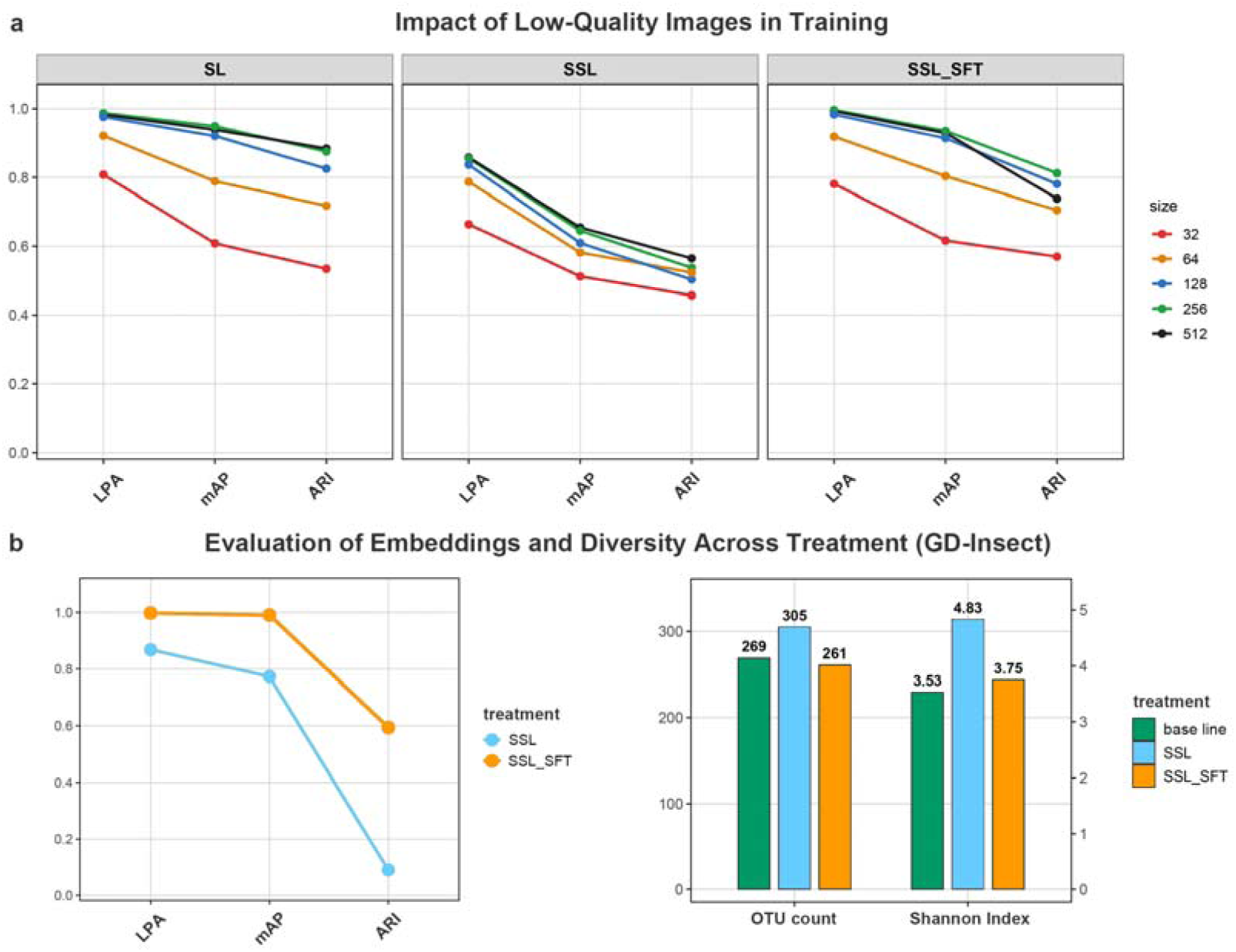
Image quality effects and real-world performance of morphOTUs. **a, Sensitivity of representation learning workflows to controlled image degradation.** Downsampling experiments on the high-quality ZZH-Lucaninae dataset reveal distinct robustness profiles across training strategies. Supervised learning (SL) and SSL+SFT maintain stable embedding quality at 512–128 px, but both decline sharply at 64–32 px. In contrast, purely self-supervised models (SSL) begin degrading below 128 px, reflecting greater vulnerability to loss of structural detail and semantic consistency. These trends explain the weaker SSL performance observed in the NHM-Carabids dataset, where beetles often occupy only a small portion of the frame and many specimens are blurred or low-detail. **b, Real-world performance of morphOTUs on the GD-Insect biodiversity survey dataset.** OTU-Former was evaluated on 4,717 habitus images spanning 269 species across 12 insect orders, with extreme class imbalance and heterogeneous imaging conditions. Fully self-supervised embeddings (SSL) produced diffuse morphological structure and inflated OTU counts, reflecting the difficulty of clustering highly diverse assemblages without labeled guidance. Fine-tuning the SSL encoder using ArcFace metric learning on only 28 common species represented by at least 30 individuals substantially improved embedding quality and morphOTU stability, yielding OTU partitions and Shannon diversity estimates closely aligned with reference species labels. These results demonstrate that light expert supervision can substantially enhance morphOTU accuracy under open-set, imbalanced, and taxonomically complex real-world conditions.

These results highlight a critical practical insight: while moderate image degradation has a limited impact on supervised or semi-supervised workflows, self-supervised feature learning is considerably more fragile to low-resolution or low-detail inputs. Consistent image quality and sufficient organismal visibility, therefore, remain essential for maximizing the benefits of self-supervised representation learning in large-scale biodiversity applications.

#### Performance of OTU-Former on a real-world dataset

To assess the applicability of morphOTUs under realistic field conditions, we evaluated OTU-Former on the GD-Insect dataset, which contains 4,717 habitus images representing 269 species across 12 insect orders. Embedding metrics were computed on the subset of species with sufficient replication for metric evaluation, whereas all 269 species, including singletons, were retained for morphOTU richness and Shannon-diversity estimation. The dataset exhibits extreme class imbalance, as 152 species are represented by a single individual, and includes both terrestrial and aquatic taxa (Shannon index = 3.53).

After self-supervised pretraining using all available images, SSL embeddings achieved LP accuracy = 0.8675, mAP = 0.7731, and ARI = 0.0906. Using a single distance threshold, morphOTU partitioning produced an overestimated 305 OTUs and a Shannon index of 4.83. These results reflect the difficulty of learning robust morphological structure from highly imbalanced and taxonomically heterogeneous data without any labeled supervision (Fig. 5b).

To test whether limited expert annotations can improve performance, we fine-tuned the SSL encoder using ArcFace metric learning applied only to 28 common species represented by at least 30 individuals. This modest supervision substantially improved embedding quality: LP accuracy increased to 0.9977, mAP to 0.9902, and ARI to 0.5940. Correspondingly, both morphOTUs (261 units) and the estimated Shannon index (3.75) better approximated the expert-assigned species labels, which refer to expert-assigned species labels used only for retrospective evaluation.

These results demonstrate that fully unsupervised pretraining combined with light supervision from a small number of common, easily diagnosed species offers a practical and efficient strategy for generating reliable diversity estimates in real-world insect assemblages. With even minimal expert correction, the resulting morphOTUs can provide ecologically interpretable and high-fidelity diversity data under open-set, imbalanced, and taxonomically complex field conditions.

## Discussion

### A new generalizable workflow for image-based biodiversity discovery

Our morphology-based OTU workflow represents more than a technical improvement in species identification: it signals a broader conceptual shift toward scalable, data-driven phenotyping in biodiversity research. By integrating self-supervised feature learning with lineage-specific clustering, we can convert raw images into standardized, abundance-based morphOTUs, reducing reliance on expert-driven labeling and enabling biodiversity assessment at scales previously unattainable via morphology. This transition parallels the transformative impact that DNA barcoding and metabarcoding had on molecular ecology two decades ago.

Beyond these operational advantages, morphOTU helps mitigate the long-standing taxonomic impediment by reducing dependence on complete expert identification and enabling preliminary organization of poorly annotated specimens (*8*, *9*, *11*). Although performance still benefits from sufficient organismal visibility and image quality, the framework can operate on moderately heterogeneous image collections. Different from many previous deep-learning approaches, which depend on closed-set supervised classification and therefore are difficult to apply under biodiversity conditions with scarce labels (*33*, *34*, *39*), morphOTU uses self-supervised learning to extract organism-centered features while reducing reliance on background artifacts (*35*, *36*). This enables scalable, open-set phenotyping even when most species remain unidentified. This claim is further supported by our real-world evaluation on the GD-Insect dataset. During supervised fine-tuning, only 28 species (10.41%) were labeled and included in training, while the remaining approximately 90% of species were entirely unseen. The model constructed morphOTUs from a mixed test set containing both seen and unseen taxa, demonstrating stable clustering and consistent phenotypic grouping under realistic, partially labeled open-set conditions.

Unlike classical morphology-based workflows, morphOTU can operate on moderately heterogeneous image collections, but its performance benefits from sufficient organismal visibility, resolution, and standardized imaging. Because morphOTUs can be generated with limited expert labels, the framework may reduce the amount of taxonomic training required for preliminary sorting and diversity assessment. The ability to visualize diagnostic structures through activation maps is particularly significant: continuous morphological traits, which are foundational but challenging to formalize, become directly observable and computationally accessible. Previous deep-learning approaches have been constrained by closed-set assumptions and limited interpretability; in contrast, morphOTU learns organism-centered features in an open-set framework, achieving supervised-level performance while maintaining robustness in systems with many undescribed taxa.

Conceptually, morphOTUs function as morphological analogues to molecular OTUs and metabarcoding-derived units (*19*, *20*). They provide a non-destructive, infrastructure-light representation of phenotypic space without requiring sequencing infrastructure or per-sample sequencing costs (*58*). Rather than replacing molecular data, morphOTUs complement and extend existing taxonomic practice: they offer rapid preliminary sorting, reveal phenotypic structure invisible to coarse categorical traits, and flag specimens that merit targeted sequencing. This integrative workflow aligns morphology, molecules, and machine learning into a coherent system for biodiversity discovery, supporting taxonomic revision and enhancing the digitization and curation of natural history collections.

### No universal distance threshold delineates morphOTUs across all lineages

Although conceptually parallel to molecular OTUs, morphOTUs differ fundamentally in their statistical properties. Molecular systems often exhibit a broad barcoding gap separating intra- and interspecific divergence (*59*, *60*), typically around a COI threshold of 0.03 (*19*), although overlap between classes may occur (*61*, *62*). Contrary to this, morphological distances widely vary within species and show substantial overlap across species. In our datasets, intra-class morphological variation often spanned cosine-distance ranges up to 0.3, illustrating that image-embedding distances do not exhibit the same threshold behavior as COI sequence divergences, reflecting the evolutionary nature of phenotypic variation (e.g., Fig. S7). Some lineages diversify through discrete, readily visible traits, while others exhibit continuous variation shaped by adaptation, sexual selection, or phenotypic plasticity. Consequently, no universal morphological threshold can reliably delineate species-level units, and morphological barcodes do not exhibit clean “gaps” (Fig. 2b).

Our dynamic-thresholding strategy accommodates this heterogeneity by generating multiple partitioning schemes, enabling stable clusters to be identified across thresholds and corroborated using Score-CAM visualizations or taxonomic knowledge (Fig. 3a). Although absolute OTU boundaries vary with threshold choice, community-level diversity metrics derived from single-threshold morphOTUs were robust, suggesting that flexible thresholding need not impede ecological inference.

Imaging conditions can also influence MorphOTU counts. Variations in background, orientation, illumination, or effective resolution can produce long branches or weakly supported nodes in UPGMA trees, highlighting individuals requiring expert review or molecular validation. The NHM-Carabids dataset (*33*) underscores this: despite its size, lower image quality reduced embedding clarity and increased uncertainty, consistent with the reliance of self-supervised representation learning on high-quality, standardized data (*63*, *64*) (Figs. S23−S27). These findings reinforce the value of standardized imaging, particularly when leveraging self-supervised representation learning, which benefits strongly from high-quality data. Although standardized imaging improves overall consistency, our degradation experiment demonstrates that the primary limitation for self-supervised representation learning is image quality, particularly organismal visibility and structural detail, rather than standardization alone. This distinction explains why the NHM-Carabids dataset, which contains many low-detail, small-scale specimens, yields weaker representations despite its relatively uniform imaging protocol.

These observations argue for a conceptual shift away from fixed-threshold morphology and toward multi-resolution, expert-informed frameworks. Such an approach is more faithful to the evolutionary complexity of phenotypes and more suited to the diversity of organisms encountered in large-scale biodiversity surveys.

### Limitations and prospects

Although morphOTU demonstrates broad applicability, several limitations point to opportunities for future development. As with molecular OTUs, morphological heterogeneity across lineages makes it difficult for a single external marker, for example, the dorsal view, to capture the full range of diagnostic traits. Integrating multiple views, combining several morphological markers, or leveraging emerging 3D and multiview technologies could significantly enhance morphOTU resolution and improve generality across taxa. The taxonomic coverage and imaging diversity of our current datasets remain limited, underscoring the need for broader evaluation across additional lineages and imaging contexts. Moreover, applying morphOTU to holometabolous insects, where larval and adult stages exhibit drastically different morphologies, remains challenging and requires future methodological extensions. Additionally, substantial intraspecific variation arising from sexual dimorphism, ontogeny, allometry, geographic differentiation, and phenotypic plasticity may cause individuals of the same species to be partitioned into multiple morphOTUs. Conversely, morphologically similar or cryptic species may be merged into a single unit. Broader sampling across sexes, life stages, populations, and environmental conditions will therefore be important for improving morphOTU stability and biological interpretability.

From a technological perspective, morphOTU was optimized for medium-sized datasets and accessible hardware. As visual foundation models continue to evolve, integrating larger, more generalizable encoders trained on global biodiversity imagery will be essential for building a truly universal morphological representation. Expanding training to include diverse environments, imaging styles, and taxonomic groups will further improve robustness and reduce domain-specific biases.

Looking ahead, image-based morphOTUs have the potential to reshape biodiversity science. They could enable real-time phenotypic monitoring in ecological observatories, support global re-examination of museum collections at an unprecedented scale, and provide early-warning systems for invasive species or emerging pests. MorphOTUs and molecular OTUs, when integrated, offer a pathway toward a unified, global framework for biodiversity assessment encompassing both phenotypic and genetic dimensions of organismal diversity across ecosystems, time periods, and spatial scales.

## Materials & Methods

### Datasets and images preparation

#### Dataset selection

To operationalize the concept of morphOTUs, image datasets must represent a standardized and comparable morphological trait within each organismal group—analogous to using a single universal marker such as COI in molecular barcoding. Most publicly available image collections include heterogeneous traits, multiple anatomical views, inconsistent angles, or variable backgrounds, making them unsuitable for constructing coherent morphological embeddings. We therefore assembled two plant datasets and four insect datasets (Table 1), each containing images captured from a single, biologically interpretable trait: floral structures, wood transverse-section anatomy, or the dorsal or lateral habitus of adult insects. After image cleaning, dataset sizes ranged from 2,368 to 63,053 images and from 7 to 291 labeled categories, reflecting the typical scale of small- to medium-sized biodiversity datasets. We used five standardized datasets for benchmarking and one additional real-world dataset for validation. Except for WOOD, all datasets exhibit characteristic long-tailed class distributions, mirroring the highly uneven species-abundance patterns that dominate real biodiversity assemblages (Fig. 6).

**Figure 6.**
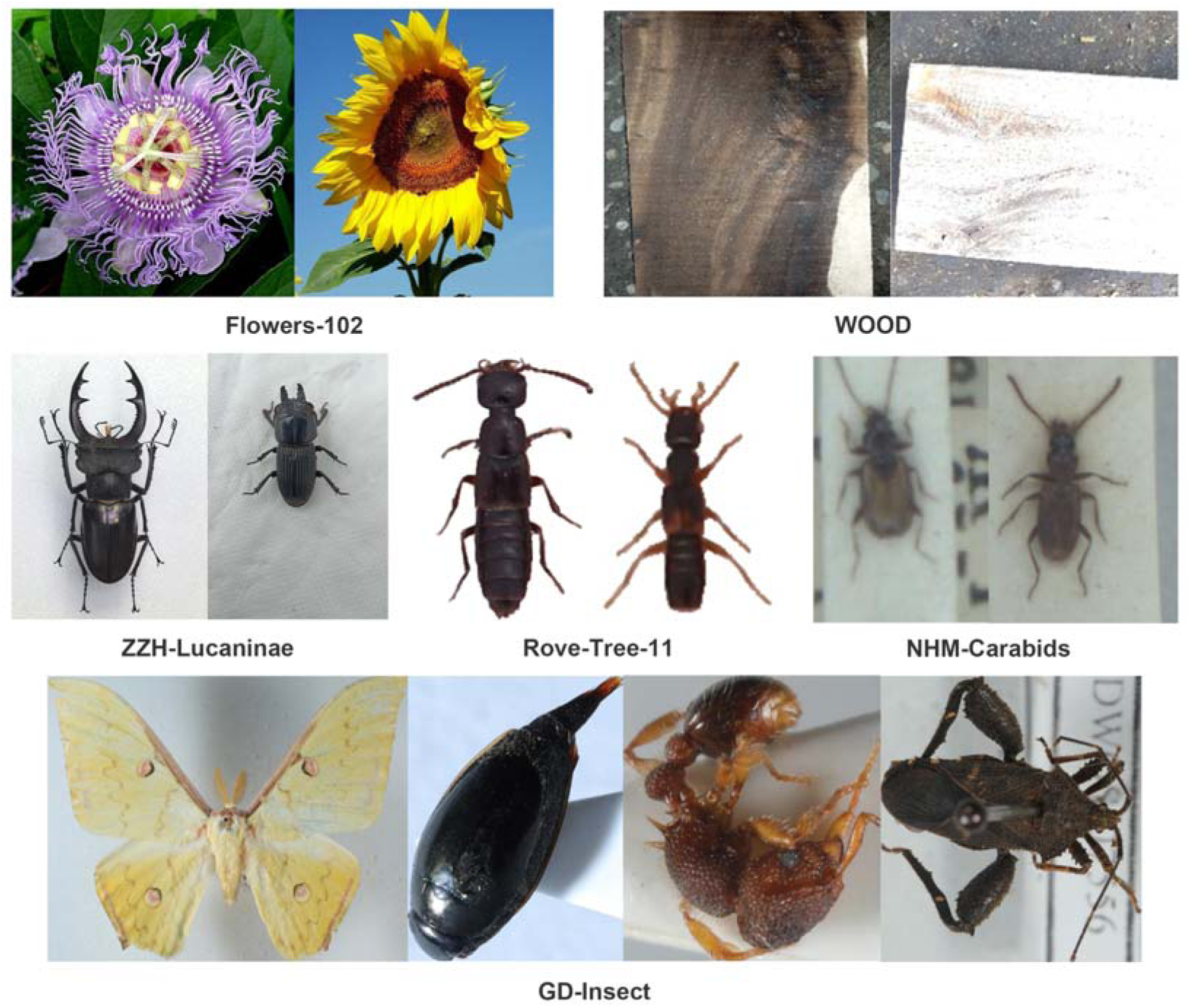
Representative datasets and morphological interpretability enabled by morphOTUs. Overview of the six datasets used in this study: Flowers-102 (*65*), WOOD (*66*), ZZH-Lucaninae (this study), Rove-Tree-11 (*67*), NHM-Carabids (*33*), and GD-Insect (this study). The five benchmark datasets comprise two plant-image datasets and three beetle-image datasets; GD-Insect provides an additional real-world insect survey dataset for validation. Selected morphological traits span floral structures, wood transverse anatomy, beetle dorsal habitus, and insect dorsal/lateral habitus.

**Table 1.**
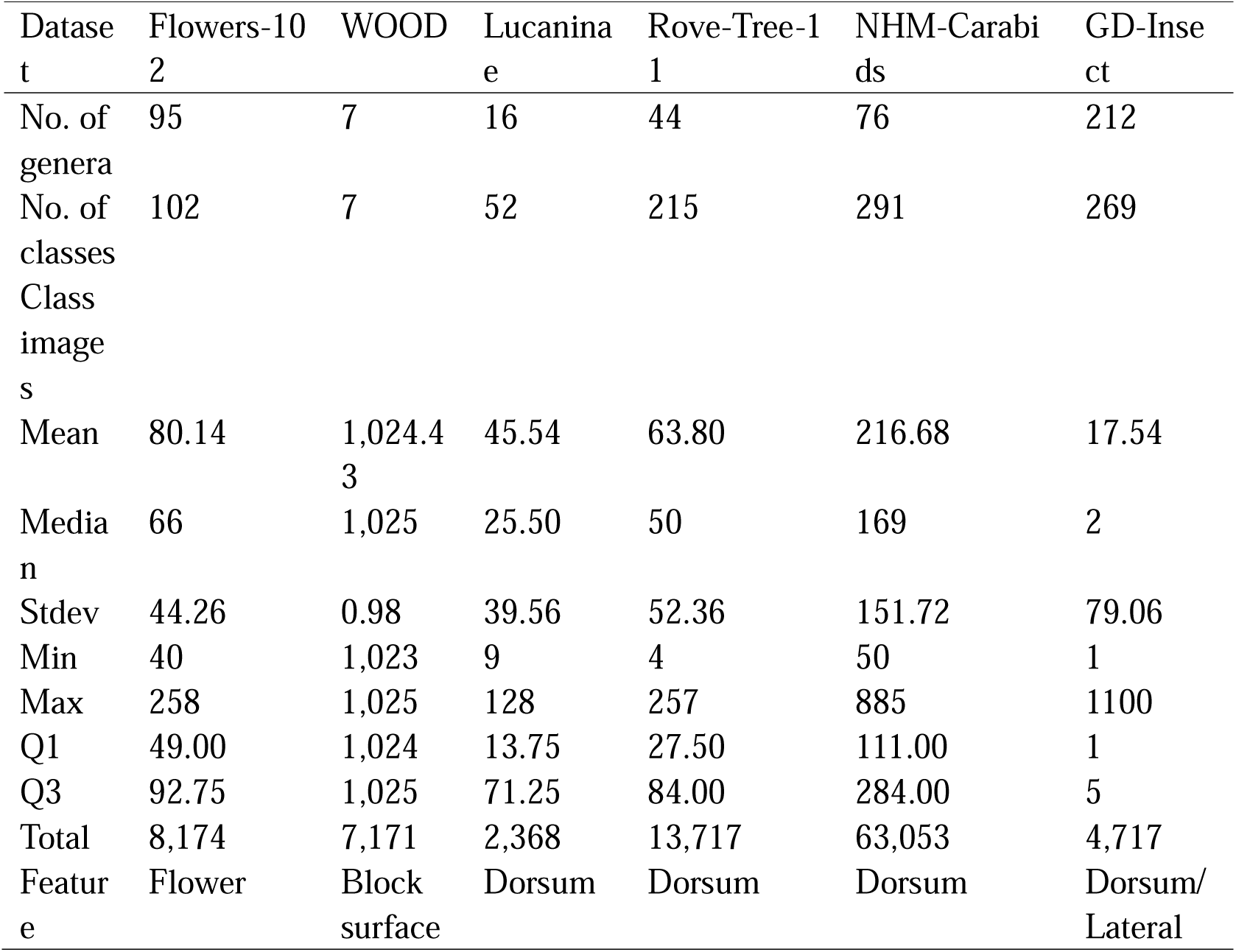
Overview of the six datasets analysed in this study. Summary statistics include the number of genera, the number of class-level categories (species, subspecies, or strains), and per-class image distribution. All numbers refer to cleaned datasets.

#### Dataset descriptions

1. Flowers-102 (*65*): The Flowers-102 dataset comprises images of 102 species from 95 genera of flowering plants. Most images are close-up views of floral structures, although non-target objects such as leaves, stems, or background vegetation may also be present. The images exhibit substantial variation in scale, pose, and illumination.
2. WOOD (*66*): The WOOD dataset includes seven wood species from multiple plant families originating from Bangladesh. Images depict transverse sections of wood captured using consumer-grade smartphone cameras. Anatomical patterns on the cut surface provide the primary morphological information relevant for species discrimination.
3. ZZH-Lucaninae (this study): This custom dataset includes 52 species from 16 genera of stag beetles. Images were captured using digital cameras and smartphone devices. After minimal cropping, each specimen is centered within the frame, and only dorsal-habitus views were retained for analysis.
4. Rove-Tree-11 (*67*): Rove-Tree-11 contains images of 215 species from 44 genera of museum-preserved staphylinid beetles. Originally curated to evaluate the phylogenetic signal contained in digitized habitus photographs (*68*), the dataset provides standardized dorsal images of pinned specimens.
5. NHM-Carabids (*33*): The NHM-Carabids dataset consists of dorsal habitus images of 291 species from 76 genera of carabid beetles from the Natural History Museum (London). Images vary widely in resolution, specimen size within the frame, and background uniformity. Some images contain relatively limited visible morphological detail because of low resolution, small specimen size within the frame, uneven backgrounds, or reduced image sharpness.
6. GD-Insect (this study): This real-world dataset was collected as part of a biodiversity survey and assessment program targeting aquatic insects in the Guangdong–Hong Kong–Macao Greater Bay Area. It contains 4,717 habitus images representing 269 species across 212 genera, 12 orders, and 68 families, with an extremely imbalanced taxonomic structure—152 species are represented by a single individual. Owing to its heterogeneity and strong long-tailed class distribution, GD-Insect was used exclusively for validating morphOTU performance under realistic diversity conditions and was not included in any comparative benchmarking experiments.

#### Image cleaning

All images were standardized before feature extraction to ensure consistent file structure, naming, and resolution across datasets. Because original files varied in naming conventions, formats, and dimensions, each image filename was prefixed with its corresponding Latin species name to facilitate downstream visualization and to guarantee unique identifiers.

Images were resized to a maximum short-edge length of 512 pixels. If the short edge exceeded this value, images were downscaled proportionally while preserving the aspect ratio. All files were converted to JPG format (*69*).

To reduce the impact of potentially mislabelled images in publicly available datasets, we created a high-confidence reference set using supervised classification. We trained an image classification model using AutoGluon-Multimodal (AutoMM) (*70*) with a ConvNeXt V2-L backbone (*45*). After fine-tuning on each dataset, the resulting model was used to predict labels for all images. Images for which the predicted label did not match the provided class label were removed from the dataset (*69*). Details of the training procedure are provided in the Training section.

#### Model architectures

We evaluated three categories of image-embedding extraction methods: (i) pretrained backbone models without task-specific training, (ii) supervised fine-tuning using AutoGluon-Multimodal (AutoMM), and (iii) the OTU-Former architecture combining self-supervised pretraining and supervised metric learning.

#### Pretrained backbone models

As baseline representations, we used the script, autogluon_training.py, to extract embeddings directly from five ImageNet-scale pretrained backbones (base-level parameter size, ∼85M): two CNNs (RegNetY-16GF, ConvNeXt V2-B), two transformer models (ViT-B, EVA02-B), and one self-supervised model (DINOv3-B). ResNet-50 (*44*) was included as an additional reference backbone.

#### Supervised learning with AutoMM

Supervised models were trained using AutoGluon-Multimodal (AutoMM) (*70*). For each dataset, images were split using stratified sampling (90% train+validation, 10% test). Data augmentation followed AutoMM’s default pipeline, including resizing, center cropping, horizontal/vertical flips, color jitter, and trivial augment. Unless otherwise noted, the maximum training run was 50 epochs, using cross-entropy loss. After fine-tuning, embeddings were extracted from the penultimate layer of the model. All supervised procedures were implemented from Zhang (*69*).

#### OTU-Former architecture

OTU-Former consists of a self-supervised pretraining (SSL) stage and a supervised fine-tuning (SFT) stage. A ViT-based encoder serves as the backbone, using the CLS token as the image-level embedding.

#### Self-supervised pretraining

The SSL stage employs a teacher–student framework updated by an exponential moving average (EMA). The student network comprises a ViT encoder followed by a 3-layer MLP projector producing fixed-dimensional L2-normalized embeddings. The teacher and student receive multi-crop views of each image with geometric and appearance augmentations.

The SSL objective combines three losses:

1. **Global self-distillation loss** between teacher and student CLS embeddings,
2. **Local-to-global distillation loss**, aligning local student crops to teacher global embeddings,
3. **Masked token regression loss**, predicting teacher patch-token outputs for masked positions.

Learning rate schedules, EMA momentum, and temperature warmup followed cosine or linear schedules as specified in Supplementary Materials.

#### Supervised fine-tuning

The SFT stage applies ArcFace metric learning (*43*) to refine class separation. For fine-tuning, the first 70% of transformer blocks were frozen, and the remaining blocks and metric head were trained. The final OTU-Former embedding is a fixed-length vector.

#### Embedding evaluation

Embeddings were evaluated using classification and clustering benchmarks, including k-nearest neighbor accuracy, linear probing performance, retrieval metrics (Recall@K, mAP), and clustering indices (NMI, ARI, silhouette score, purity). Additional metrics for metric-learning models (intra-class variance, inter-class distance, embedding norms) and UMAP visualizations (*71*) were computed using standard implementations. Linear probe accuracy was used as a post hoc measure of embedding separability rather than as an operational open-set classifier. For open-set experiments, the probe was trained and evaluated on labeled evaluation splits that included species withheld from representation learning, allowing assessment of whether unseen species formed linearly separable clusters. Retrieval metrics and clustering indices were computed using reference labels only for evaluation. For GD-Insect, singleton species were retained for morphOTU richness and Shannon-diversity calculations but excluded from metrics requiring train–test or positive-pair evaluation, including linear probing and mAP. ARI was calculated over all specimens using expert-assigned species labels unless otherwise specified.

#### Defining MorphOTUs

We delineated image-based morphological operational taxonomic units (morphOTUs) by hierarchical clustering in the learned image-embedding space, followed by threshold-based tree partitioning. Image embeddings were L2-normalized, and pairwise cosine distances were computed to construct a UPGMA dendrogram using average-linkage hierarchical clustering.

#### Partitioning across distance thresholds

To obtain clusterings at multiple resolutions, we generated a series of partitioning schemes by applying distance cutoffs across the same UPGMA tree. For each cutoff, we recorded cluster membership, the number of resulting clusters, and summary statistics describing the distribution of clade sizes. Clade stability was assessed using jackknife resampling of the original embeddings.

#### Evaluation of partition quality

Partition quality was quantified using clustering-based and label-based indices, including normalized mutual information (NMI), adjusted mutual information (AMI), adjusted Rand index (ARI), silhouette score, purity, homogeneity, completeness, V-measure, and BCubed precision/recall/F-score. Additional summaries, including splitting and lumping indices and the proportion of monophyletic groups, were computed from the partition assignments. Diversity estimates (richness and Shannon indices) were calculated under alternative filtering schemes, with optional exclusion of low-abundance morphOTUs.

#### Two-stage dynamic threshold scanning

To identify informative distance thresholds, we implemented a two-stage dynamic scanning procedure. Label-based metrics, including BCubed-F, were used only for retrospective evaluation and threshold benchmarking. They were not part of the intended label-free morphOTU workflow. In the coarse scan, cutoffs were evaluated across a broad distance range, and partition quality metrics were computed for each threshold. The per-metric optima were aggregated to define a candidate threshold window. Within this window, a fine scan was conducted using smaller step sizes, during which partition assignments, summary statistics, and diversity indices were recomputed. All clustering and threshold-scanning procedures were implemented in custom scripts and are detailed in Supplementary Methods.

#### Experimental scenarios

We evaluated morphOTU performance under multiple controlled experimental scenarios designed to reflect conditions commonly encountered in image-based biodiversity assessments. All scenarios were applied using the same embedding extraction and clustering procedures described above. Unless otherwise noted, supervised learning (SL) and supervised fine-tuning (SFT) used stratified dataset splits of 90% for training and 10% for testing.

#### Baseline closed-set evaluation

Baseline experiments assessed model performance when all test classes were also present in the training set. For SL, we used four backbone families—ConvNeXt V2, RegNetY, ViT, and EVA02—each represented by three parameter-scale variants (large, medium, and small; Table S2). For SSL and SSL+SFT, we used ViT-T, ViT-S, and ViT-B as backbones. SSL was trained on all available images for each dataset; during SFT, 90% of the images were used for fine-tuning, with the remaining 10% reserved for testing.

#### Open-set evaluation with unknown species

To assess performance when the test data include classes absent from training, we constructed nested open-set datasets in which the proportion of unknown-class images in the test set was set to 0%, 25%, 50%, and 75%. The corresponding complements served as closed-set data, generating the datasets Dun0, Dun25, Dun50, and Dun75 (Table S1). For each proportion, we evaluated SL, SSL, and SSL+SFT models using identical backbone architectures.

#### Impact of labeled-data availability

To examine the dependence of SFT on labeled sample size, we varied the proportion of labeled images available during SFT while holding the SSL stage constant. Using the Dun25 split as the base scenario (25% open-set test, 75% used for SSL), we subsampled the labeled portion of the training set to 75%, 50%, 25%, and 10% of its original size, generating the nested datasets Dun25, Dun25_FT50, Dun25_FT25, and Dun25_FT10. SSL was conducted using all images available under the Dun25 scenario, while only the subsampled labeled fractions were used in the ArcFace SFT stage.

#### Limited images per species

To evaluate the effect of reduced within-class sample diversity, we controlled the number of training images per species while keeping the set of species fixed. Using the NHM-Carabids dataset as an example, species with at least 105 images were designated as the closed-set. For each species, we randomly sampled 100, 50, 25, or 10 images for training. Remaining images from these species formed the closed-set test subset, while all other species served as open-set test classes. This produced the nested datasets Dsub100, Dsub50, Dsub25, and Dsub10 (Table S1).

#### Cross-dataset generalization

To test cross-dataset transfer, we trained SL, SSL, and SSL+SFT models on one beetle dataset (A) and evaluated them on the other two datasets (B and C). In addition to direct transfer, we evaluated an additional variant (SSL_newSFT) in which SSL pretrained weights from dataset A were fine-tuned on the labeled subset of dataset B or C using ArcFace. All other procedures were kept consistent with the baseline and open-set scenarios.

#### Low-quality images in training

To quantitatively assess the effect of image degradation on representation learning, we conducted a controlled downsampling experiment using the high-quality ZZH-Lucaninae dataset. Images were resized to fixed short-edge lengths of 512, 256, 128, 64, and 32 pixels to simulate progressively lower levels of visual detail. For each resolution, SL, SSL, and SSL+SFT models were trained independently using the corresponding downsampled images. Embedding quality was evaluated using the original 512-pixel test images to ensure comparability across conditions.

Because real-world image degradation can arise from multiple factors, including focus, illumination, contrast, motion blur, and specimen size within the frame, short-edge downsampling serves as a controlled and quantifiable proxy rather than a full representation of all quality loss modes. Nevertheless, this procedure provides a standardized way to examine how reduced image resolution influences the ability of different learning paradigms to extract stable morphological features.

#### Test in a real-world dataset

To evaluate the performance of OTU-Former under realistic biodiversity conditions, we conducted a two-stage assessment (SSL and SSL+SFT) using the GD-Insect dataset. During supervised fine-tuning, only species represented by ≥30 individuals were used (28 species in total, accounting for 10.41% of all species). This design simulates a common scenario in biodiversity surveys in which only a small subset of widespread and easily identifiable species can be reliably annotated, while the majority of taxa remain unlabeled.

For OTU construction, a single fixed cosine-distance threshold of 0.3 was applied to the UPGMA tree. This value was chosen based on empirical observations from the three beetle datasets and provides a consistent comparison point for diversity estimation. It should be noted that we used 0.3 as a fixed benchmark threshold, not as an optimized dataset-specific cutoff, to test whether a threshold derived from independent beetle datasets could provide usable diversity estimates in a heterogeneous survey dataset. The resulting morphOTUs were then used to compute richness and Shannon diversity indices.

#### Morphological interpretability analysis

To examine the spatial focus of model features, we generated visual explanation maps using Score-CAM (*55*) applied to trained ViT-T models. For each beetle dataset, we selected three genera comprising closely related species and distinct sample sizes (Table 2). Heatmaps were produced for representative images and were inspected alongside clustering and embedding patterns. Score-CAM visualizations were implemented using GradCam_heatmap.py.

**Table 2.** Subsets consisting of all images from a single genus. Each subset contains all available images for the listed genus; species counts, and per-species image means are calculated after dataset cleaning.

| Dataset | Genus | No. species | No. images | Mean per sp. |
| --- | --- | --- | --- | --- |
| Lucaninae | *Epidorcus* | 2 | 230 | 115.00 |
| Lucaninae | *Prosopocoilus* | 7 | 313 | 44.71 |
| Lucaninae | *Lucanus* | 21 | 879 | 41.86 |
| Rove-Tree-11 | *Paederus* | 3 | 458 | 152.67 |
| Rove-Tree-11 | *Lathrobium* | 11 | 1,175 | 106.82 |
| Rove-Tree-11 | *Philonthus* | 46 | 3,411 | 74.15 |
| NHM-Carabids | *Ophonus* | 9 | 1,645 | 182.78 |
| NHM-Carabids | *Amara* | 21 | 4,635 | 220.71 |
| NHM-Carabids | *Bembidion* | 51 | 13,265 | 260.10 |

## Supporting information

Supplementary information, including methods, figures, tables, and scripts for this study

## Supplementary Materials

All supplementary materials, including Supplementary Figures S1–S29, Supplementary Tables S1–S5, Supplementary Methods, and related analysis scripts, are available online. References (72–78) are cited only in the Supplementary Materials.

## Data Availability

All datasets, scripts, and code generated or applied in this study are uploaded and deposited in Dryad with the following link: http://datadryad.org/share/LINK_NOT_FOR_PUBLICATION/GAKk7UXRPMkpb03QlNbTHeZlEeqUsiSx8_3Av4ejif8 for peer-review only. These data will be released immediately upon the acceptance of the manuscript.

## Author Contribution

ZZ conceived the study, designed the analyses, performed data processing and model experiments, and drafted the manuscript. MY contributed to dataset curation, taxonomic validation, and interpretation of biological results. MO contributed to study refinement and provided critical revisions to the manuscript. WC assisted with computational analysis and manuscript editing. XL contributed to data organization and quality control. LY assisted in algorithm evaluation and model validation. XS supervised the biological components and provided key scientific guidance. FZ supervised the computational framework, coordinated the project, secured funding, and finalized the manuscript.

## Materials & Correspondence

Correspondence and material requests should be addressed to **F. Zhang**.

## Use of Artificial Intelligence

We used AI-assisted tools in limited capacities, specifically Claude-Sonnet-3.5 for improving Python scripts and GPT-5.1-Instant for refining the English writing of the manuscript. All scientific ideas, study design, analyses, and interpretations were conceived and validated solely by the authors.

