## Supplementary information, including methods, figures, tables, and scripts for this study for "When AI encounters natural history: Morphological OTUs reshape our understanding of Earth’s life"

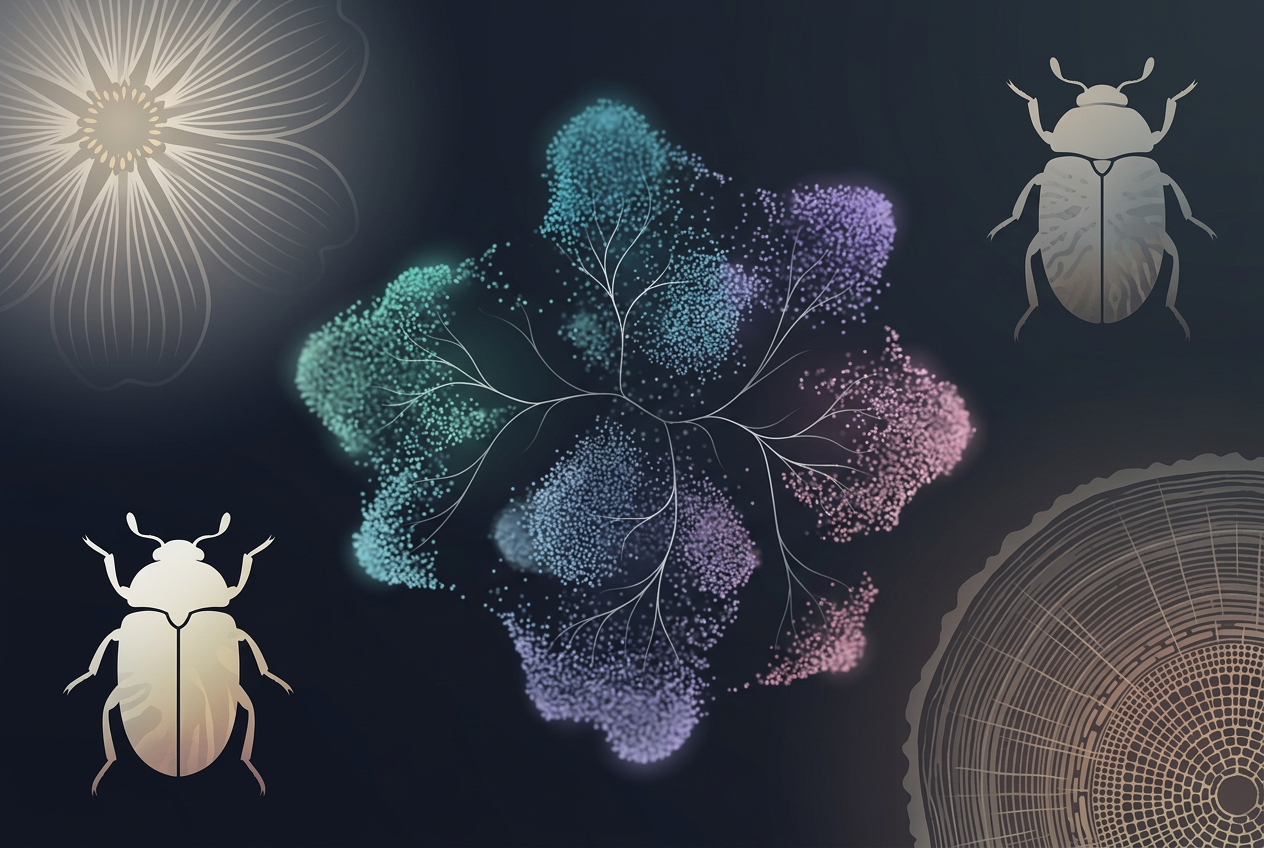

**Contents**

**Supplementary Methods**

**S1.** Image Pre‑processing and Cleaning

**S2.** Pretrained Backbone Models

**S3.** Supervised Learning (SL) Pipeline

**S4.** OTU‑Former Architecture

**S5.** Defining morphOTU

**S6.** Case scenarios

**Supplementary Tables**

**Table S1.** Dataset partitioning schemes and nomenclature conventions.

**Table S2.** Dataset partitioning rules (Dun, Dsub, FT, genus subsets).

**Table S3.** Default hyperparameters for self-supervised pretraining.

**Table S4.** Default augmentation for global and local crops.

**Table S5.** Default hyperparameters for ArcFace finetuning.

**Supplementary Figures**

**Fig. S1–S4.** Metrics embeddings for different models and scenarios

**Fig. S5–S6.** Distance models for all and ViT

**Fig. S7–S16.** Distance distribution for raw cosine and cosine among five datasets

**Fig. S17–S19.** Effects of open‑set and SSL/SFT combinations on embedding metrics across datasets

**Fig. S20–S21.** Scenario‑wise performance comparison under raw, SL_train_size, SSL_train_size, SFT_finetune_size, and class_size

**Fig. S22.** Cosine‑distance comparison across scenarios for all five datasets

**Fig. S23.** Intra‑class cosine‑distance distribution under all scenarios across datasets

**Fig. S24–S26.** Pretrained vs SL vs SSL vs SSL_SFT comparisons across multiple backbones

**Fig. S27.** Additional scenario‑wise comparisons across datasets and treatments

**Fig.** **S28.** Genus‑level evaluation of morphOTU performance across nine closely related beetle groups

**Supplementary Scripts**

- **clean_figs.py**
- **detect_bounding_box.py**
- **download_TreeOfLife-200M.p**
- **embeddings_tree.py**
- **autogluon_training.py**
- **GradCam_heatmap.py**
- **SSL.py**
- **split_dataset.py**

**References (71–77)**

### **Overall**

This Supplementary Information provides the full methodological, technical, and diagnostic details supporting the analyses presented in the main manuscript. All extended descriptions, parameter settings, and implementation details that exceed the scope of the main Methods section are compiled here to ensure transparency, reproducibility, and ease of review.

The SI contains three major components:

(1) Supplementary Methods,

(2) Supplementary Tables,

(3) Supplementary Figure, as well as all scripts used in the study.

First, the Supplementary Methods include detailed procedures for image cleaning, dataset preprocessing, and error filtering, including the full AutoGluon pipeline used to detect and remove mislabeled images. We also provide the complete specifications of all model architectures—pretrained backbones, supervised AutoMM models, and the full OTU‑Former design—together with hyperparameters, augmentation strategies, optimization schedules, and training stability considerations for both the SSL and SFT stages. Technical details for embedding extraction, clustering evaluation, metric‑learning diagnostics, and UMAP visualization are also documented.

Second, the SI contains the complete algorithmic description of the MorphOTU pipeline, including:

• hierarchical clustering procedures,

• cosine‑distance computation and normalization,

• multi‑threshold partitioning,

• bootstrap‑based clade stability analysis, and

• the two‑stage dynamic threshold scanning method (coarse and fine scans).

All clustering quality metrics, diversity estimation procedures, and abundance‑filtering strategies (analogous to metabarcoding noise filtering) are provided in full.

Third, we provide complete specifications of all experimental scenarios evaluated in the study, including open‑set tests, labeled‑data‑size reduction, limited images per species, and cross‑dataset generalization—together with dataset partitioning rules for all nested datasets (Dun0/25/50/75, Dun25_FT subsets, Dsub subsets, and genus‑level subsets).

The Supplementary Figures present embedding distributions, distance diagnostics, threshold‑scanning curves, diversity‑index comparisons, UMAP visualizations, and Score‑CAM heatmaps used to evaluate model interpretability. Supplementary Tables summarize dataset statistics, OTU‑Former hyperparameters, threshold‑scan outputs, and cluster‑quality summaries. All scripts used in preprocessing, model training, embedding extraction, clustering, visualization, and interpretability analysis (e.g., clean_figs.py, extract_embeddings.py, GradCam_heatmap.py) are listed and documented.

### **S1. Dataset and Image Preparation**

### **1.1 Dataset Selection**

The concept of morphOTU requires using standardized morphological traits—analogous to the use of the COI universal barcode in animals—so that images within and across datasets remain morphologically comparable. Most publicly available biodiversity image datasets contain heterogeneous traits, multiple viewpoints, varied backgrounds, or inconsistent imaging conditions, making them unsuitable for the present study.

To meet morphOTU criteria, we assembled five datasets, including two plant datasets and three beetle datasets. The chosen morphological traits were:

- flowers (Flowers‑102),
- wood transverse-section texture (WOOD),
- dorsal body view of beetles (ZZH‑Lucaninae, Rove‑Tree‑11, NHM‑Carabids).

After cleaning, dataset sizes ranged from 2,368 to 63,053 images, with 7–291 classes, representing typical small‑ to medium‑scale biodiversity datasets. Except for WOOD, all datasets display strong long‑tailed distributions, consistent with real-world species abundance patterns.

Below, we briefly describe each dataset (two representative images may be shown for each):

**Flowers‑102 (*64*):** This dataset contains 95 genera and 102 flower species. Most images are close‑up shots where flowers occupy a substantial portion of the frame, though leaves, stems, and non‑biological backgrounds may be present. Images vary widely in scale, pose, and illumination.

**WOOD (*65*):** This dataset includes seven wood species from Bangladesh. Each sample is photographed using a smartphone, focusing on the transverse wood block surface, where texture and pore patterns may serve as key diagnostic features.

**ZZH‑Lucaninae:** A custom dataset created by our team, containing 16 genera and 52 species of Lucaninae beetles. Images were taken using a Canon R5 Mark II camera and smartphone, followed by light cropping to ensure that the beetle dorsum is centered in the frame.

**Rove‑Tree‑11 (*66*):** This museum-based dataset includes 44 genera and 215 species of rove beetles. The original study used images to test phylogenetic signal recovery from digital images (*67*).

**NHM‑Carabids (*33*):** Collected from the Natural History Museum (London), this dataset contains 76 genera and 291 species of carabid beetles. The original study showed that CNNs can classify beetle dorsum images at the species level. However, compared with the other datasets, NHM‑Carabids exhibits lower and more variable image quality, with many images showing low resolution, small specimen size, motion blur, or cluttered backgrounds—patterns that are also reflected in its consistently lower performance in downstream analyses.

### **1.2. Image Cleaning**

The five datasets exhibit substantial differences in file naming conventions, image resolution, and overall quality. To create a clean and standardized benchmark for representation-learning experiments, we implemented the following procedure:

**(1) File renaming and class-prefix normalization**

To simplify visual inspection and ensure filename uniqueness, each image was given a Latin-name class prefix. This was especially helpful for cross-dataset comparisons and embedding visualization.

**(2) Resolution standardization**

If the short side exceeded 512 pixels, the image was resized proportionally so that the short side became 512 pixels.

All images were stored in JPEG (.jpg) format (implemented in clean_figs.py).

(3) Automated identification of mislabelled samples

Public biodiversity datasets can contain mislabelled images. To build a high-quality, “error‑free” reference set:

- We fine-tuned an AutoGluon-Multimodal (AutoMM) image-classification model (*69*).
- The backbone was ConvNeXt V2‑L (*45*), a large-capacity vision model.
- After supervised training, all images were re-predicted.
- Images whose predicted labels conflicted with the dataset labels were removed (extract_embeddings.py).

Details of the training procedure are provided in the Training section.

### **S2. Pretrained Backbone Models**

In the Pretrained condition, embeddings were extracted directly from backbone models without any additional training (implemented in extract_embeddings.py). These pretrained embeddings serve as a baseline for comparing against all subsequent SL‑, SSL‑, and SFT‑based models.

We selected base‑level architectures (~85M parameters) across both CNN and Transformer families to ensure comparable model capacity. The following pretrained backbones were used:

- RegNetY‑16GF (*46*): CNN
- ConvNeXt V2‑B (*45*): CNN
- ViT‑B (*47*): Vision Transformer
- EVA02‑B (*48*): Vision Transformer
- DINOv3‑B (*49*): Self‑supervised Vision Transformer

All pretrained weights were accessed through the AutoGluon‑Multimodal model zoo.

Embeddings were extracted using standard preprocessing pipelines provided by AutoMM.

### **S3. Supervised Learning (SL)**

We used AutoGluon‑Multimodal (AutoMM) to perform supervised training, data augmentation, model fine‑tuning, classification evaluation, and embedding extraction (extract_embeddings.py). Unless otherwise specified, stratified sampling was used for all dataset splits.

Dataset partitioning for SL followed:

- 90% for training + validation
- 10% for held‑out testing (implemented via split_dataset.py)

Within the 90% portion, AutoMM automatically performs stratified and adaptive‑ratio sampling to create internal training/validation splits.

AutoMM’s augmentation pipeline used the following transformations:

- resize_shorter_side
- center_crop
- random_horizontal_flip
- random_vertical_flip
- color_jitter
- trivial_augment

These augmentations help mitigate variation in scale, pose, illumination, and viewpoint across datasets.

Training configuration:

- Maximum epochs: 50
- Loss function: Cross‑Entropy (AutoMM default)
- Backbone initialization: from pretrained weights listed in Section 2.1
- Evaluation + embedding extraction: performed after training using AutoMM’s built‑in inference module

The resulting embeddings form the SL condition for all downstream comparisons.

### **S4. OTU‑Former Architecture**

OTU‑Former is a two‑stage representation learning framework consisting of (i) self-supervised pretraining (SSL) and (ii) supervised metric finetuning (SFT). The two stages can be used independently or in combination. The design goal is to obtain high-quality morphological embeddings with strong intra-class compactness and inter-class separation, suitable for morphologically derived OTUs (morphOTUs). Default hyperparameters are provided in **Table S2**.

### **4.1 Overview of the Two‑Stage Framework**

The architecture follows a DINO‑style teacher–student self‑distillation paradigm (*36*) with an additional masked token regression objective in Stage 1, followed by ArcFace-based metric supervision (*43*) in Stage 2.

We adopt ViT-family backbones, and unless noted otherwise, the CLS token embedding is used as the image-level representation for downstream tasks (*47*).

A motivation for this architecture is that mainstream SSL methods often become unstable or collapse on small and medium-sized datasets (*71–75*). OTU‑Former includes several stabilization strategies—EMA teacher, teacher centering, cosine schedules—to explicitly address these issues.

### **4.2 Stage 1: Self-Supervised Pretraining (Teacher–Student Distillation + Masked Token Regression)**

*Backbone and Projector*

The student network is a ViT encoder producing token sequence

$\left( \text{CLS},\text{patch}_{1},\ldots,\text{patch}_{N} \right)$.

A 3‑layer MLP projector (GELU activations; output dimension = 256) maps the CLS token to a unit‑normalized embedding.

The teacher network shares the same architecture, but its weights are updated via an exponential moving average (EMA) of the student:

$$\theta_{t}\leftarrow m\text{ }\theta_{t}+(1-m)\text{ }\theta_{s}$$

with cosine-scheduled momentum from 0.995→0.999, stabilizing training (*36*, *76*).

To further suppress collapse, we maintain a teacher-embedding center updated with 0.9 EMA, consistent with self-distillation literature.

*Multi‑crop Views and Augmentation*

We adopt multi-crop training (*36*), which is particularly effective for fine-grained insect and floral structures:

- Two global crops (224 px): preserving coarse body shape and overall morphology
- Six local crops (96 px): emphasizing fine anatomical and textural details
- Strong geometric augmentation:
  - Random Rotation up to 180°
  - Horizontal Flip / Vertical Flip
- Photometric augmentation: ColorJitter, RandomGrayscale
- Blur augmentation: Gaussian blur (larger kernel for global crops)
- ImageNet mean–std normalization

Large rotations and vertical flips are essential for datasets in which insects or flowers may appear at arbitrary orientations.

*SSL Objectives*

The SSL loss integrates three complementary objectives:

1. Global‑to‑global self-distillation loss
   - Aligns student CLS embeddings to teacher CLS embeddings
   - Teacher temperature follows a linear warmup schedule to stabilize early SSL training
   - Encourages global semantic alignment
2. Local‑to‑global distillation loss
   - Aligns local-crop student embeddings to teacher global embeddings
   - Strengthens representation of fine local details
3. Masked token regression (patch-level alignment)
   - Randomly mask 50% of patch tokens
   - Align normalized student and teacher patch embeddings via cosine regression
   - Captures fine-grained textures and structural cues important for beetles, flowers, and wood grain

These objectives jointly encourage robust, transferable representations capable of supporting fine-grained taxonomic distinctions.

*Training Schedules and Optimization*

- Cosine schedules for learning rate and EMA momentum
- Linear warmup for teacher temperature
- Student optimized with AdamW (LR 5×10⁻⁴, weight decay 0.05)
- Gradient clipping (max-norm 3.0) to mitigate instability on small datasets

The SSL stage yields embeddings with strong generalization capacity even before supervised finetuning.

### **4.3 Stage 2: Supervised Metric Finetuning (ArcFace)**

To adapt embeddings for downstream classification and species-level discrimination, we apply ArcFace metric learning (*43*). This improves intra-class compactness and inter-class separation:

*Finetuning Strategy*

- Initialize encoder from the Stage‑1 teacher checkpoint
- Freeze the first 70% of ViT blocks to avoid overfitting on limited labeled data
- Finetune the remaining transformer blocks + the metric head
- Metric embedding dimension = fixed (default is 256)

*Metric Head and Loss*

The supervised head is:

Linear(embed_dim→512) → ReLU → Linear

We apply ArcFace with:

- scale $s=64$
- margin $m=28.6$(value interpreted according to implementation)

Optimization:

- AdamW, LR $=1\times{10}^{-4}$, weight decay 0.05
- ArcFace parameters are optimized jointly with the model

This stage outputs fixed‑dimensional normalized embeddings optimized for taxonomic separability.

### **4.4 Embedding Evaluation Metrics**

To characterize representation quality, we evaluate embeddings across four dimensions. These metrics are supported by literature summarized in *Embedding_Metrics_Literature_Summary.md.*

(1) Classification Transferability

- k‑NN accuracy (k = 1, 5, 20)
- Linear probing accuracy
  Widely used in SSL research (*53*, *77*).

(2) Retrieval Performance

- Recall@K (1, 5, 10)
- mAP
  Standard in metric learning and retrieval benchmarks.

(3) Clustering Quality

- NMI, ARI, Silhouette Score, Purity
  Evaluates label‑cluster agreement and geometry.

(4) Metric‑Learning‑Specific Geometry Metrics

- Intra‑class variance
- Inter‑class distance
- Embedding norms
  Reflect class compactness, separation, and geometric stability.

(5) Embedding Visualization

We use UMAP (umap-learn v*0*.*5*.*1*) (*70*) for low-dimensional visualization of high‑dimensional embeddings.

### **4.5 Summary**

OTU‑Former integrates a stabilized teacher–student SSL pretraining stage with metric learning-based supervised finetuning to produce high-quality morphological embeddings.

Key innovations include:

- Collapse-resistant SSL design (EMA teacher, centering, multi-crop, temperature warmup)
- Patch-level masked regression for fine-grained morphological cues
- ArcFace-based metric finetuning for robust class separation
- Comprehensive embedding evaluation across transfer, retrieval, clustering, and geometry metrics

### **S5. Defining morphOTUs**

We delineated morphological operational taxonomic units (morphOTUs) directly from learned image embeddings by hierarchical clustering on a cosine-distance matrix, followed by systematic partitioning across a range of distance cutoffs. This procedure expands the concise description in the main text (***morphOTU_methods_main_text.md***) with full implementation details.

### **5.1 Pairwise Distances and Tree Construction**

For each specimen, we extracted a fixed-embedding from the chosen backbone or OTU‑Former model. Embeddings were L2-normalized, and pairwise cosine distances were computed to form a specimen-by-specimen distance matrix. To reduce potential distortions in high-dimensional spaces:

- Optional PCA whitening was applied to stabilize pairwise distances before cosine computation.
- Optional local scaling (Mutual‑Proximity style) rescaled distances using each specimen’s k‑nearest‑neighbour distance as a local density factor, reducing the effect of heterogeneous local density in the embedding space.

From the resulting distance matrix, we constructed a UPGMA dendrogram (average-linkage hierarchical clustering), which served as a unified backbone for all subsequent morphOTU partitions.

We examined global pairwise distance distributions and, when available, intra-class distance summaries to identify candidate regions where a “gap‑like” separation may occur between within‑ and between-species distances. These diagnostics were treated as heuristic, not as evidence of a universal barcoding gap.

### **5.2 No Universal Species Threshold for Morphology**

Morphological disparity varies substantially across lineages, so we did not assume any universal species-level cutoff (e.g., DNA barcoding’s 0.97 rule). Instead, we explicitly modeled morphOTU delimitation as a family of hierarchical partitions, each corresponding to a distance threshold on the same UPGMA tree.

For each cutoff, we recorded:

- the resulting number of morphOTUs,
- the composition of clusters,
- splitting and lumping relative to reference species labels (when available),
- and partition quality metrics.

This produced a full “morphOTU profile” across distance space.

### **5.3 Bootstrap Support for Internal Clades**

To evaluate the internal stability of clades that underlie morphOTU partitions, we conducted bootstrap resampling of embedding dimensions (feature resampling):

1. Sample a subset of embedding dimensions with replacement.
2. Recompute pairwise distances.
3. Rebuild the UPGMA dendrogram.
4. Record whether each internal clade is recovered.

The bootstrap proportion then quantifies clade stability, providing an internal support value analogous to phylogenetic bootstraps.

### **5.4 Partition Quality Metrics**

We evaluated morphOTU partitions using a comprehensive suite of complementary metrics, drawing from cluster validation, information theory, and metric-learning literature (summarized in ***embedding_metrics_literature_summary.md***).

Clustering quality metrics

- NMI, AMI, ARI
- Silhouette Score
- Purity

Label-based agreement indices (when reference labels available)

- Homogeneity / Completeness / V-measure
- BCubed Precision / Recall / F-score

Partition summary statistics

- Splitting and lumping indices
- Proportion of monophyletic species
- OTU-to-species ratios

These metrics capture how well morphOTU partitions reflect known species structure and how cluster cohesion/separation behaves across cutoffs.

### **5.5 Evaluating the Practical Consequences of a Single Cutoff**

Ideally, morphOTU delimitation would incorporate expert-guided inspection, for instance, visually confirming morphological coherence via CAMs or consulting taxonomic expertise. Because such inspection is rarely feasible at scale, we evaluated the practical consequences of selecting a single (possibly suboptimal) cutoff.

For each cutoff, we computed diversity indices analogous to metabarcoding analyses:

- richness,
- Shannon diversity,
- Simpson diversity.

We computed these indices under three filtering regimes:

1. No filtering,
2. Filtering morphOTUs with abundance ≤ 2,
3. Filtering morphOTUs with abundance ≤ 5

(similar to noise filtering in metabarcoding pipelines).

We compared these diversity summaries against ground-truth label summaries to assess the practical validity of morphOTUs generated from a single threshold.

Across datasets, BCubed F‑score tended to produce the most stable optimal thresholds, ARI performed moderately well, and the Silhouette score was consistently the least reliable.

### **5.6 Two‑Stage Dynamic Threshold Scanning**

To identify plausible distance cutoffs, we developed a two-stage dynamic threshold scanning procedure, combining coarse exploration with fine localization.

Step 1: Coarse scan

- Scan cutoffs in the range [0.05, 1.00]
- Step size 0.05
- For each cutoff:
  - Generate morphOTU partitions
  - Compute the six key quality metrics
    (NMI, ARI, AMI, BCubed F-score, V-measure, Silhouette score)

For each metric, we identify its operational optimum, typically the cutoff yielding the maximum metric value.

Step 2: Aggregation and fine scan

We aggregate all per-metric optima to define a candidate cutoff window, and then expand this window by ±0.15, clipping the final interval to [0.05, 1.00].

Within this refined interval:

- perform a fine scan (step = 0.01 or 0.02),
- compute all partitions and partition-quality metrics,
- compute richness and diversity indices (with and without abundance filtering).

This yields a robust and interpretable profile of morphOTU performance around the dynamically identified threshold region.

### **5.7 Summary**

The morphOTU delimitation process:

- avoids assuming universal species-level cutoffs,
- evaluates cluster structure across a continuous range of distance thresholds,
- quantifies clade stability via bootstrap resampling,
- integrates multiple complementary partition-quality metrics,
- and examines the practical consequences of cutoff selection for diversity estimation.

These steps collectively make morphOTUs a transparent, reproducible, and taxonomy-agnostic framework for species-level grouping based solely on learned morphological representations.

### **S6. Case Scenarios**

To evaluate the applicability, robustness, and practical limits of morphOTU delimitation from image embeddings, we conducted a series of controlled case scenarios. These scenarios go beyond the baseline closed‑set evaluation and emulate realistic challenges encountered in biodiversity research, including open‑set conditions, limited labeled data, and extreme class imbalance. Dataset partition rules are summarized in **Table S1**.

### **6.1 Baseline Test (Closed‑set Evaluation)**

Because embeddings extracted directly from pretrained backbones are not expected to perform well, our baseline focuses on scenarios requiring model training or fine‑tuning. The closed‑set test—where all test classes also appear in the training set—provides an upper bound (“ceiling”) for all evaluation metrics and serves as the primary reference for subsequent comparisons.

*Supervised Learning (SL) Baseline*

For SL, we selected four backbone families:

- CNNs: ConvNeXt v2, RegNetY
- ViTs: ViT, EVA02

Each backbone family includes three model sizes:

- Large (~85M parameters)
- Medium (~20M parameters)
- Small (~5M parameters)

(Details in Table XXX.)

Dataset splitting follows the default 90% train + validation / 10% test scheme, denoted Dun0.

*SSL and SSL→SFT Baseline*

For SSL and subsequent SFT (ArcFace fine‑tuning):

- We used ViT‑T, ViT‑S, and ViT‑B as backbones.
- All images were used in SSL pretraining.
- SFT used the same 90% training and 10% test partition as SL.

In baseline evaluation, we report embedding quality, distance distributions, partition quality, and diversity metrics. We additionally replaced species labels with genus‑level labels (Dun0_genus) to provide a simplified contrast, anticipating that genus‑level morphOTUs should be easier to recover.

### **6.2 Scenario 1: Open‑set Evaluation with Unknown Species**

A crucial use case for morphOTUs is the discovery or characterization of unknown species that do not appear in the training set. We therefore examined whether SL, SSL, and SSL→SFT embeddings produce meaningful structure in open-set scenarios.

We constructed nested datasets where the proportion of unknown (open‑set) test images was: 0%, 25%, 50%, 75%

The remaining 100%, 75%, 50%, 25% of images formed the closed‑set (training + validation) portion. These datasets correspond to: Dun0, Dun25, Dun50, Dun75 (Table S1).

Because ViT‑T performed consistently well in baseline tests, all subsequent scenario analyses used ViT‑T backbones, ensuring that observed differences primarily reflect scenario conditions rather than model‑architecture effects.

We evaluated how the generalization ability of SL and SSL_SFT models changed as the amount of open‑set data increased, focusing on: cluster structure preservation, morphOTU stability, and diversity-estimate accuracy under open‑set conditions.

### **6.3 Scenario 2: Impact of Labeled Data Size (ArcFace SFT Sensitivity)**

For SL, the size of labeled data is equivalent to total training data, so no special analysis is needed. However, ArcFace SFT is expected to be sensitive to the number of labeled specimens.

This scenario reflects real practice where: (1) Many images exist, (2) But experts can only identify a fraction of them, (3) While SSL pretraining can still use *all* images.

We tested SFT sensitivity to labeled data size under the Dun25 framework: (1) Dun25 has 25% open-set test images, (2) The remaining 75% of images were used for SSL pretraining, (3) We then used subsets of the remaining images for SFT: 75%, 50%, 25%, **10%** of total images, corresponding to Dun25, Dun25_FT50, Dun25_FT25, Dun25_FT10. All subsets were nested.

We evaluated how decreasing labeled data affects embedding quality, morphOTU accuracy, and cluster geometry, providing insights into the minimum annotation effort required for useful morphOTUs.

### **6.4 Scenario 3: Limited Images per Species / Highly Imbalanced Classes**

Many biodiversity datasets are long‑tailed, with few images for most species. To assess how reduced intra-class diversity affects generalization, we fixed the class set but varied the number of images per class in training.

Using the NHM‑Carabids dataset, we: (1) selected classes with >105 images as closed‑set species. (2) for each class, randomly sampled 100 images as training data; (3) remaining images of these species were used as closed-set test data; (4) species absent from training were treated as open-set test data.

We then subsampled training species to produce: Dsub100, Dsub50, Dsub25, Dsub10, corresponding to 100, 50, 25, and 10 images per class. All subsets were nested.

This scenario evaluates sensitivity to shrinking coverage of morphological variation.

### **6.5 Scenario 4: Extreme Open‑set Transfer Across Datasets**

To test the cross-lineage transferability of embeddings, we examined an extreme scenario in which the entire test set consists of species unseen during training.

Across the three beetle datasets (A, B, C), we evaluated: (1) SL (trained on A) → tested on B and C; (2) SSL (trained on A) → tested on B and C; (3) SSL→SFT (trained on A) → tested on B and C

We additionally introduced a new SFT variant:

- SSL_newSFT:
  - Use SSL pretrained on dataset A
  - Perform ArcFace SFT on dataset B or C
  - Then evaluate on that same B or C.

This scenario evaluates whether SSL learns lineage‑general morphological features and whether rapid adaptation via SFT can restore performance when switching to a new taxonomic group.

### **6.6 Scenario 5: Morphological Interpretability (Score-CAM Analysis)**

To verify whether SL and OTU‑Former models learn meaningful morphological features (rather than background noise), we applied Score-CAM (*55*) using the ViT‑T model from baseline tests (script: *GradCam_heatmap.py*).

We examined: (1) whether attention focuses on the organism rather than the background, (2) which anatomical regions the model considers discriminative, and (3) whether expert taxonomists agree with the highlighted regions.

*Subset Selection for Interpretability*

For each beetle lineage, we constructed three subsets of closely related species by:

1. Selecting three genera that: (a) exhibit overlapping or adjacent clusters in UMAP space, (b) show apparent morphological similarity, (c) differ markedly in species richness or image abundance.
2. Extracting images for these genera.
3. Generating Score-CAM heatmaps for each specimen.

This scenario also evaluates the performance of OTU‑Former on very small datasets (as small as 230 images), providing a stress test of interpretability and generalization under extreme data constraints.

### **6.7 Summary**

Across all scenarios—closed‑set baseline, open‑set generalization, decreasing labeled data, reduced intra-class diversity, cross‑dataset transfer, and interpretability tests, our analysis evaluates not only embedding quality but also the practical usability of morphOTUs under real-world constraints.

These case scenarios collectively reveal: (1) how much labeled data is needed for robust SFT, (2) when SSL or SSL→SFT is preferable to SL, (3) how morphOTUs behave under open-set and cross-lineage conditions, and (4) whether learned features correspond to meaningful morphological structures.

### **Supplementary Tables**

**Table S1. Dataset partitioning schemes and nomenclature conventions.** Nomenclature abbreviations are as follows: D, dataset; un, unknown; FT, fine-tuning; G, genus; sub, subset. Values expressed as percentages (%) represent the proportion of the total image pool, while integers denote absolute image counts.

| Dataset | Train data  (validation data included in SL) | Test data from  closed-set classes | | Test data from  open-set classes | Note |
| --- | --- | --- | --- | --- | --- |
| Dun0 | 90% | | 10% | 0% | Unknown classes lacking. |
| Dun0_genus | 90% | | 10% | 0% | Genus name as the label. |
| Dun25 | 67.5% | | 7.5% | 25% | Unknown test images are nested among three. |
| Dun50 | 45% | | 5% | 50% |  |
| Dun75 | 22.5% | | 2.5% | 75% |  |
| Dun25_FT10 | 10% | | | 25% | 10%, 25%, 50%, and 75% (Dun25) of the data used for ArcFace fine-tuning are nested. |
| Dun25_FT25 | 25% | | |  |  |
| Dun25_FT50 | 50% | | |  |  |
| Dsub10 | 10 per class | | Remaining from known classes | Open-set classes | 10, 25, 50, 100 are nested images. |
| Dsub25 | 25 per class | |  |  |  |
| Dsub50 | 50 per class | |  |  |  |
| Dsub100 | 100 per class | |  |  |  |
| G_XXX |  | |  |  | All from the same genus. |

**Table S2. Model details mined from Hugging Face Hub. CNN, Convolutional Neural Network; ViT, Vision Transformer.**

| Name | Model card | Params (M) | GMACs | Activations (M) | Image train size | Architecture | Pretraining method |
| --- | --- | --- | --- | --- | --- | --- | --- |
| ResNet-50 | resnet50.a1_in1k | 25.6 | 4.1 | 11.1 | 224 | CNN | Supervised |
| ConvNeXt v2-F | convnextv2_femto.fcmae_ft_in1k | 5.2 | 0.8 | 4.6 | 224 | CNN | Supervised |
| ConvNeXt v2-N | convnextv2_nano.fcmae_ft_in22k_in1k | 15.6 | 2.5 | 8.4 | 224 | CNN | Supervised |
| ConvNeXt v2-B | convnextv2_base.fcmae_ft_in22k_in1k | 88.7 | 15.4 | 28.8 | 224 | CNN | Supervised |
| ConvNeXt v2-L | convnextv2_large.fcmae_ft_in22k_in1k | 198.0 | 101.1 | 126.7 | 384 | CNN | Supervised |
| RegNetY-400MF | regnety_004.tv2_in1k | 4.3 | 0.4 | 3.9 | 224 | CNN | Supervised |
| RegNetY-4GF | regnety_040.ra3_in1k | 20.6 | 4.0 | 12.3 | 224 | CNN | Supervised |
| RegNetY-16GF | regnety_160.sw_in12k_ft_in1k | 83.6 | 16.0 | 23.0 | 224 | CNN | Supervised |
| ViT-T | vit_tiny_patch16_224.augreg_in21k_ft_in1k | 5.7 | 1.1 | 4.1 | 224 | ViT | Supervised |
| ViT-S | vit_small_patch16_224.augreg_in21k_ft_in1k | 22.1 | 4.3 | 8.2 | 224 | ViT | Supervised |
| ViT-B | vit_base_patch16_224.augreg2_in21k_ft_in1k | 86.6 | 16.9 | 16.5 | 224 | ViT | Supervised |
| EVA02-T | eva02_tiny_patch14_224.mim_in22k | 5.5 | 1.7 | 9.1 | 224 | ViT | Supervised |
| EVA02-S | eva02_small_patch14_224.mim_in22k | 21.6 | 6.1 | 18.3 | 224 | ViT | Supervised |
| EVA02-B | eva02_base_patch14_224.mim_in22k | 85.8 | 23.2 | 36.6 | 224 | ViT | Supervised |
| DINOv3-B | vit_base_patch16_dinov3.lvd1689m | 85.6 | 23.6 | 34.1 | 256 | ViT | Self-supervised |

**Table S3. Default hyperparameters for self-supervised pretraining.**

| Category | Hyperparameter | Default | Notes |
| --- | --- | --- | --- |
| Data | Global crop size | 224 | Aligned to patch size when needed |
| Data | Local crop size | 96 | Aligned to patch size when needed |
| Data | Number of local crops | 6 | Two global crops are always used |
| Objective | Mask ratio (ρ) | 0.5 | Fraction of patch tokens used in masked token loss |
| Objective | λlocal | 1.5 | Weight for local→global distillation |
| Objective | λmask | 1.0 | Weight for masked token regression |
| Teacher | EMA momentum (start) | 0.995 | Cosine schedule |
| Teacher | EMA momentum (end) | 0.999 | Cosine schedule |
| Temperature | Student temperature (τs) | 0.1 | Constant |
| Temperature | Teacher temperature (start) | 0.04 | Linear warmup |
| Temperature | Teacher temperature (end) | 0.07 | Warmup for first 70% of iterations |
| Optimizer | Optimizer | AdamW | Student parameters |
| Optimizer | Learning rate | 5e-4 | Cosine LR schedule |
| Optimizer | Weight decay | 0.05 |  |
| Training | Pretrain epochs | 50 |  |
| Training | Warmup epochs | 3 | LR warmup |
| Stabilization | Gradient clipping | 3.0 |  |

**Table S4. Default augmentation for global and local crops.**

| Component | Global crops | Local crops |
| --- | --- | --- |
| RandomResizedCrop scale | (0.4, 1.0) | (0.05, 0.4) |
| RandomRotation degrees | 180 | 180 |
| HorizontalFlip p | 0.5 | 0.5 |
| VerticalFlip p | 0.5 | 0.5 |
| ColorJitter | (0.4, 0.4, 0.4, 0.1) | (0.4, 0.4, 0.4, 0.1) |
| RandomGrayscale p | 0.2 | 0.2 |
| GaussianBlur | kernel = 23  sigma = (0.1, 2.0) | kernel = 7  sigma = (0.1, 2.0) |
| Normalization | ImageNet mean/std | ImageNet mean/std |

**Table S5. Default hyperparameters for ArcFace finetuning.**

| Category | Hyperparameter | Default | Notes |
| --- | --- | --- | --- |
| Finetune | Finetune epochs | 20 |  |
| Finetune | Finetune learning rate | 1e-4 | AdamW |
| Embedding | Metric embedding dim | 256 |  |
| Regularization | Frozen transformer blocks | first 70% | Freeze early blocks |
| ArcFace | Scale (s) | 64 |  |
| ArcFace | Margin (m) | 28.6 | Value depends on implementation convention |

### **Supplementary Figures**

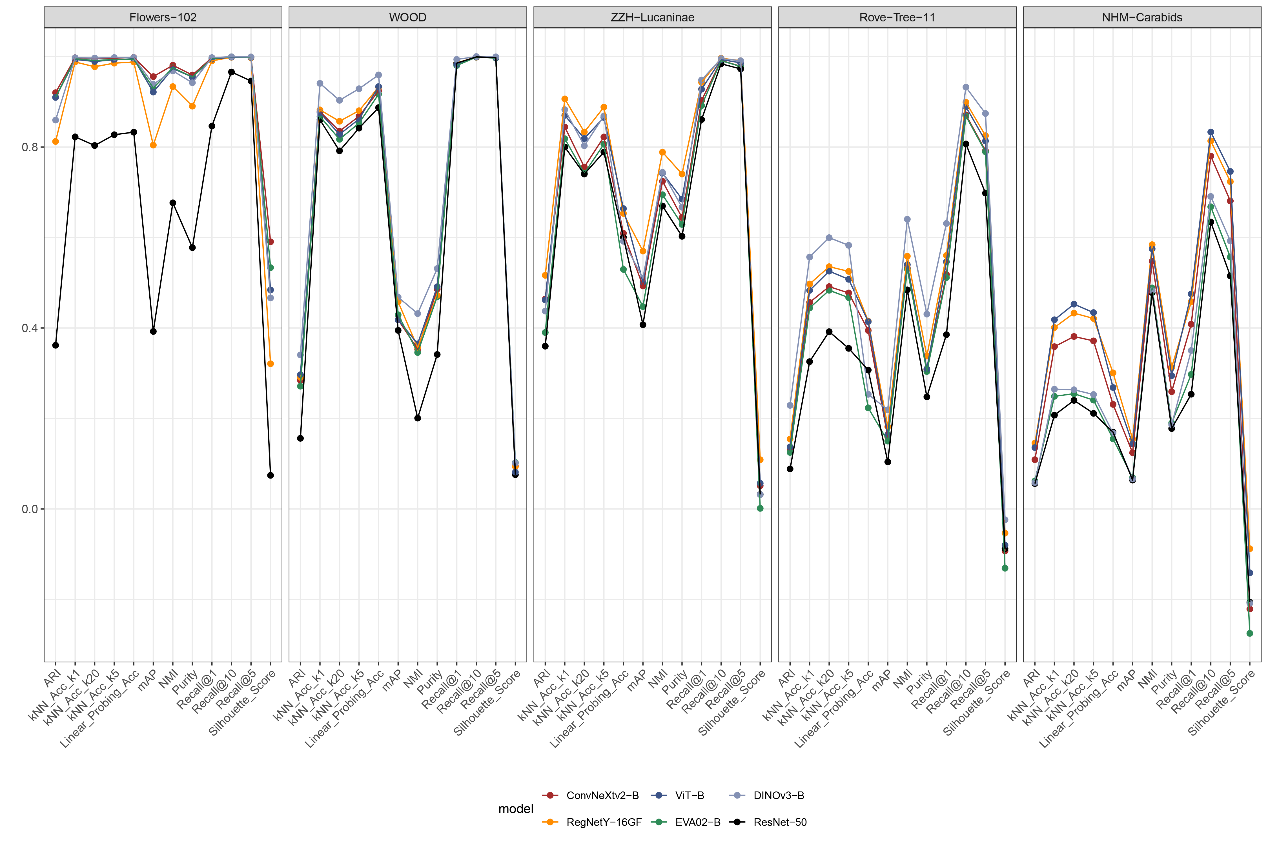

**Figure S1 | Baseline embedding performance of ImageNet‑scale pretrained backbones across all five datasets.**

Evaluation of six commonly used CNN and Transformer architectures without task‑specific training: ConvNeXtV2‑B, ViT‑B, DINOv3‑B, RegNetY‑16GF, EVA02‑B, and ResNet‑50. Models were assessed using a suite of classification, retrieval, and clustering metrics, including ARI, k‑NN accuracy (k = 1, 5), linear probing accuracy, mAP, purity, NMI, recall@10, and silhouette score. Panels correspond to the Flowers‑102, WOOD, ZZH–Lucaninae, Rove‑Tree‑11, and NHM‑Carabids datasets. Pretrained backbones show moderate but variable embedding quality depending on dataset difficulty, with consistent underperformance of ResNet‑50 relative to modern CNN and Transformer architectures.

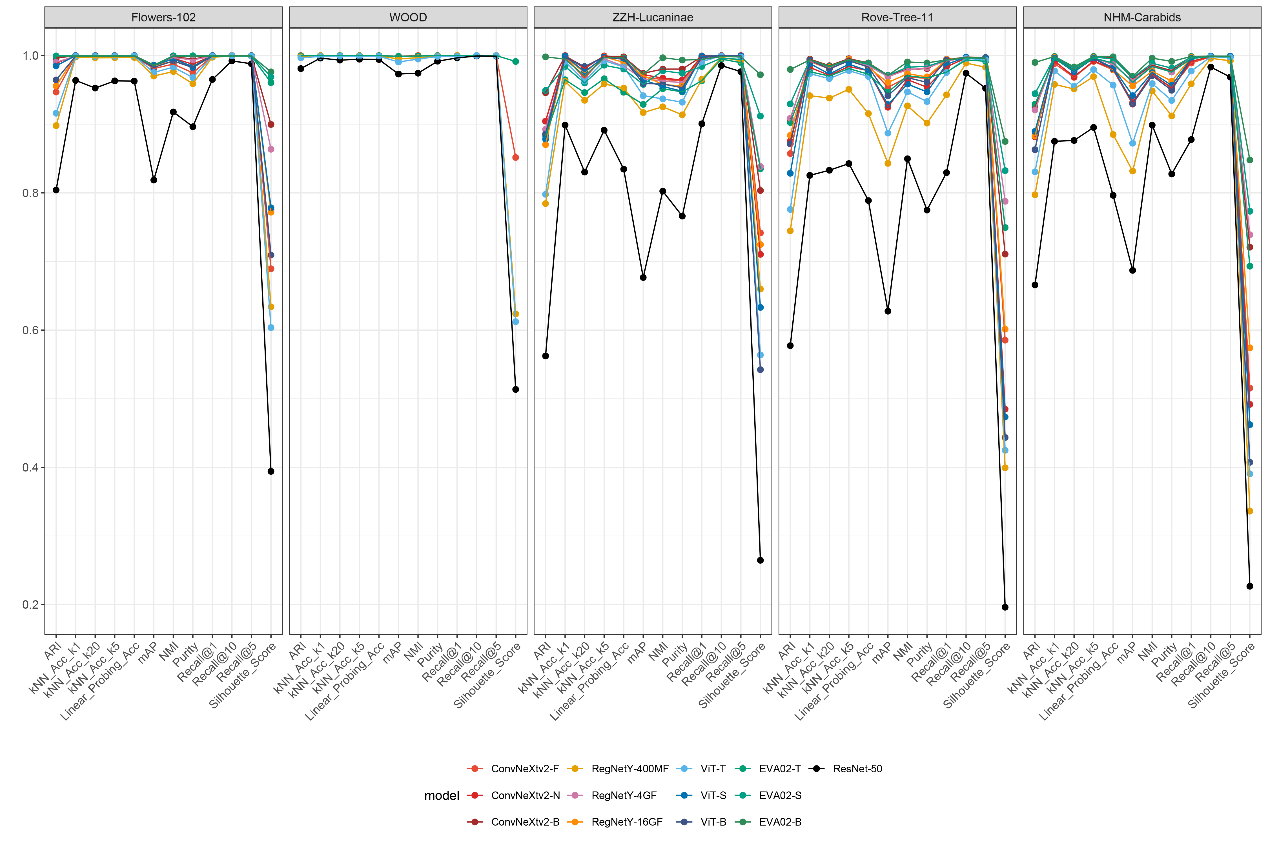

**Figure S2 | Embedding performance of supervised fine‑tuned backbone architectures across all five datasets.**

ConvNeXtV2 (F/N/B), RegNetY (400MF/4GF/16GF), ViT (T/S/B), EVA02 (T/S/B), and ResNet‑50 were fine‑tuned using AutoGluon‑Multimodal and evaluated on Flowers‑102, WOOD, ZZH–Lucaninae, Rove‑Tree‑11, and NHM‑Carabids. Metrics include ARI, k‑NN accuracy (k = 1, 5), linear‑probing accuracy, mAP, purity, NMI, recall@10, and silhouette score.

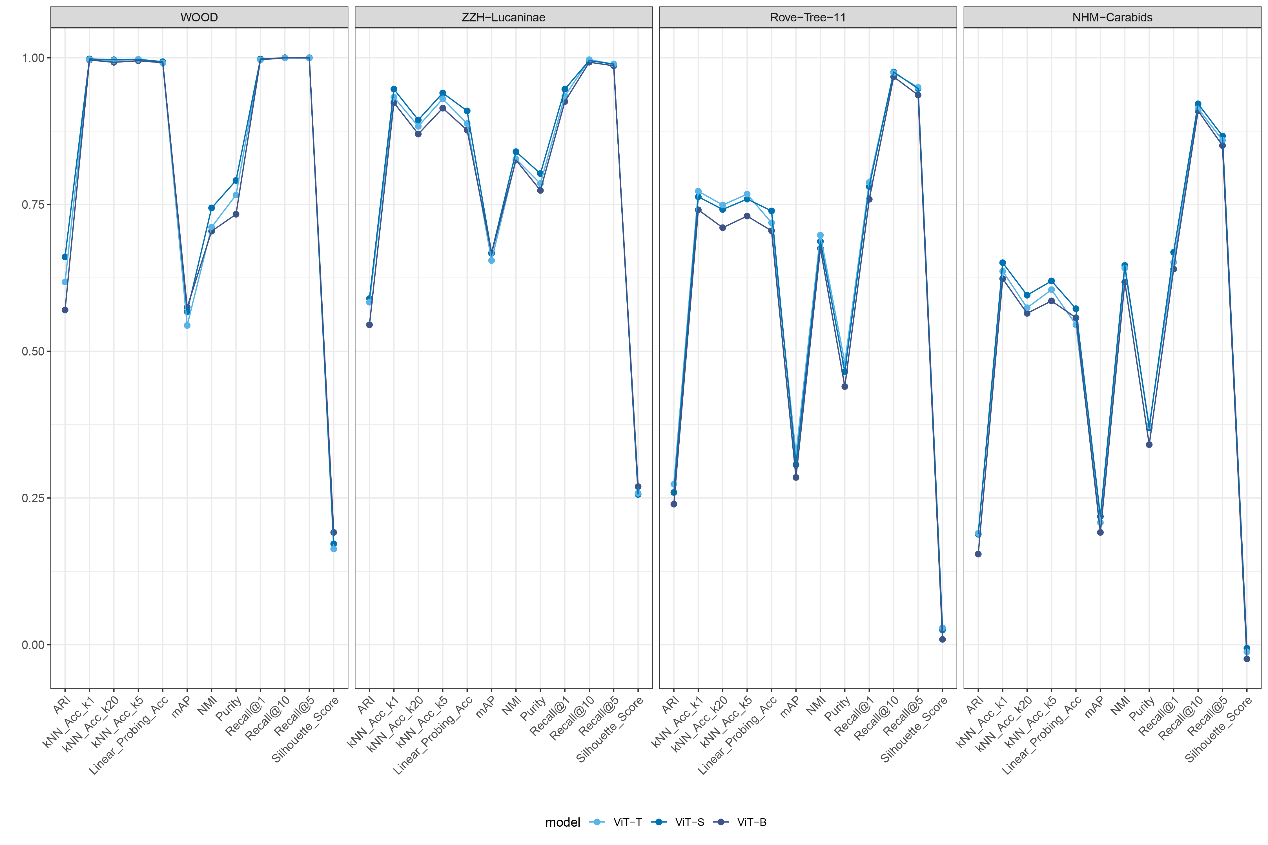

**Figure S3 | Embedding performance of self‑supervised ViT encoders across four datasets.**

Self‑supervised models (ViT‑T, ViT‑S, ViT‑B) were trained using the OTU‑Former SSL framework and evaluated on WOOD, ZZH–Lucaninae, Rove‑Tree‑11, and NHM‑Carabids. Metrics include ARI, k‑NN accuracy (k = 1, 5), linear‑probing accuracy, mAP, purity, NMI, recall@10, and silhouette score. Performance varies across datasets, with WOOD and ZZH–Lucaninae showing the strongest SSL gains and NHM–Carabids exhibiting the largest variability, consistent with its lower image quality. These results highlight the effectiveness of self‑supervised learning for extracting organism‑centred morphological features even without labeled data.

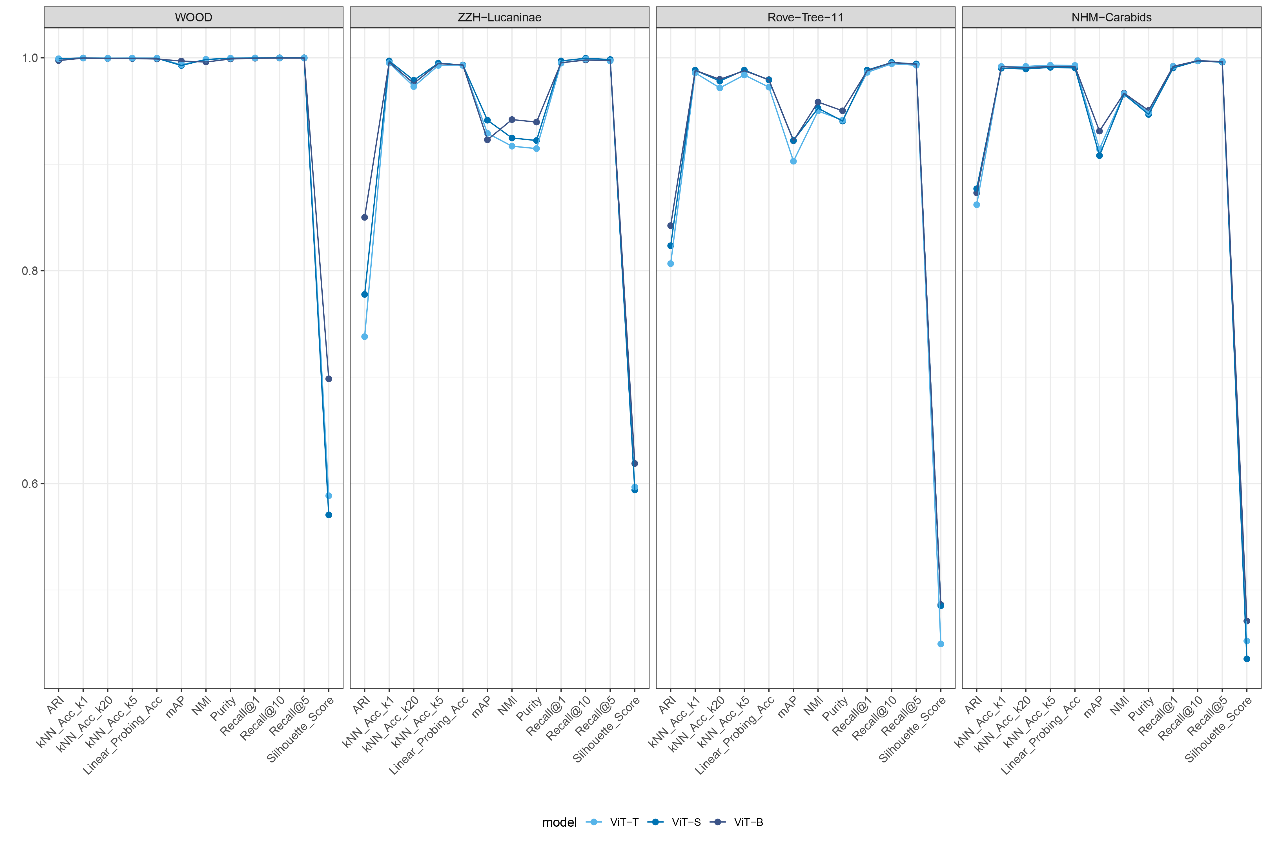

**Figure S4 | Embedding performance of ViT encoders after self‑supervised pretraining followed by ArcFace supervised fine‑tuning (SSL+SFT).**

ViT‑T, ViT‑S, and ViT‑B backbones were pretrained using the OTU‑Former SSL framework and subsequently fine‑tuned with ArcFace metric learning. Embedding quality was evaluated on WOOD, ZZH–Lucaninae, Rove‑Tree‑11, and NHM‑Carabids using ARI, k‑NN accuracy (k = 1, 5), linear‑probing accuracy, mAP, purity, NMI, recall@10, and silhouette score. The NHM–Carabids dataset shows the largest variation due to lower image quality, yet SSL+SFT still yields stable and high‑quality embeddings suitable for downstream morphOTU delineation.

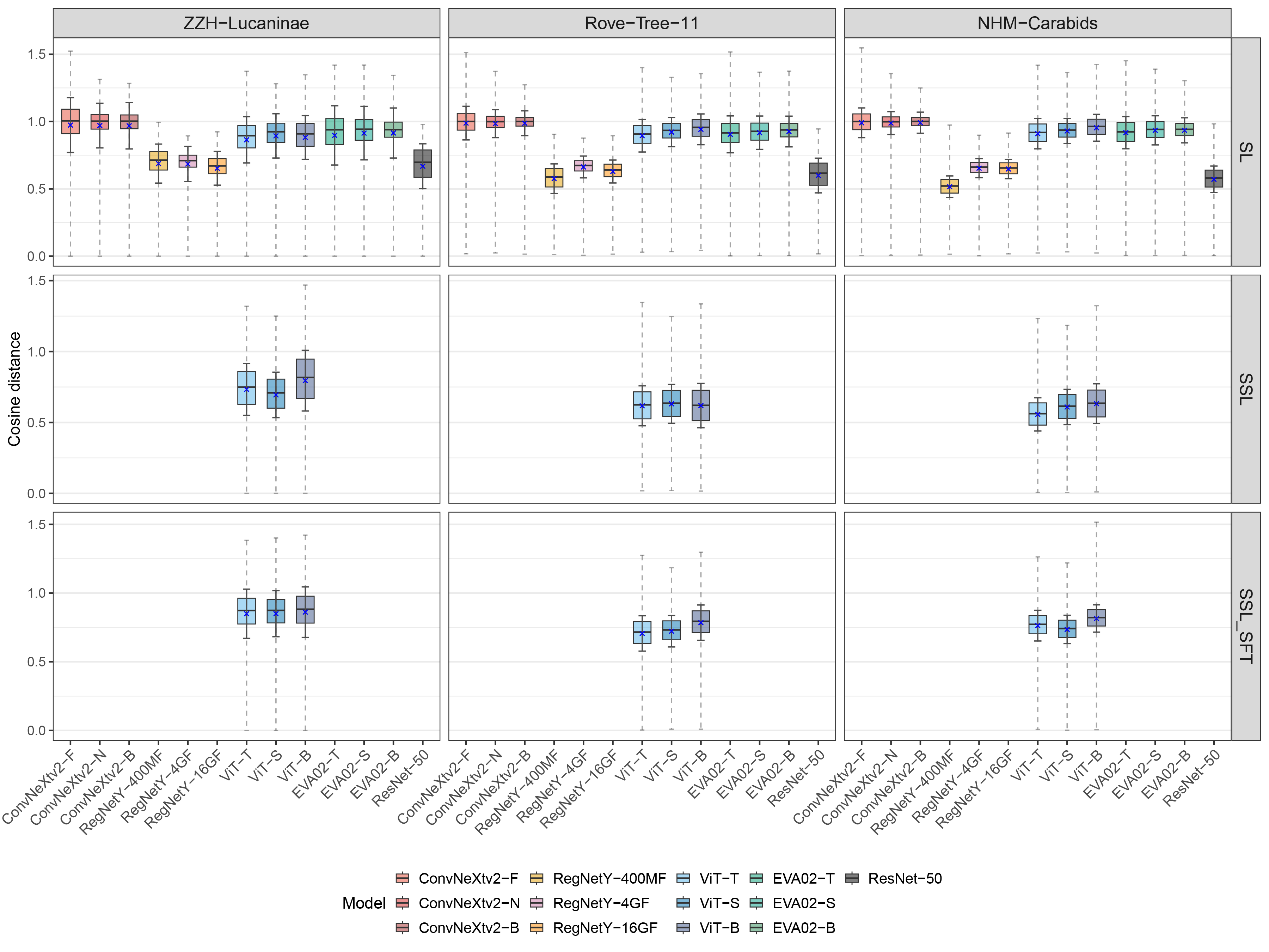

**Figure S5 | Pairwise cosine distance distributions for embeddings produced by supervised learning (SL), self‑supervised learning (SSL), and SSL followed by ArcFace fine‑tuning (SSL+SFT).**

Boxplots show the distribution of all pairwise cosine distances for each backbone architecture under three training paradigms across the ZZH–Lucaninae, Rove‑Tree‑11, and NHM–Carabids datasets. Models include ConvNeXtV2 (F/N/B), RegNetY (400MF/4GF/16GF), ViT (T/S/B), EVA02 (T/S/B), and ResNet‑50.

Supervised models generally exhibit wider and more separated distance ranges than SSL‑only encoders, reflecting the explicit enlargement of interclass margins during training. SSL embeddings show more compact distance distributions, indicating tighter intra‑class structure but reduced inter‑class separation. After ArcFace fine‑tuning (SSL+SFT), distance distributions expand and become more distinct, closely resembling or exceeding supervised baselines. These patterns highlight how different training objectives shape embedding geometry and influence downstream morphOTU formation.

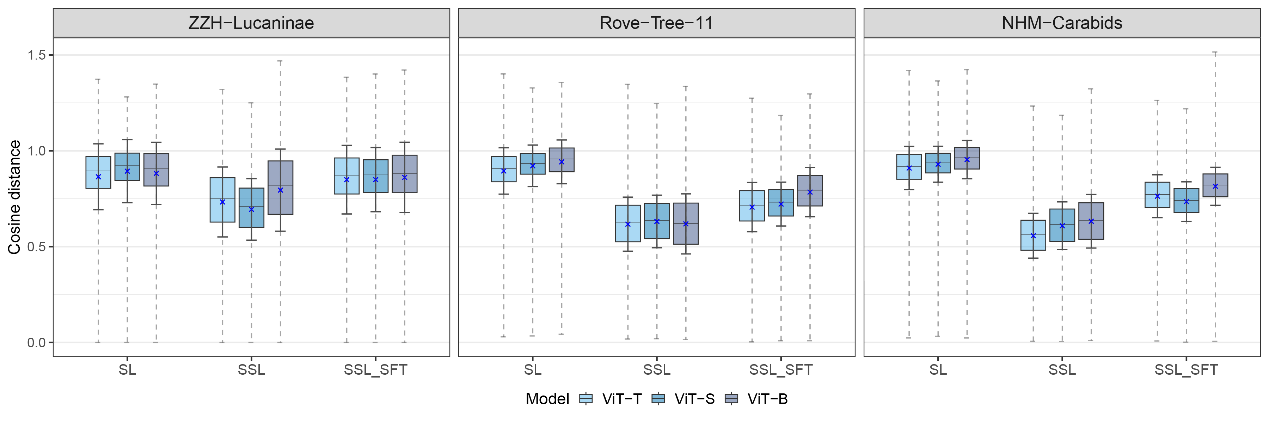

**Figure S6 | Intra‑class cosine distance distributions for ViT‑T, ViT‑S, and ViT‑B encoders across supervised learning (SL), self‑supervised learning (SSL), and SSL followed by ArcFace fine‑tuning (SSL+SFT).**

Boxplots show the distribution of cosine distances among images belonging to the same species for the ZZH–Lucaninae, Rove‑Tree‑11, and NHM‑Carabids datasets. Lower intra‑class distances indicate tighter and more coherent phenotypic clustering.

SSL models exhibit reduced intra‑class distances relative to SL, reflecting compact organism‑centred representations learned without labels. ArcFace fine‑tuning (SSL+SFT) slightly increases intra‑class distances—consistent with the enlargement of inter‑class margins—yet maintains cohesive clusters with minimal dispersion. Together, these results demonstrate that SSL provides strong within‑class structure and that metric learning preserves class cohesion while enhancing separability for downstream morphOTU delineation.

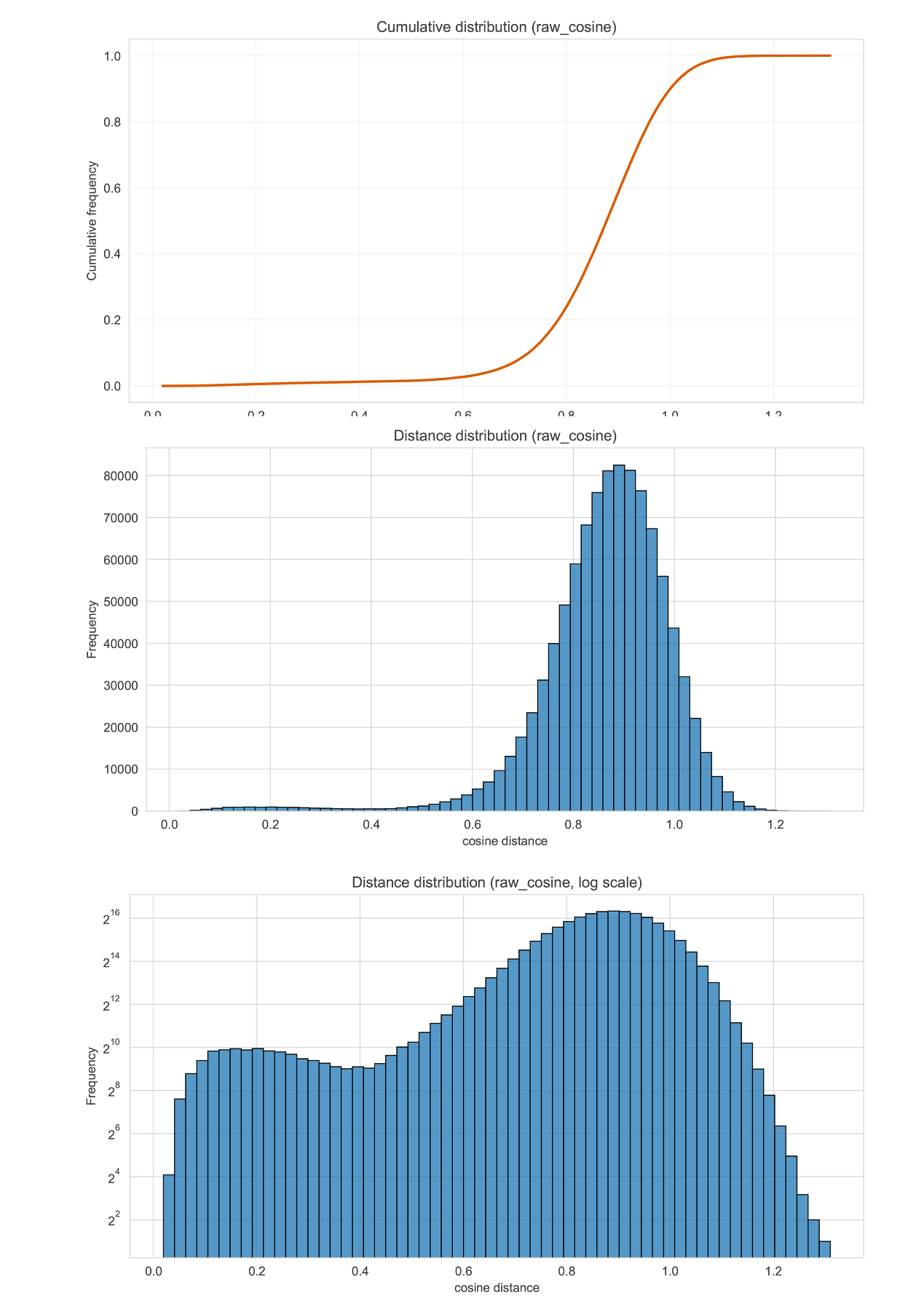

**Figure S7 | Distributional properties of raw cosine distances among all image pairs in the embedding space of the Flower dataset under the SL model.**

The top panel shows the cumulative distribution function (CDF) of raw cosine distances, the middle panel shows the corresponding histogram, and the bottom panel displays the same histogram on a log‑scaled frequency axis to emphasize the tail structure. Distances were computed from SSL+SFT embeddings on a representative dataset.

Cosine distances form a broad, unimodal distribution without discrete separation between intra‑ and inter‑class distances, consistent with the absence of a morphological “barcoding gap”. The log‑scaled histogram highlights the long, continuous tail of rare but large distances. These patterns underscore the necessity of lineage‑specific and dynamic thresholds for morphOTU delineation rather than fixed global cutoffs.

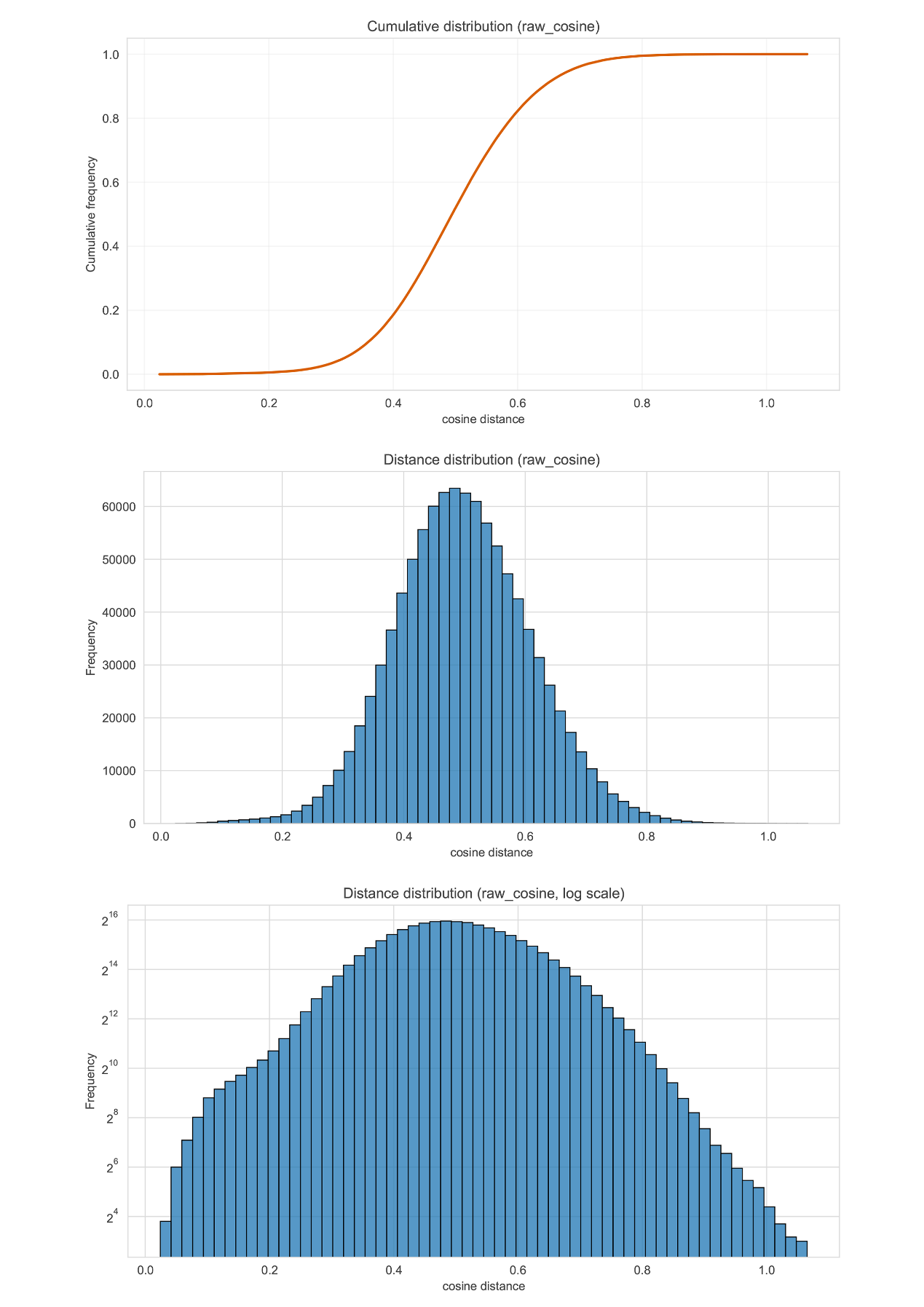

**Figure S8 | Distributional properties of raw cosine distances among images belonging to the Flower dataset under the SSL+SFT model.**

The top panel shows the cumulative distribution function (CDF) of intra‑class cosine distances, the middle panel shows the corresponding histogram, and the bottom panel displays the histogram on a log‑scaled frequency axis to highlight rare small‑distance events. Distances were computed from SSL+SFT embeddings on a representative dataset.

Intra‑class distances form a narrow, approximately Gaussian distribution centred around moderate similarity values, reflecting consistent morphological clustering within species. The log‑scaled histogram reveals very few extremely small distances, indicating that even within species, individuals occupy a distributed region of phenotype space rather than collapsing into a point. These results support the view that morphOTUs should account for realistic within‑species variance and avoid overly strict similarity thresholds.

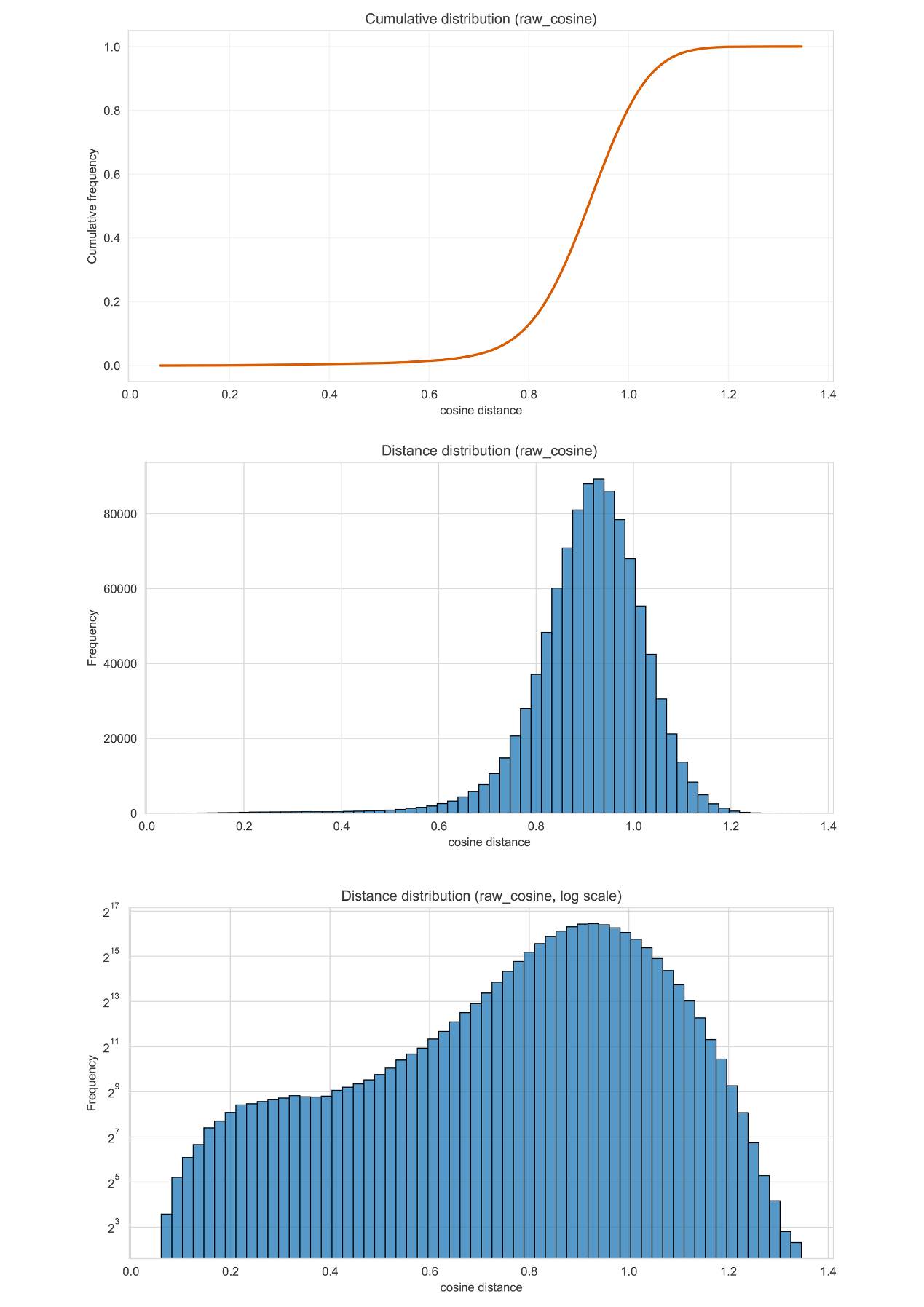

**Figure S9 | Distributional properties of raw cosine distances among images of NHM‑Carabids under the SL model.**

The top panel shows the cumulative distribution function (CDF) of inter‑class cosine distances, the middle panel shows the corresponding histogram, and the bottom panel presents the same histogram on a log‑scaled frequency axis to highlight the long-distance tail. Distances were computed from SSL+SFT embeddings on a representative dataset.

Inter‑class distances exhibit a broad, right‑shifted distribution relative to intra‑class distances, with most values concentrated between 0.85 and 1.05, and a heavy tail extending beyond 1.2. The log‑scaled histogram emphasizes the substantial heterogeneity among species‑level morphological differences. The continuous nature of this distribution, without multimodality or discrete separation from intra‑class distances, further supports the absence of a universal “barcoding gap” in morphological embedding space.

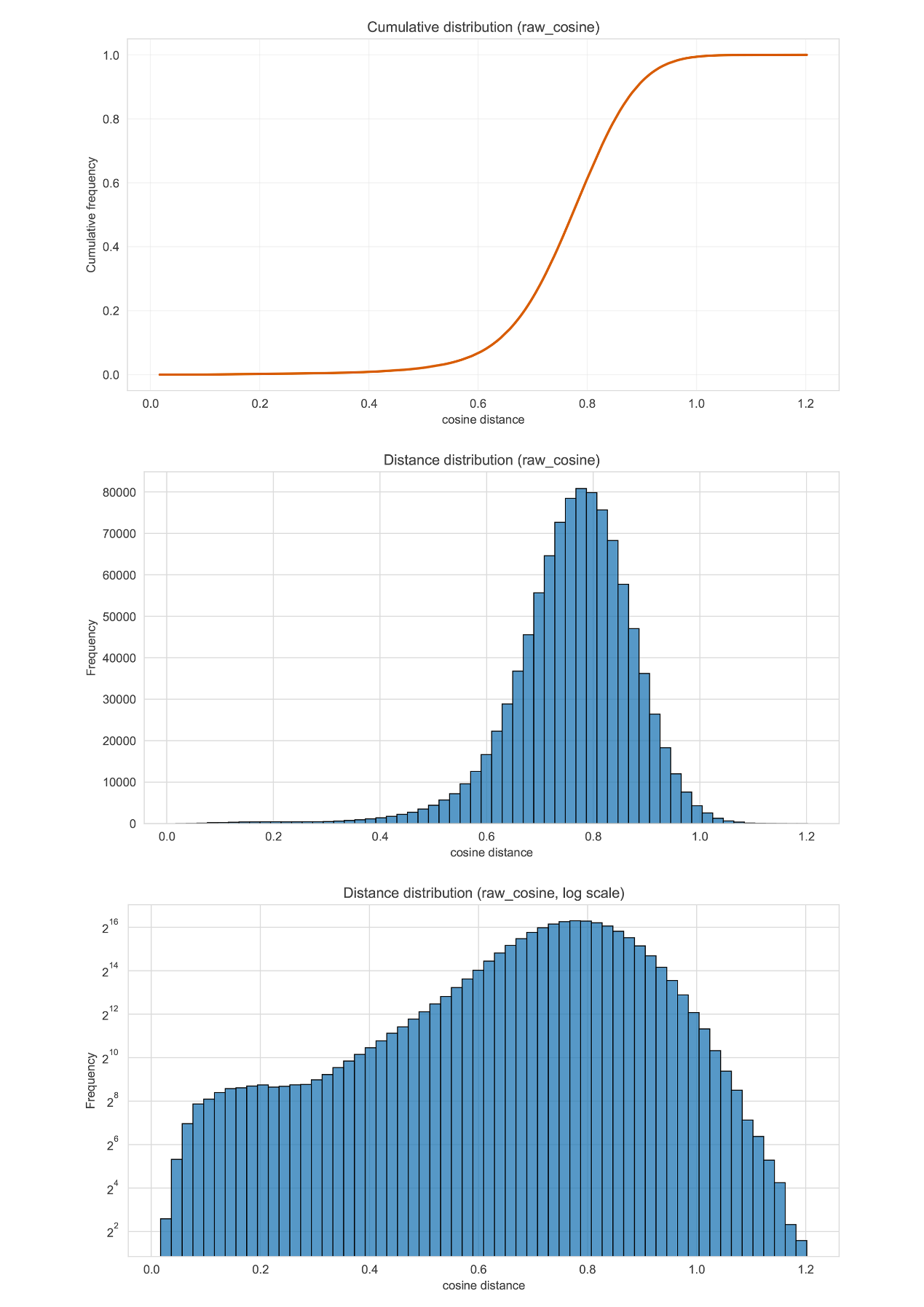

**Figure S10 | Distributional characteristics of raw cosine distances computed from image embeddings of NHM‑Carabids under the SLL+SFT model.**

The top panel shows the cumulative distribution function (CDF) of all pairwise cosine distances, the middle panel presents the corresponding histogram, and the bottom panel displays the same histogram on a log‑scaled frequency axis to highlight low‑frequency distance ranges.

The distribution spans a continuous range of cosine distances, with most pairs concentrated around intermediate‑to‑high values and a long, gradually decreasing tail. The log‑scaled view reveals the full extent of rare small‑ and large‑distance events that are not visible on a linear scale. Together, these panels illustrate the broad and smoothly varying structure of distances in the embedding space.

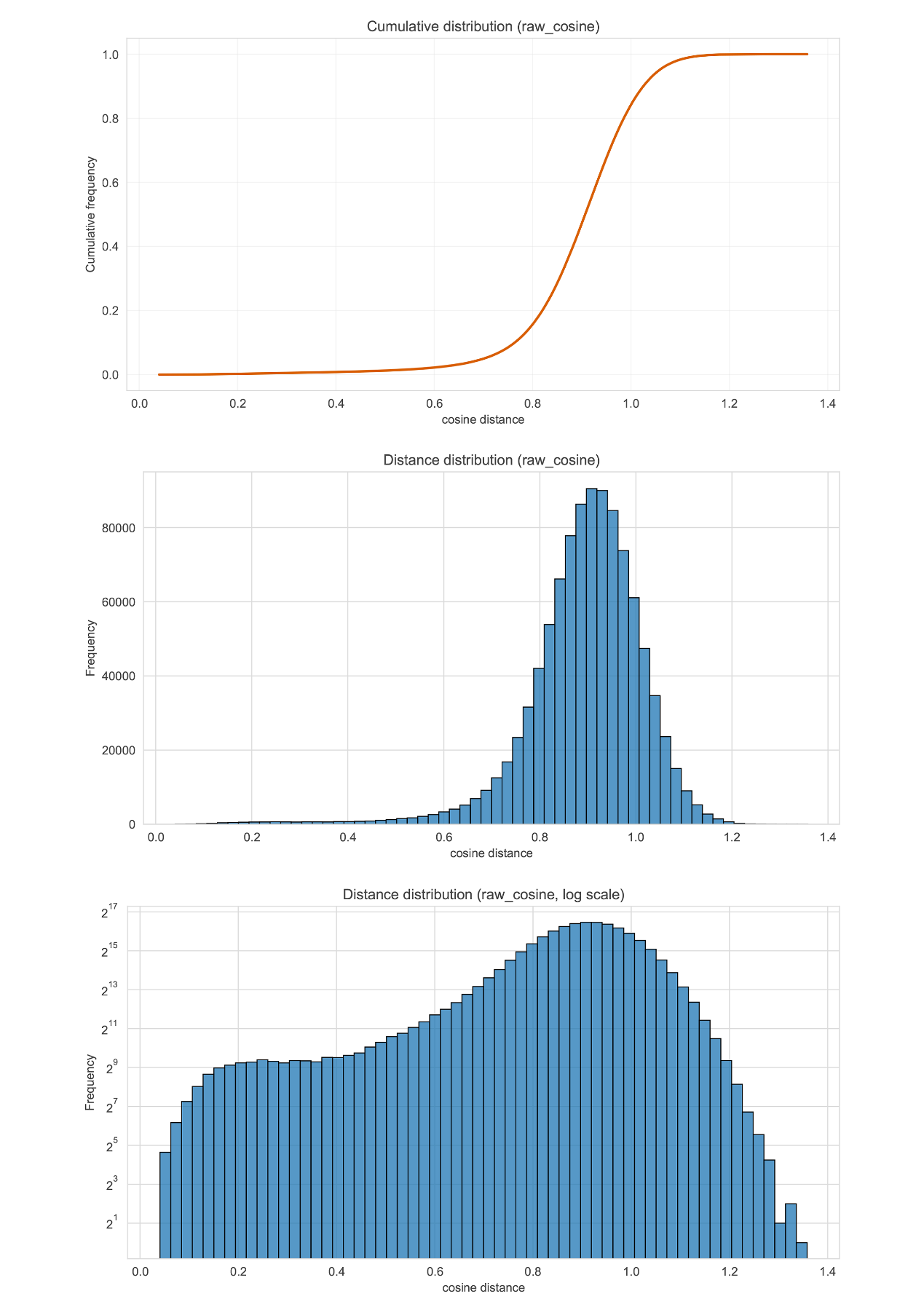

**Figure S11 | Raw cosine‑distance distributions derived from all pairwise comparisons in the embedding space of Rove-Tree-11 under the SL model.**

The top panel presents the cumulative distribution function (CDF) of pairwise cosine distances, the middle panel shows the corresponding histogram on a linear frequency scale, and the bottom panel displays the same histogram using a logarithmic frequency scale to reveal low‑probability regions.

The distances span a broad continuous range, with a prominent concentration near high‑similarity values and a gradually declining tail extending toward larger distances. The log‑scaled histogram highlights the presence of rare but informative extreme‑distance events that are not visible on a linear scale. These views collectively illustrate the smooth, non‑discrete structure of embedding‑space distances within the dataset.

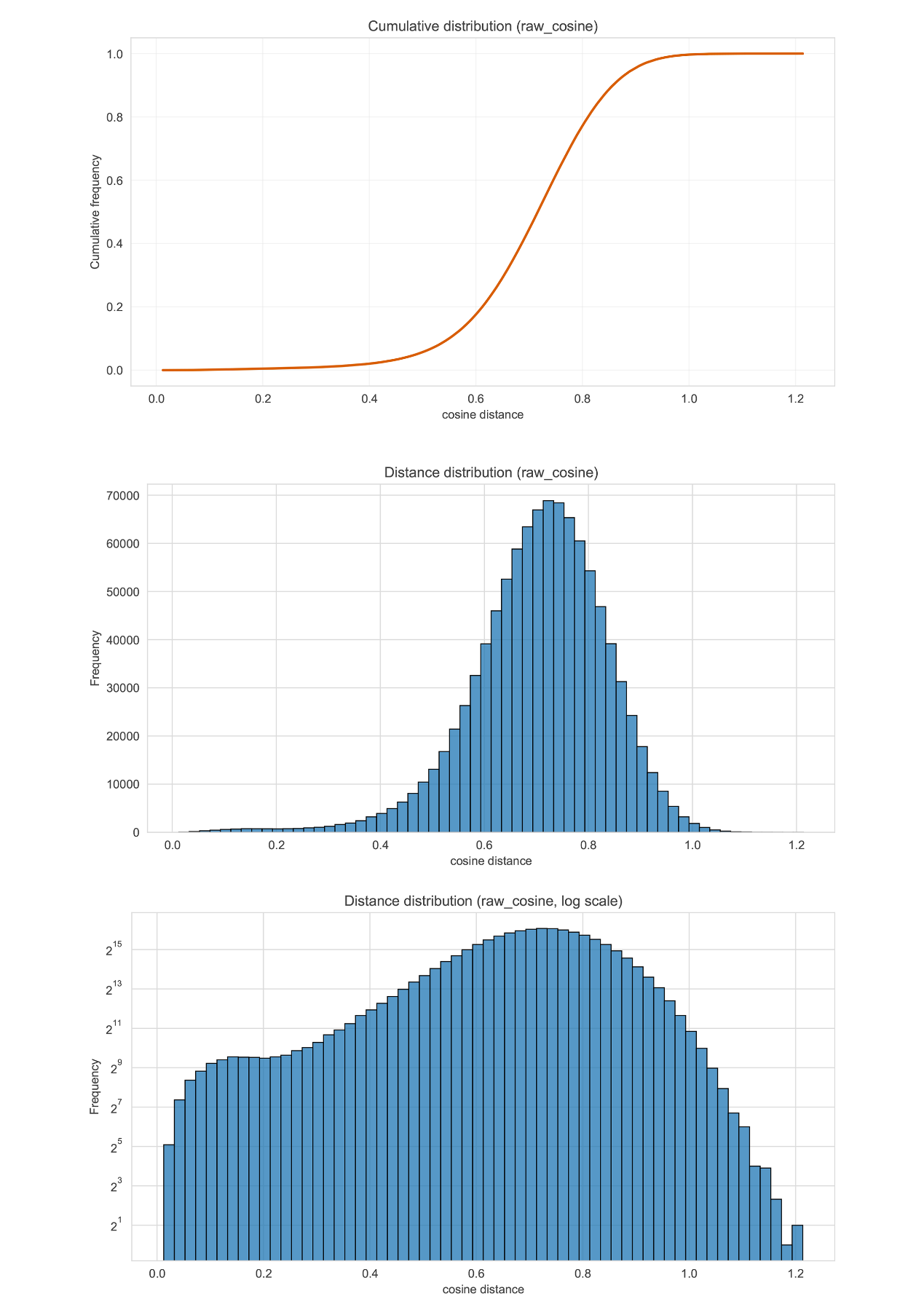

**Figure S12 | Raw cosine‑distance distributions computed from all pairwise comparisons in the embedding space of Rove-Tree-11 under the SLL+SFT model.**

The top panel displays the cumulative distribution function (CDF) of cosine distances, the middle panel shows the corresponding histogram on a linear frequency scale, and the bottom panel presents the same histogram using a logarithmic frequency axis to reveal rare distance values.

The distribution covers a continuous range of distances, with a prominent central region and smoothly decreasing tails toward both lower and higher similarity extremes. The log‑scaled histogram highlights the fine structure of low‑frequency events that are not apparent on a linear scale. These complementary views outline the overall geometry of pairwise distances produced by the embedding model.

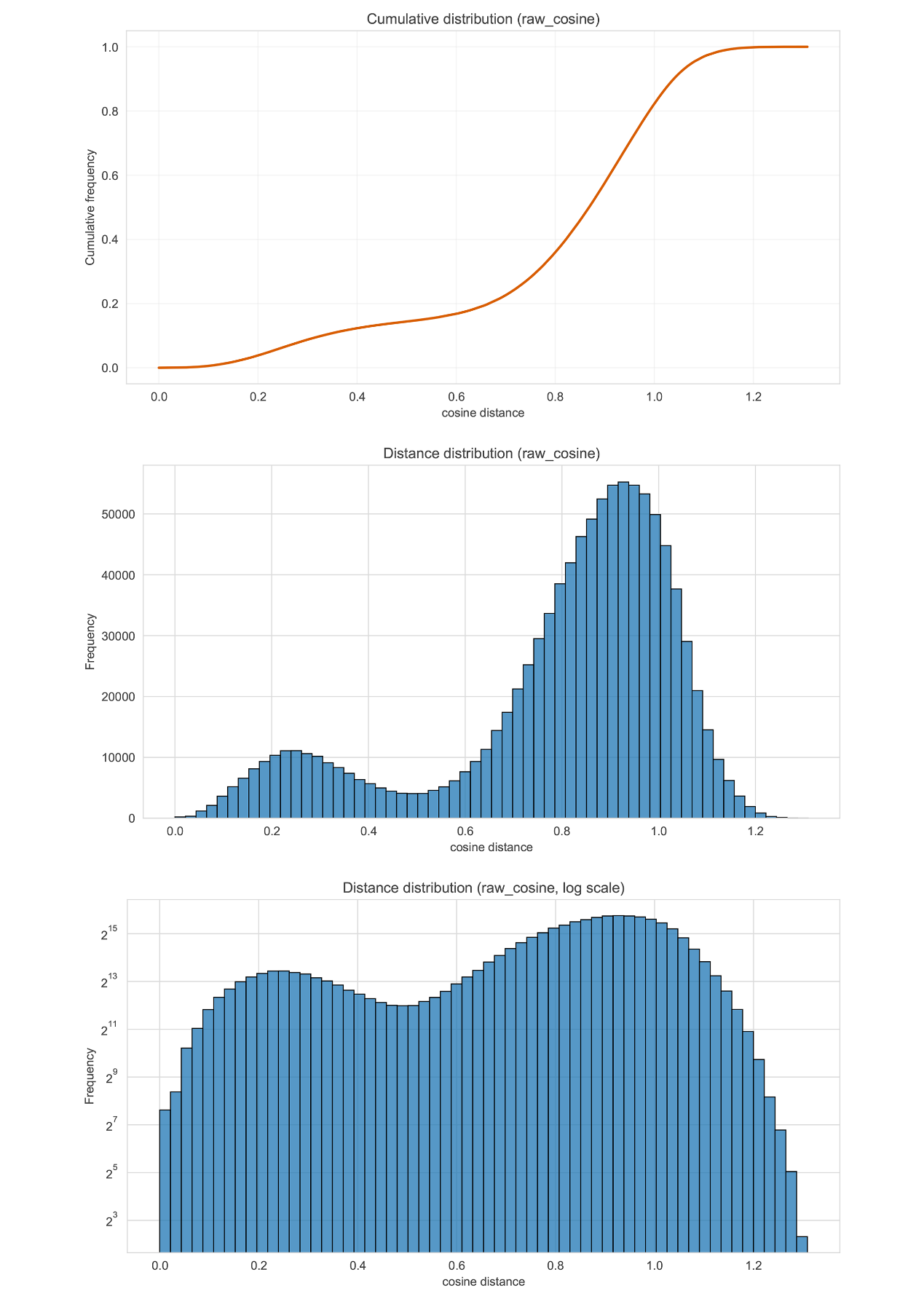

**Figure S13 | Raw cosine‑distance distributions computed from all pairwise image comparisons in the embedding space of WOOD under the SL model.**

The top panel shows the cumulative distribution function (CDF) of cosine distances, the middle panel presents the corresponding histogram on a linear frequency scale, and the bottom panel displays the same histogram using a logarithmic frequency axis to expose low‑frequency patterns.

The histogram reveals a pronounced multimodal structure, with multiple peaks indicating that image pairs cluster into several distinct similarity regimes. The log‑scaled representation highlights subtle troughs and secondary modes not apparent on a linear scale. These complementary views illustrate non‑uniform density features within the embedding space, suggesting heterogeneous relationships among samples in this dataset.

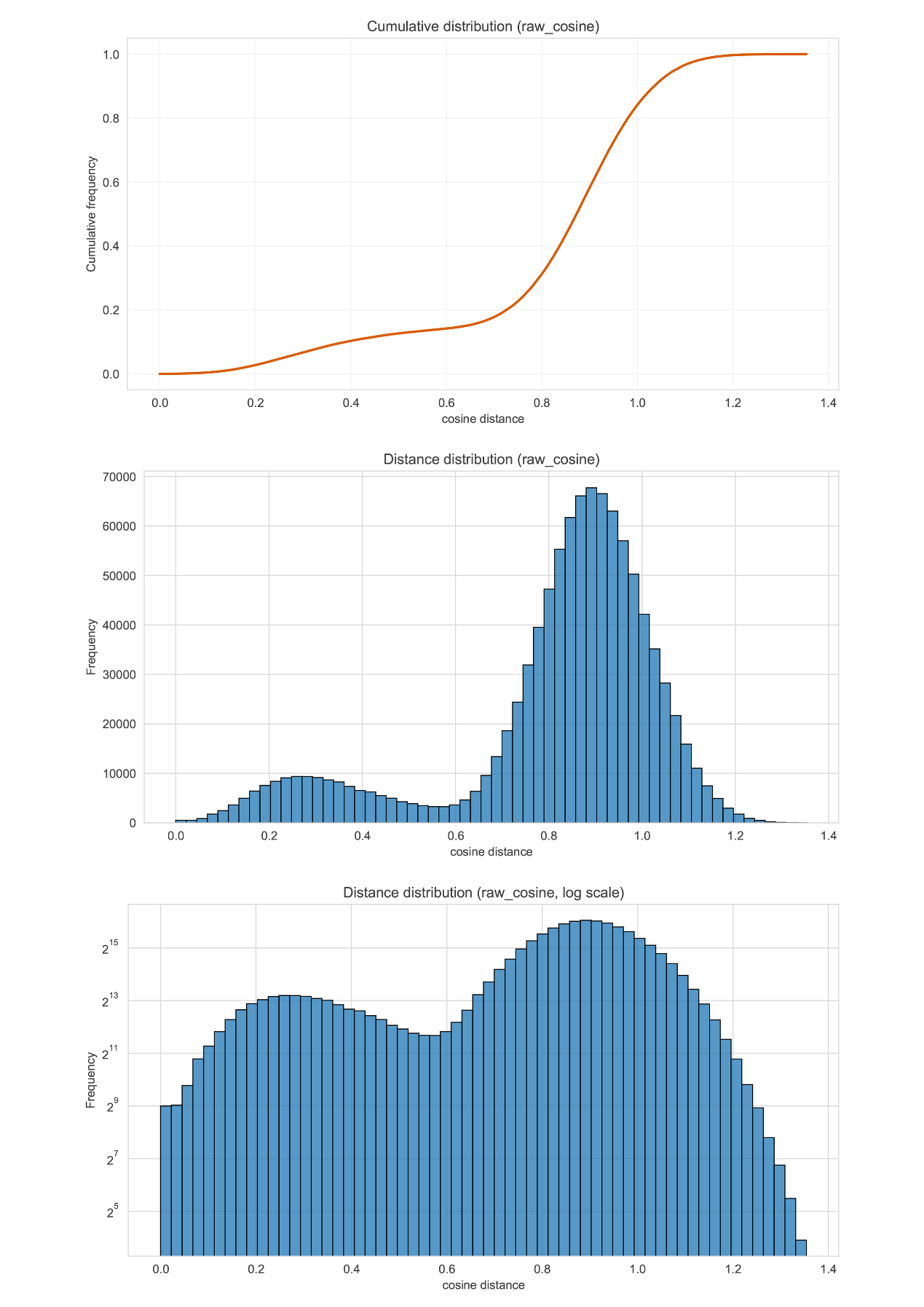

**Figure S14 | Raw cosine‑distance distributions derived from all pairwise image comparisons in the embedding space of WOOD under SLL+SFT model.**

The top panel shows the cumulative distribution function (CDF) of cosine distances, the middle panel presents the corresponding histogram on a linear frequency scale, and the bottom panel displays the same histogram with a logarithmic frequency axis to reveal low‑frequency features.

The histogram exhibits a clearly bimodal structure, with one cluster of image pairs concentrated at lower cosine distances and another forming a dominant peak at higher distances. The log‑scaled view highlights subtle valleys between these peaks as well as the extent of the distribution tails. Together, these panels indicate substantial heterogeneity within the embedding space, with samples forming multiple distinct similarity regimes.

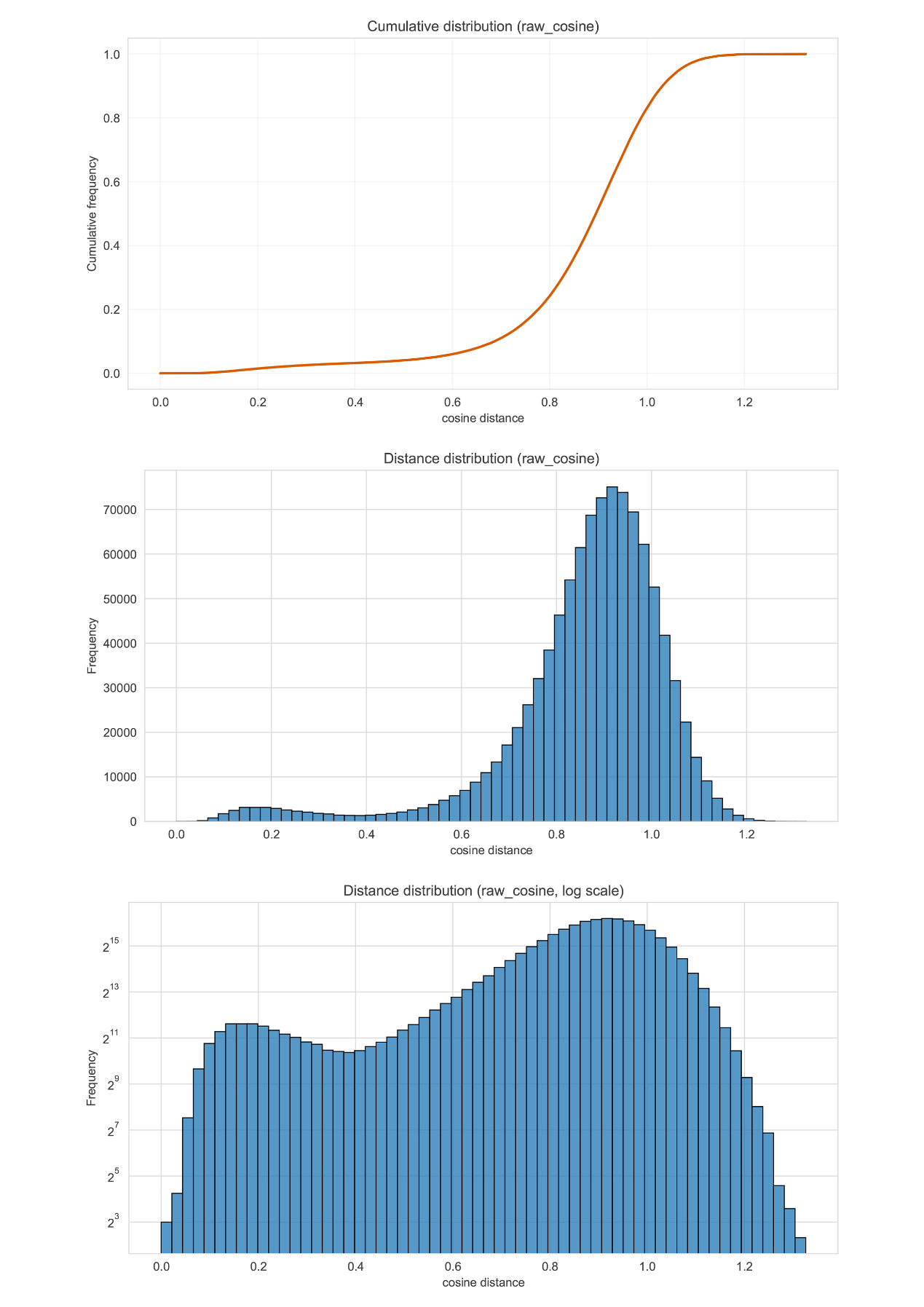

**Figure S15 | Raw cosine‑distance distributions computed from all pairwise image comparisons in the embedding space of ZHH-Lucaninae under the SL model.**

The top panel shows the cumulative distribution function (CDF) of cosine distances, the middle panel displays the corresponding histogram on a linear frequency scale, and the bottom panel presents the same histogram using a logarithmic frequency axis to highlight low‑frequency details.

The histogram reveals an asymmetric multimodal structure: a smaller peak at lower cosine distances is followed by a broad dominant peak at higher distances, separated by a shallow trough. The log‑scaled view emphasizes the shape of the distribution tails and the relative sparsity of intermediate‑distance pairs. These complementary representations illustrate complex, non‑uniform similarity relationships among samples within the embedding space.

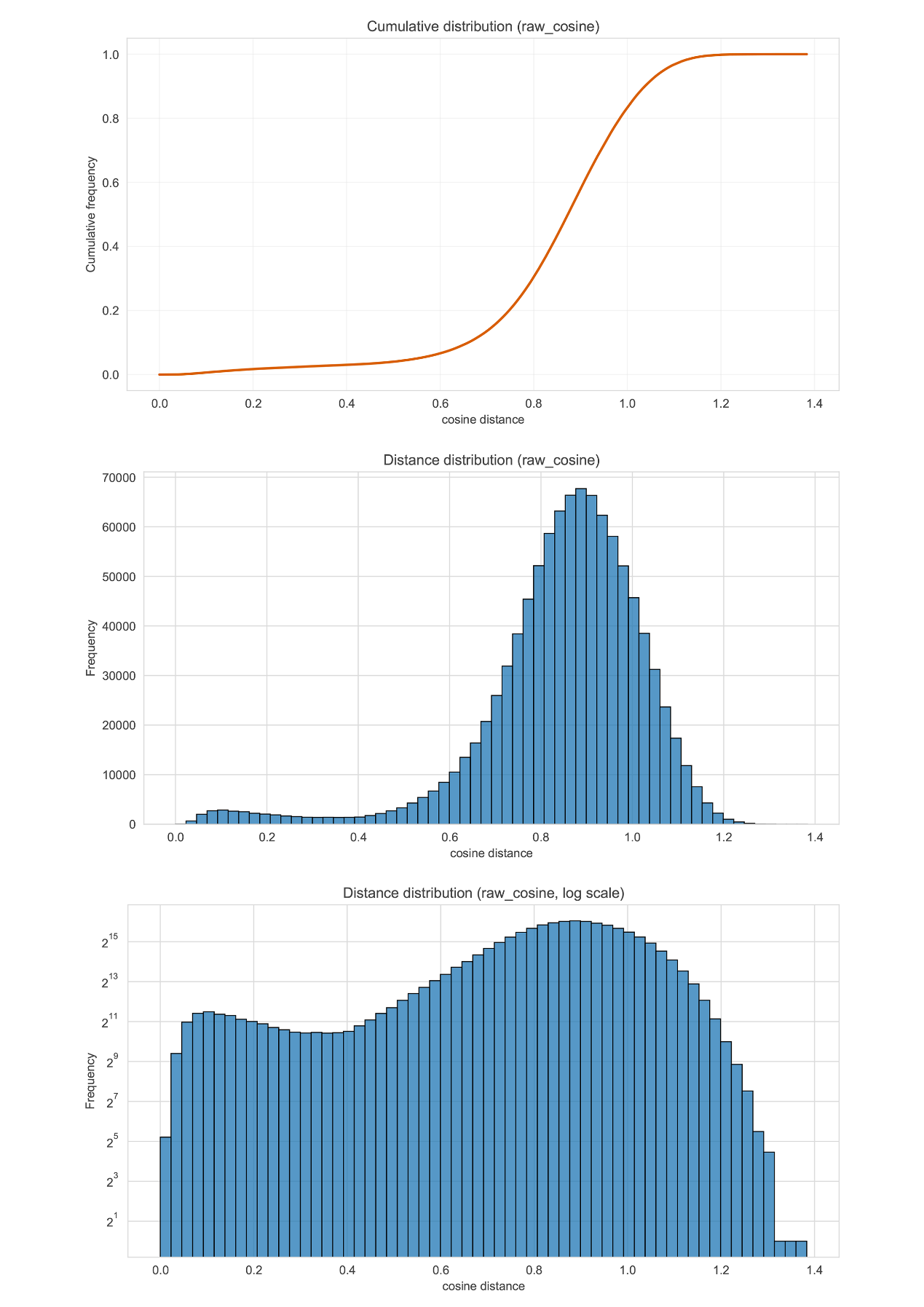

**Figure S16 | Raw cosine‑distance distributions computed from all pairwise image comparisons in the embedding space of ZHH-Lucaninae under the SLL+SFT model.**

The top panel shows the cumulative distribution function (CDF) of cosine distances, the middle panel presents the corresponding histogram on a linear frequency scale, and the bottom panel displays the same histogram with a logarithmic frequency axis to reveal features in the low‑frequency range.

The histogram exhibits a broad primary peak at higher cosine distances, accompanied by a secondary rise at lower distances and a long right‑skewed tail. The log‑scaled view emphasizes the gradual tapering of the distribution and clarifies the prevalence of rare, extreme‑distance events. These complementary representations capture the extended and heterogeneous structure of pairwise similarities within the embedding space.

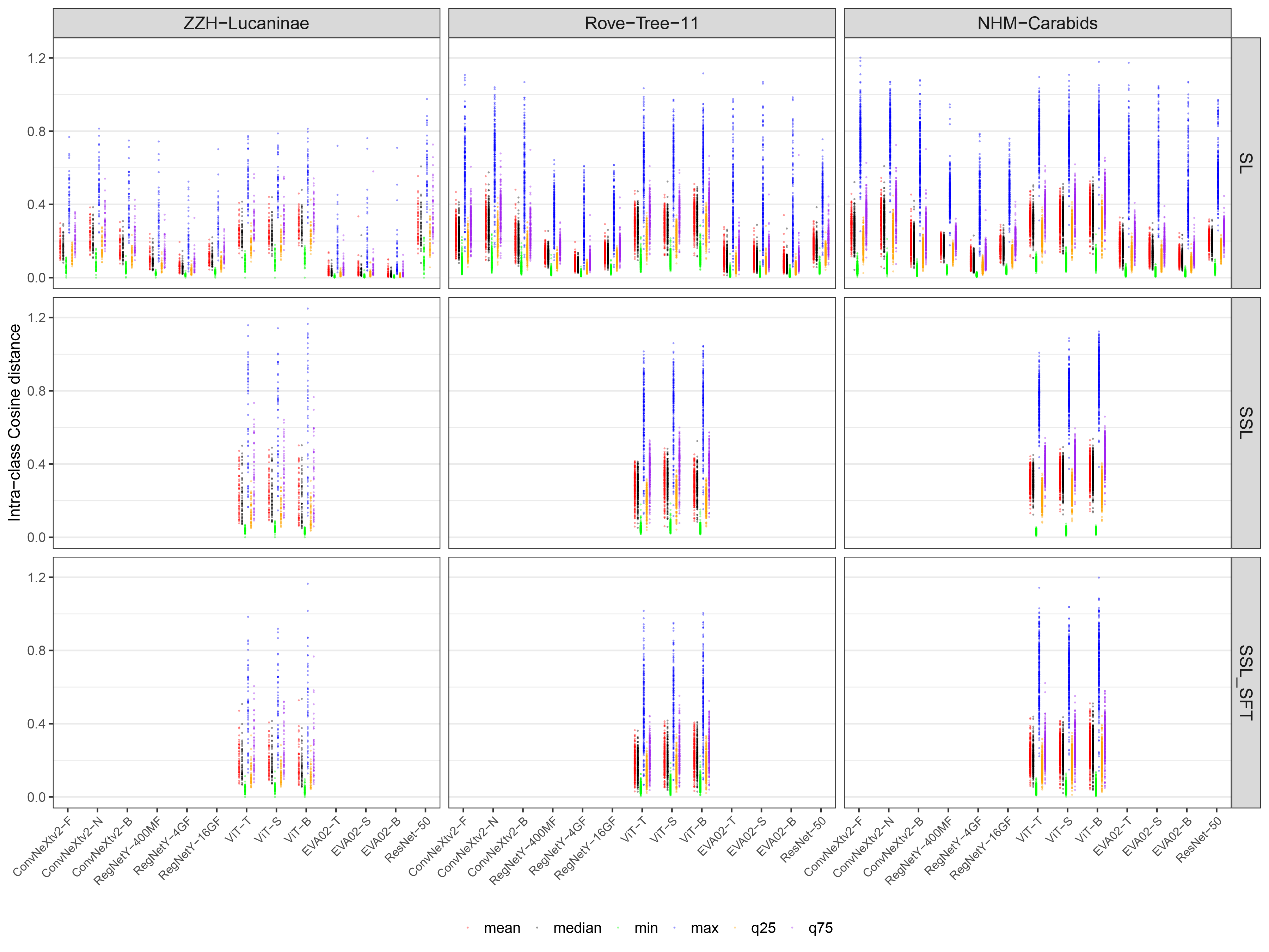

**Figure S17 | Summary statistics of intra‑class cosine distances across multiple model architectures and datasets under three training regimes.**

Each panel corresponds to one dataset (columns: ZZH–Lucaninae, Rove‑Tree‑11, NHM–Carabids) and one training condition (rows: SL, SSL, SSL+SFT). Within each panel, models are shown along the x‑axis, and for each model, intra‑class distances are summarized using five statistics: mean (red), median (black), minimum (green), maximum (blue), and the first and third quartiles (orange and brown).

The distributions reveal substantial variation in intra‑class compactness across model families and training regimes. Some models produce tightly clustered embeddings with low variance, while others yield broader or skewed intra‑class distance ranges. Differences among datasets illustrate how taxonomic composition and image characteristics influence embedding geometry under the same model and training conditions.

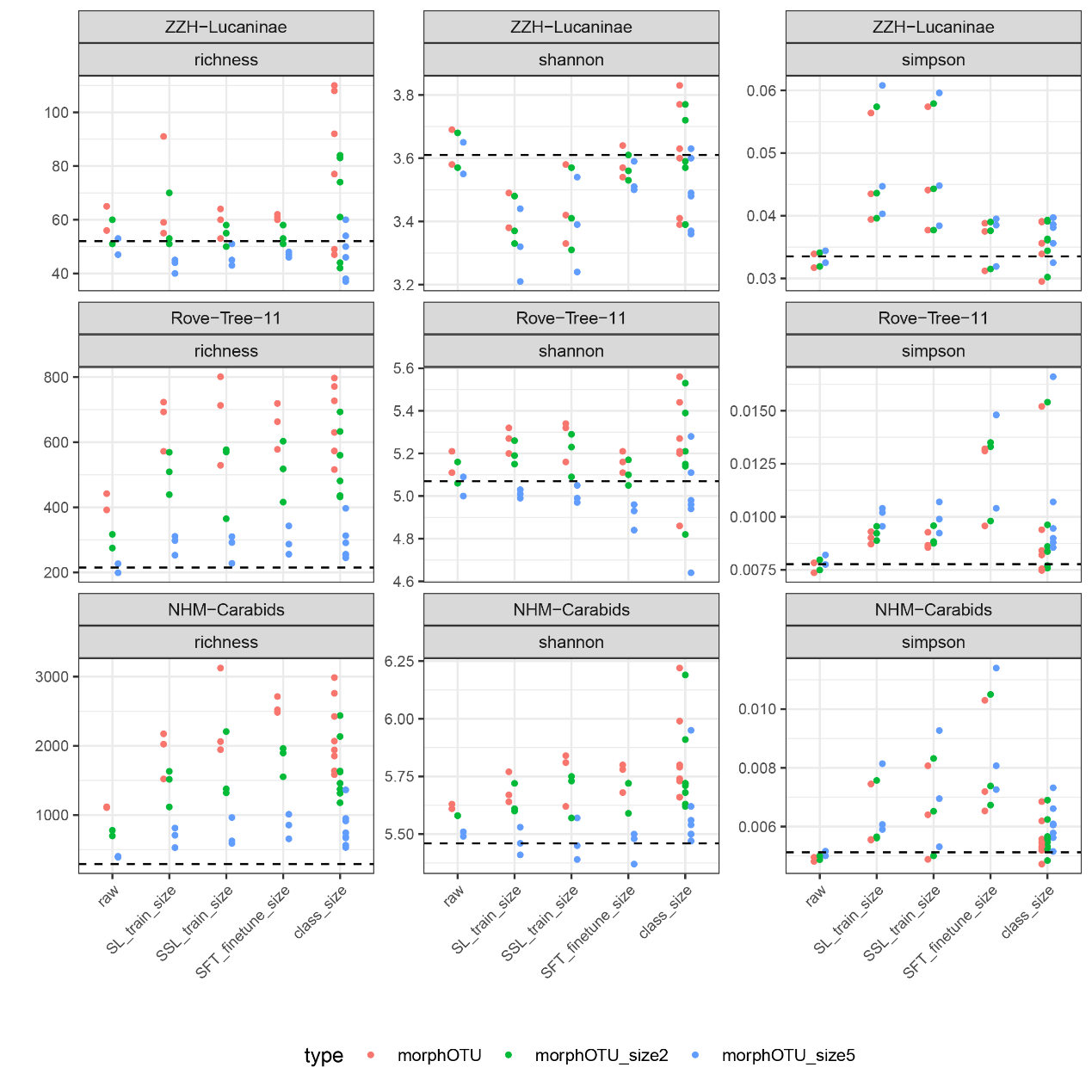

**Figure S18 | Estimated richness, Shannon diversity, and Simpson diversity of morphOTUs across datasets and embedding‑based OTU definitions.**

Columns correspond to three diversity metrics (richness, Shannon, Simpson), and rows correspond to the three datasets (ZZH–Lucaninae, Rove‑Tree‑11, NHM–Carabids). Within each panel, points represent diversity estimates computed under different embedding conditions (raw, SL, SSL, SSL‑train‑size, SSL‑finetune, SFT‑finetune, and class‑size‑based baselines). Colors indicate different OTU definitions: morphoOTU (red), morphoOTU_size2 (green), and morphoOTU_size5 (blue).

Dashed horizontal lines mark diversity values calculated directly from ground‑truth species labels, providing a reference for comparing embedding‑derived estimates. Results show systematic differences among OTU definitions, with finer OTU thresholds generally producing higher richness and more fragmented diversity patterns. Variation across embedding conditions reveals the sensitivity of diversity estimation to the geometry of the underlying embedding space.

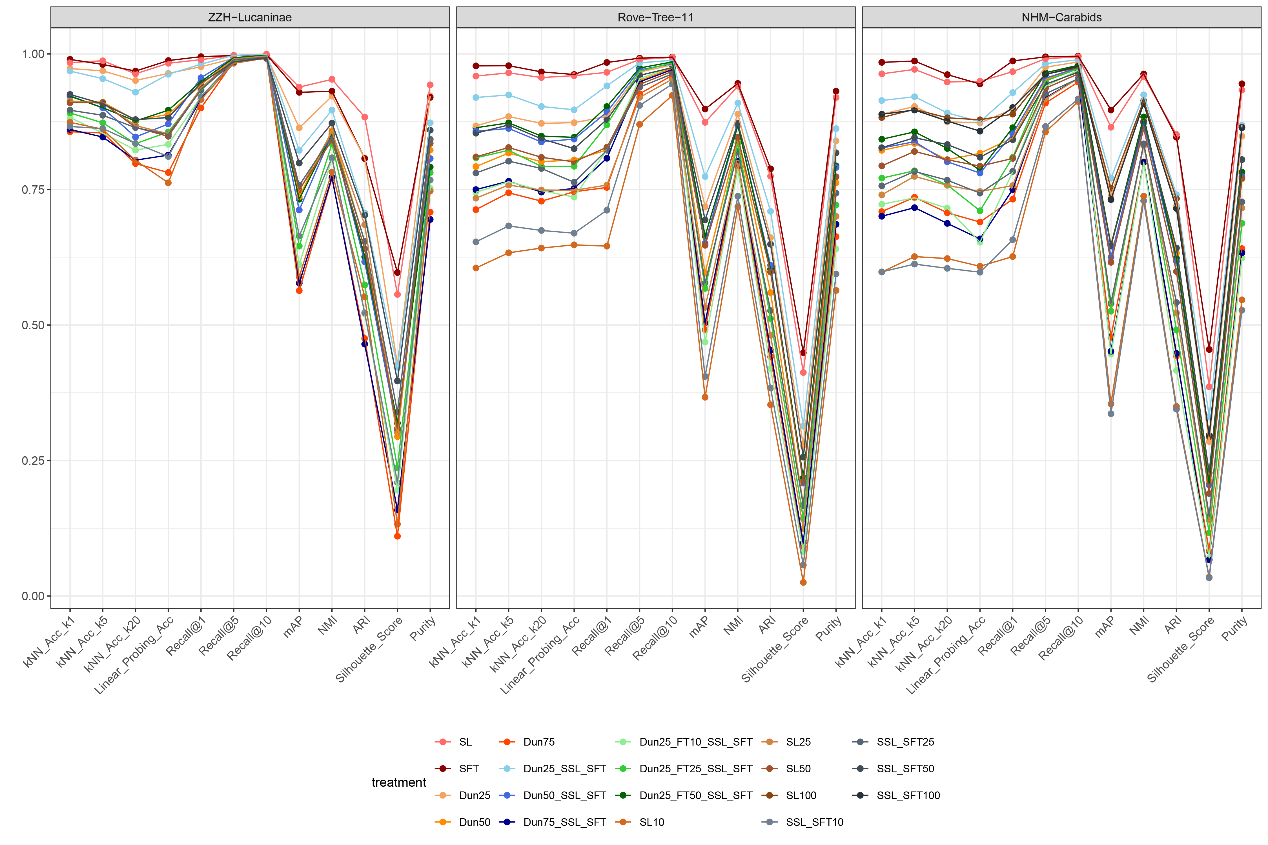

**Figure S19 | Performance of different training treatments across multiple embedding‑quality metrics for three datasets.**

Each panel corresponds to one dataset (ZZH–Lucaninae, Rove‑Tree‑11, NHM–Carabids). Within each panel, lines represent different training treatments, including supervised learning (SL), self‑supervised learning (SSL), supervised fine‑tuning (SFT), SSL+SFT combinations, and downsampled training‑set variants (Dun25–Dun100).

Metrics shown along the x‑axis include kNN classification accuracy at multiple neighborhood sizes, linear‑probe accuracy, recall at different thresholds, mean average precision (mAP), normalized mutual information (NMI), adjusted Rand index (ARI), silhouette score, and cluster purity.

Across datasets, treatments vary substantially in performance, with some methods showing consistent strength across most metrics, while others excel only in classification or clustering‑based measures. Patterns of divergence among treatments highlight how training regimes, fine‑tuning strategies, and data‑availability constraints shape the geometric and semantic quality of the resulting embedding spaces.

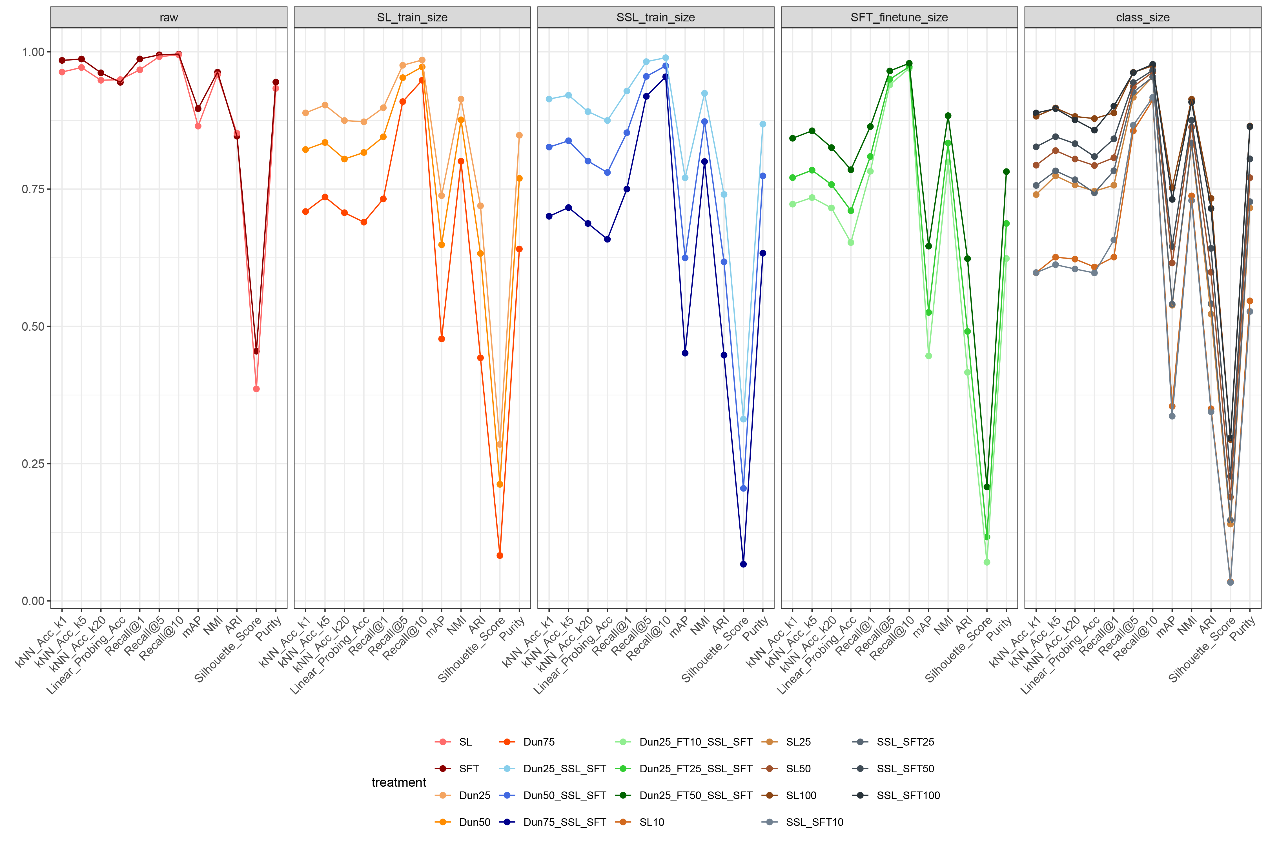

**Figure S20 | Performance of raw embeddings and multiple training treatments under different dataset‑size and class‑size conditions.**

Columns represent five conditions: raw embeddings, supervised learning with varying training‑set size (SL_train_size), self‑supervised learning with varying training‑set size (SSL_train_size), supervised fine‑tuning with varying sample size (SFT_finetune_size), and class‑size–controlled baselines (class_size). Within each panel, lines represent different treatments, including full‑data models, SSL+SFT pipelines, and downsampled training regimes (Dun25–Dun100).

Metrics along the x‑axis include kNN classification accuracy at multiple neighborhood sizes, linear‑probe accuracy, recall at multiple thresholds, mean average precision (mAP), normalized mutual information (NMI), adjusted Rand index (ARI), silhouette score, and cluster purity.

The results show clear impacts of dataset size and class‑size constraints on embedding quality. Training‑set reduction generally degrades classification‑style metrics more strongly than clustering‑based metrics, while fine‑tuning partially counteracts these effects. Class‑size baselines reveal how label imbalance and sampling structure influence downstream evaluation independent of model architecture.

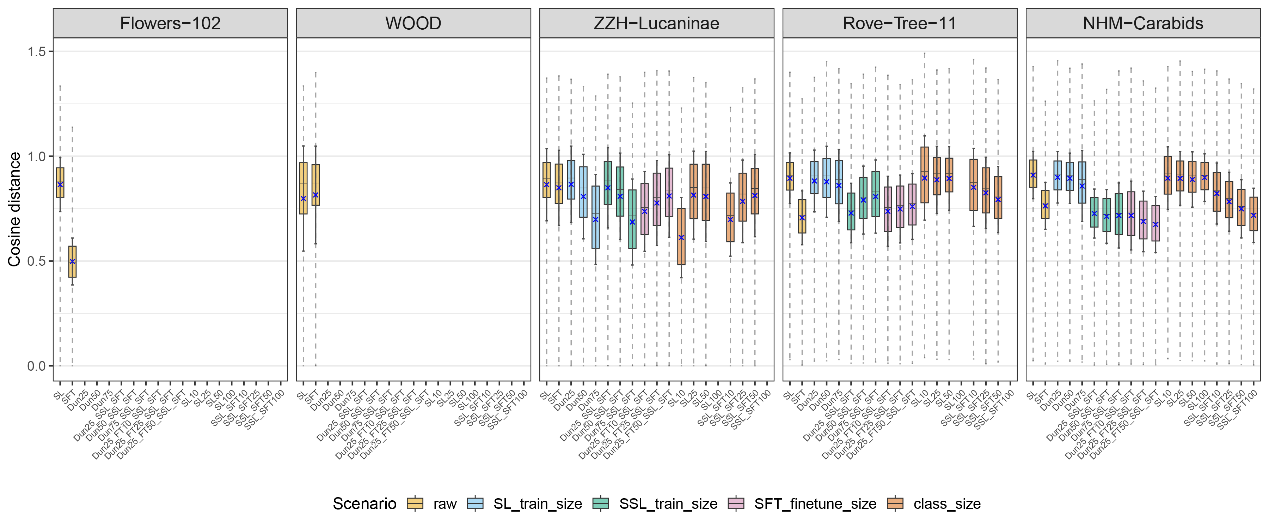

**Figure S21 | Boxplot summaries of cosine‑distance distributions across five datasets under different modeling scenarios.**

Each panel represents one dataset (Flowers‑102, WOOD, ZZH–Lucaninae, Rove‑Tree‑11, NHM–Carabids). Within each panel, boxplots show the distribution of pairwise cosine distances for a range of scenarios: raw embeddings, supervised learning with reduced training‑set sizes (SL_train_size), self‑supervised learning with reduced training‑set sizes (SSL_train_size), supervised fine‑tuning with reduced sample sizes (SFT_finetune_size), and class‑size–controlled baselines.

Boxes represent the interquartile range with medians, whiskers represent distributional spread, and blue × symbols mark the mean cosine distance.

Across datasets, distance distributions shift noticeably depending on training regime, data availability, and class‑size structure. Reduced‑data scenarios typically produce broader and higher‑variance distance distributions, while fine‑tuning and SSL‑based models tend to yield more compact embeddings. Differences among datasets highlight varying intrinsic morphological diversity and dataset difficulty.

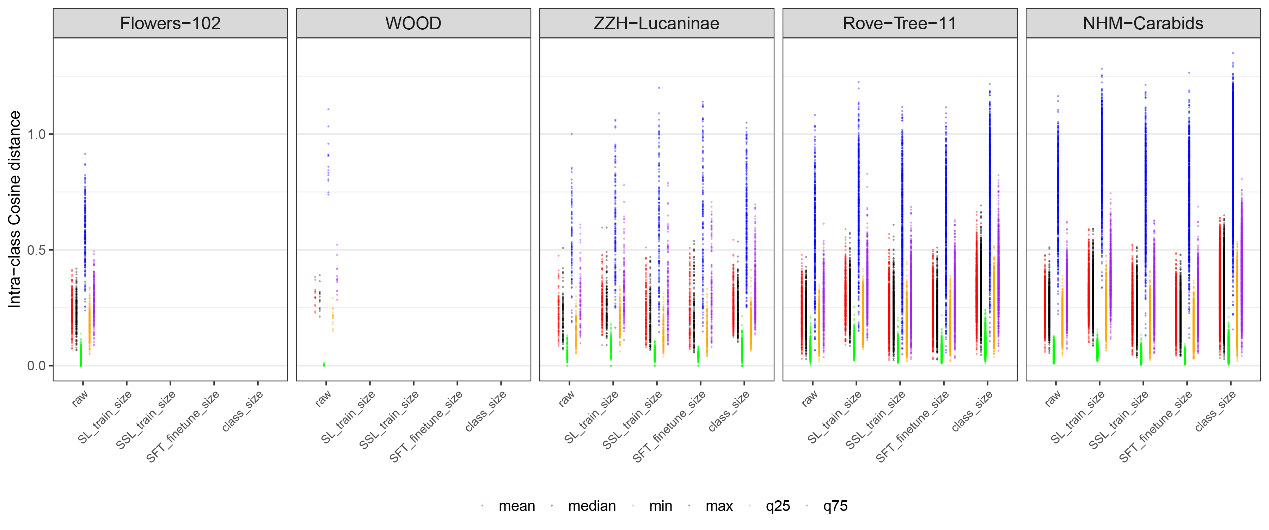

**Figure S22 | Summary statistics of intra‑class cosine distances under five modeling scenarios across five datasets.**

Panels correspond to different datasets (Flowers‑102, WOOD, ZZH–Lucaninae, Rove‑Tree‑11, NHM–Carabids). Within each panel, points represent intra‑class cosine‑distance summary statistics computed for multiple scenarios: raw embeddings, supervised learning with reduced training size (SL_train_size), self‑supervised learning with reduced training size (SSL_train_size), supervised fine‑tuning with limited samples (SFT_finetune_size), and class‑size–controlled baselines.

For each scenario, the distribution of intra‑class distances is summarized using mean (red), median (black), minimum (green), maximum (blue), and the first and third quartiles (yellow and purple).

Patterns differ strongly among datasets: some exhibit compact and low‑variance intra‑class structure, while others show wide spreads or high upper tails. Training‑set reduction and class‑size effects are visible as upward shifts in quartiles and maxima, whereas fine‑tuning often compresses intra‑class variability. These trends demonstrate how embedding quality is shaped jointly by dataset properties, model training regime, and sampling constraints.

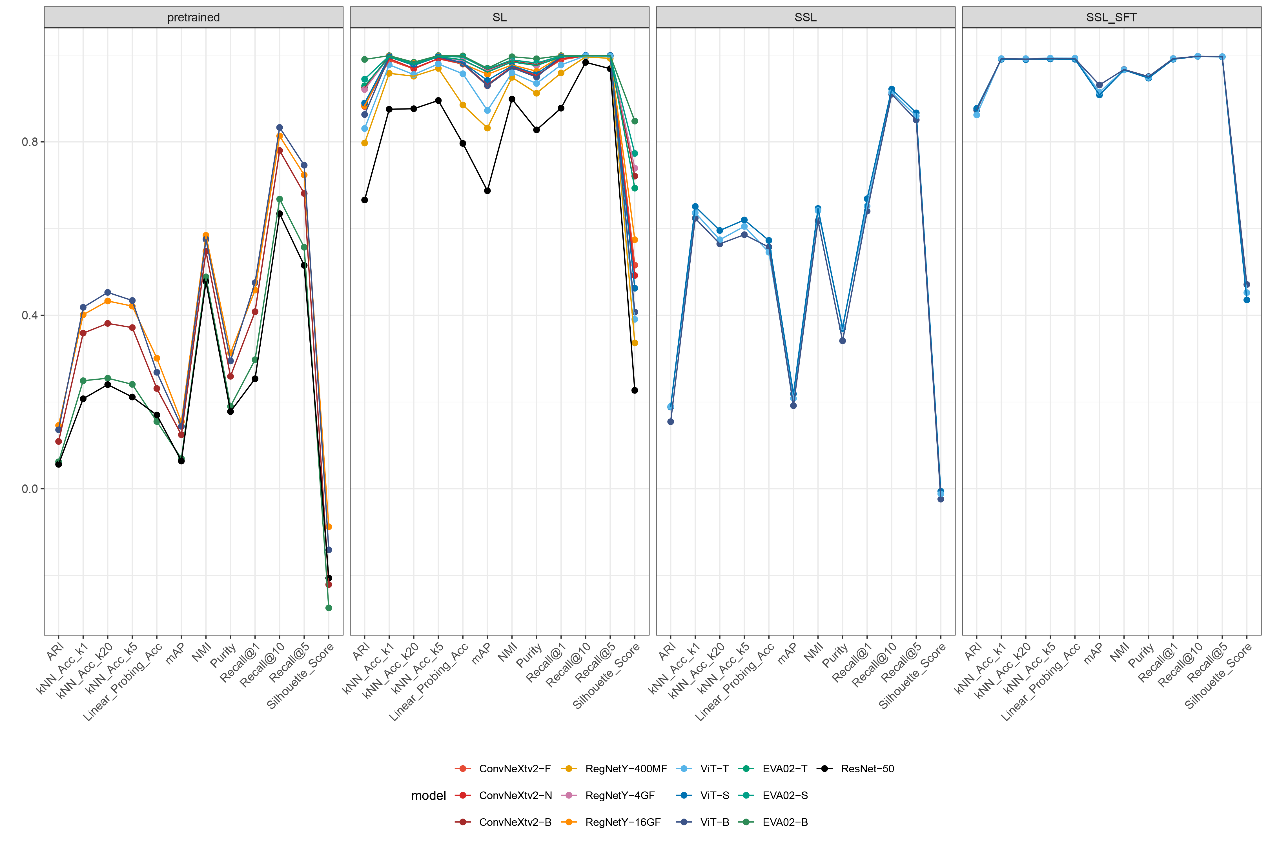

**Figure S23 | Performance of individual model architectures across multiple embedding‑quality metrics under four training regimes.**

Panels represent different training conditions: pretrained (no task‑specific training), supervised learning (SL), self‑supervised learning (SSL), and SSL followed by supervised fine‑tuning (SSL_SFT). Within each panel, lines correspond to different model architectures, including convolutional networks (ConvNeXtV2 family, ResNet‑50), transformer‑based models (ViT variants), and next‑generation hybrid architectures (RegNetY, EVA02).

Metrics along the x‑axis include adjusted Rand index (ARI), kNN accuracies at different neighborhood sizes, linear‑probe accuracy, mean average precision (mAP), purity, recall at multiple thresholds, normalized mutual information (NMI), and silhouette score.

Clear differences emerge between architectures and training regimes: SSL_SFT models consistently achieve high performance across most metrics, SL models show strong results but with greater variability across architectures, SSL‑only models excel in some clustering metrics, while pretrained models show broad performance gaps but retain model‑specific patterns. These results highlight how architectural design and training strategy jointly influence embedding geometry and downstream task utility.

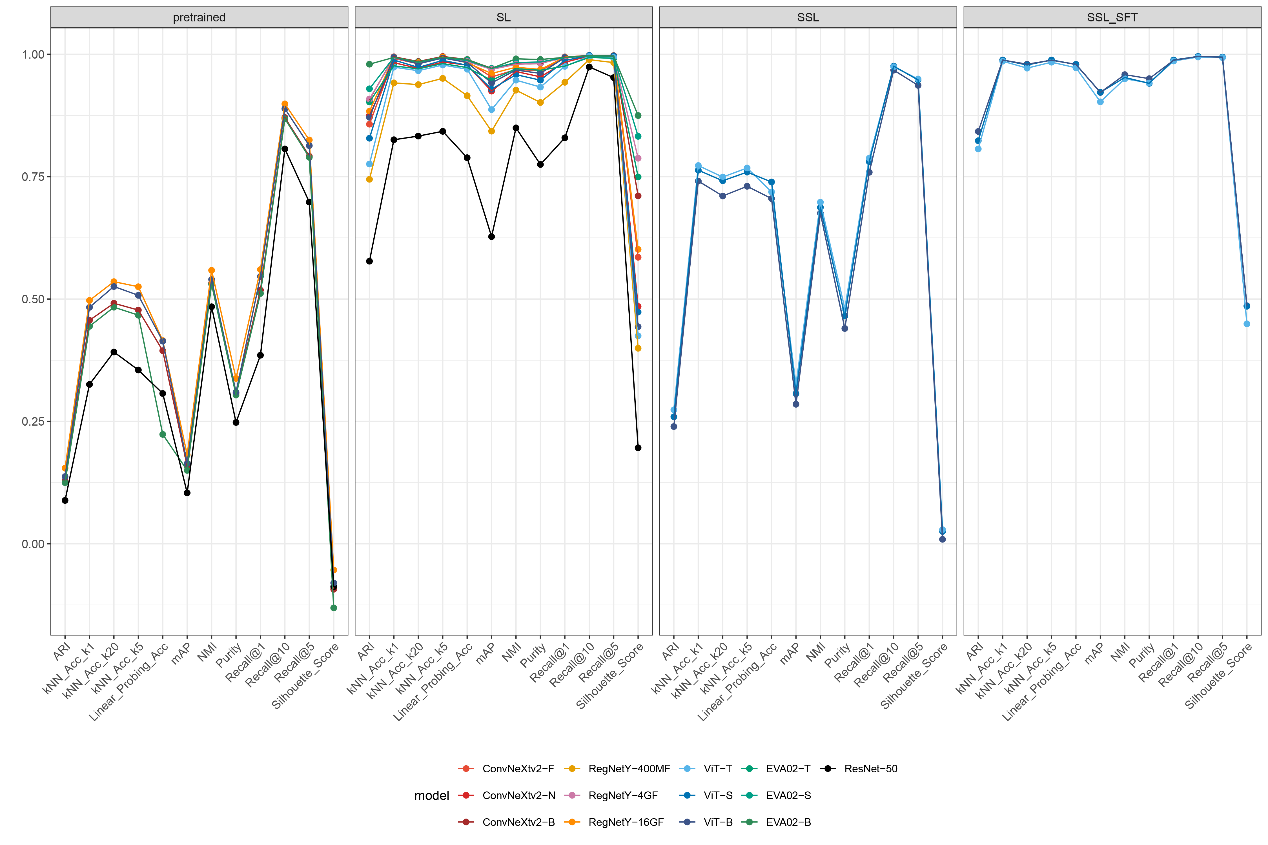

**Figure S24 | Comparison of embedding‑quality metrics across model architectures under four training conditions.**

Panels correspond to pretrained models, supervised learning (SL), self‑supervised learning (SSL), and self‑supervised learning followed by supervised fine‑tuning (SSL_SFT). Within each panel, lines represent different architectures, including ConvNeXtV2 variants, RegNetY models, ViT variants, EVA02 models, and ResNet‑50.

Metrics along the x‑axis include adjusted Rand index (ARI), kNN accuracies at multiple neighborhood sizes, linear‑probe accuracy, mean average precision (mAP), purity, recall at multiple thresholds, normalized mutual information (NMI), and silhouette score.

Across all training regimes, architectural differences strongly influence embedding quality. SSL_SFT consistently yields high scores with low variance across models, SL produces strong but architecture‑dependent performance, SSL captures structure well in clustering‑based metrics, and pretrained models show wide variation and generally lower accuracy. These trends highlight interactions between model capacity, representation bias, and training supervision in shaping embedding‑space geometry.

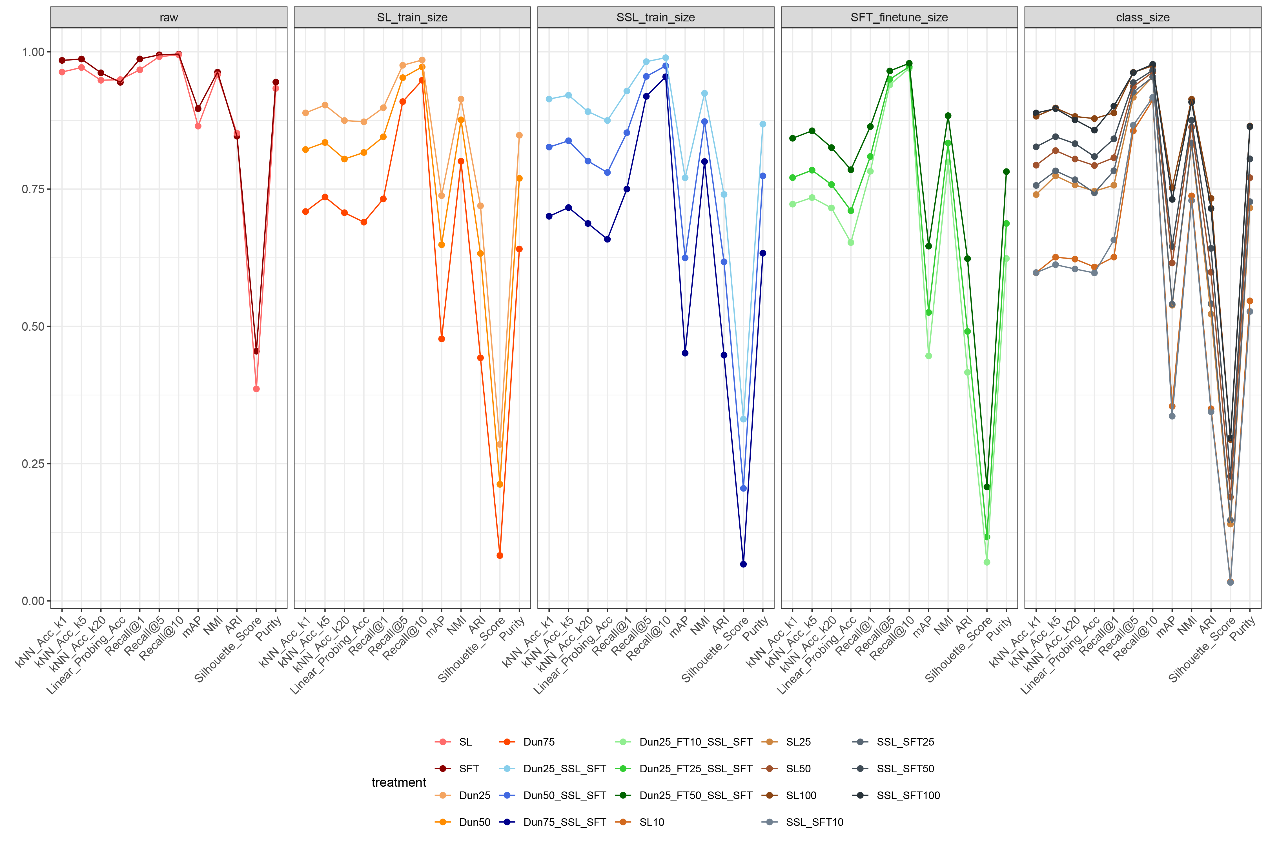

**Figure S25 | Performance of multiple training treatments across embedding‑quality metrics under five evaluation scenarios of NHM–Carabids.**

Columns correspond to raw embeddings, supervised learning with reduced training size (SL_train_size), self‑supervised learning with reduced training size (SSL_train_size), supervised fine‑tuning with limited sample size (SFT_finetune_size), and class‑size–controlled baselines (class_size). Within each panel, lines represent distinct treatments, including full‑data SL and SFT models, SSL‑based models, SSL–SFT pipelines, and downsampled training‑set variants (Dun25–Dun100).

Metrics along the x‑axis include kNN classification accuracy at several neighborhood sizes, linear‑probe accuracy, recall at different thresholds, mean average precision (mAP), normalized mutual information (NMI), adjusted Rand index (ARI), silhouette score, and cluster purity.

The results illustrate how training‑set availability and class‑size structure shape embedding quality. Reduced training sizes systematically lower performance across many metrics, with stronger effects on classification‑style measures than on clustering metrics. Fine‑tuning typically improves both accuracy and cluster structure, while class‑size baselines reveal the impact of label imbalance and sampling complexity. These patterns highlight the interaction between data scale, training supervision, and structural constraints in determining embedding performance.

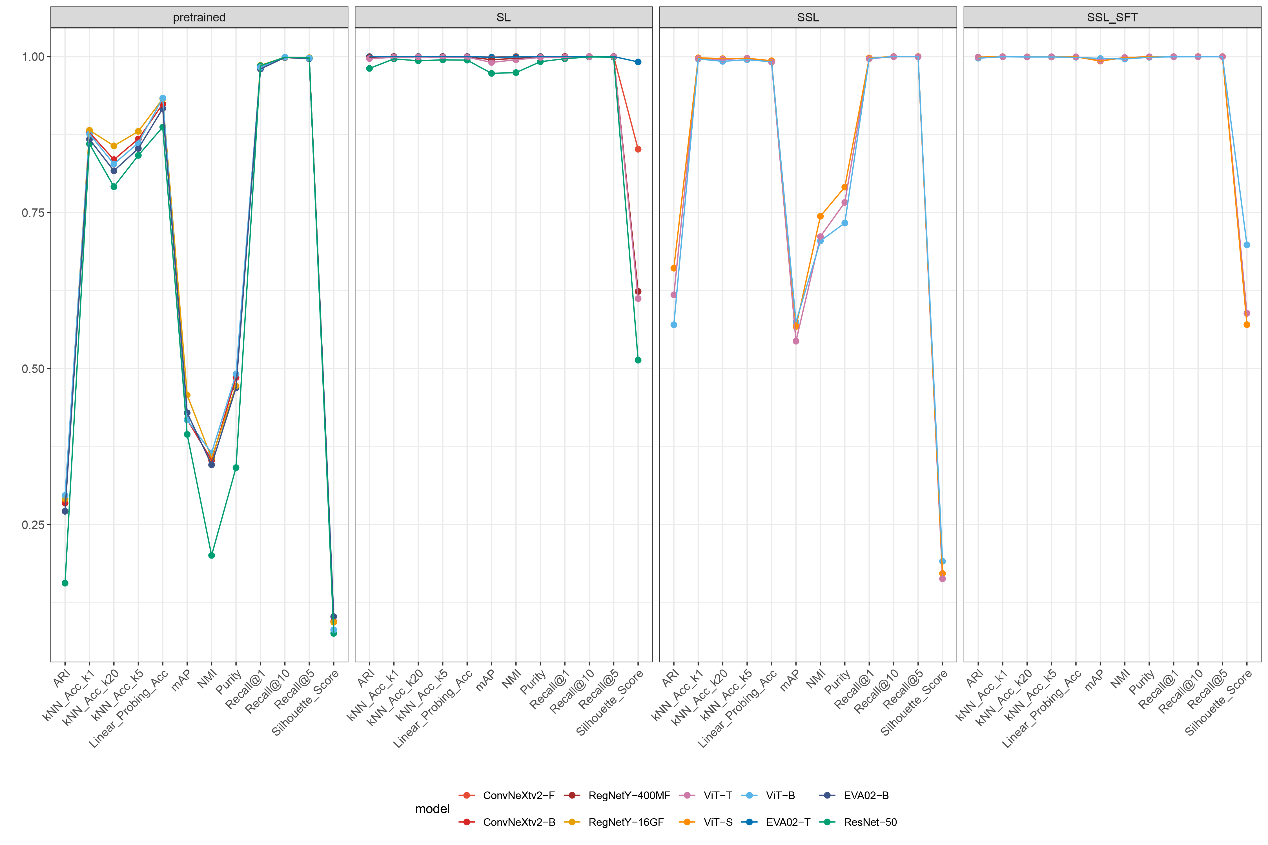

**Figure S26 | Performance of a subset of model architectures across multiple embedding‑quality metrics under four training conditions of WOOD.**

Panels represent pretrained models, supervised learning (SL), self‑supervised learning (SSL), and SSL followed by supervised fine‑tuning (SSL_SFT). Within each panel, lines correspond to different architectures, including ConvNeXtV2 variants, RegNetY models, ViT variants, EVA02 models, and ResNet‑50.

Metrics along the x‑axis include adjusted Rand index (ARI), kNN accuracies at different neighborhood sizes, linear‑probe accuracy, mean average precision (mAP), purity, recall at multiple thresholds, normalized mutual information (NMI), and silhouette score.

The simplified model set reveals consistent patterns across training regimes. SSL_SFT models achieve uniformly strong and stable scores, SL models show high performance with moderate architectural variation, SSL models excel in several clustering metrics while showing more variability in accuracy‑related measures, and pretrained models display wide performance dispersion reflecting their differing default representational biases. Overall, the results demonstrate how architecture and training strategy jointly determine the structure and utility of image‑embedding spaces.

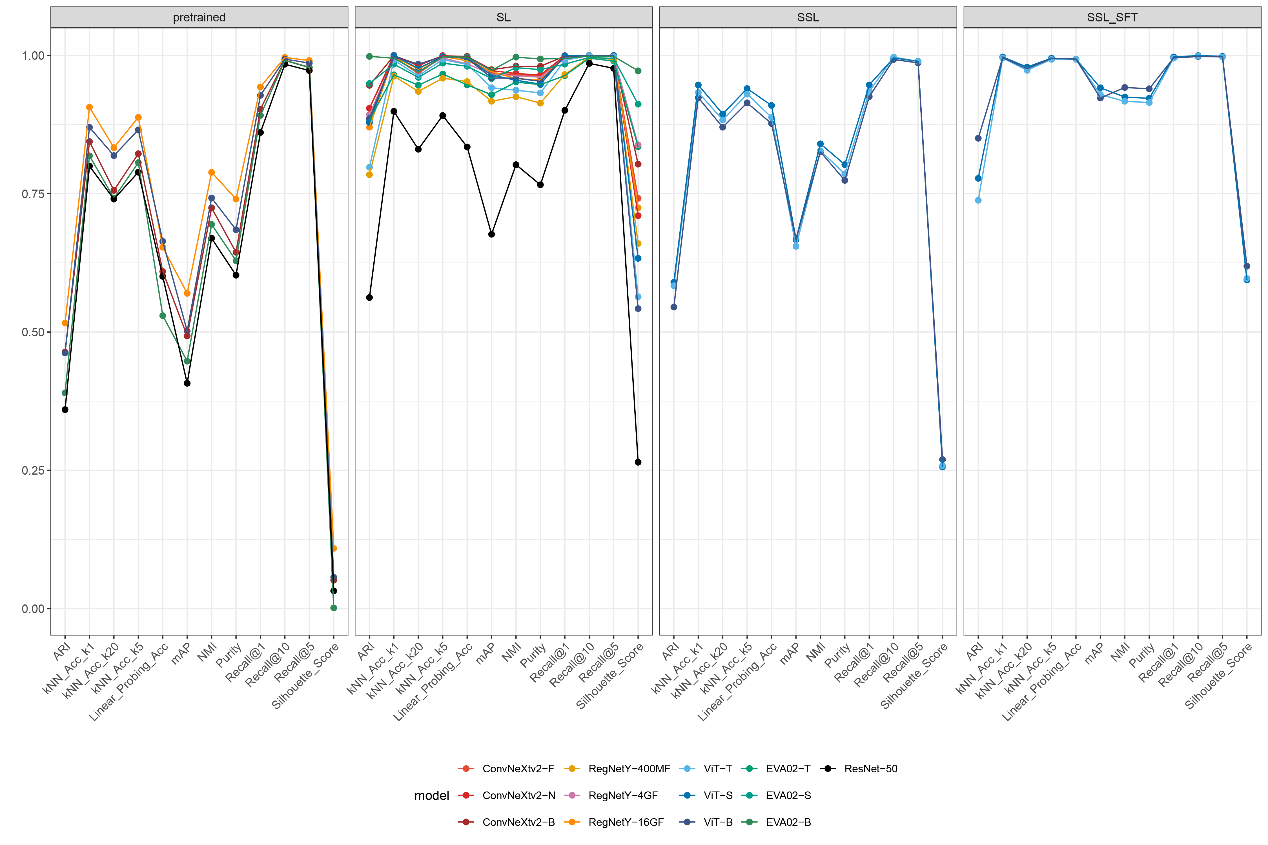

**Figure S27 | Performance of diverse model architectures across multiple embedding‑quality metrics under pretrained, supervised (SL), self‑supervised (SSL), and SSL‑plus‑supervised‑fine‑tuning (SSL_SFT) training conditions of ZHH-Lucaninae.**

Each panel corresponds to a different training regime. Within panels, lines represent individual model architectures, including ConvNeXtV2 variants, RegNetY models, ViT variants, EVA02 models, and ResNet‑50.

Metrics along the x‑axis include adjusted Rand index (ARI), kNN classification accuracies at multiple neighborhood sizes, linear‑probe accuracy, mean average precision (mAP), purity, recall at different thresholds, normalized mutual information (NMI), and silhouette score.

Across regimes, models show clear and consistent performance patterns. SSL_SFT models achieve uniformly high and stable scores across nearly all metrics. SL models exhibit strong performance with noticeable architecture‑dependent variability. SSL models capture clustering structure well but show greater dispersion across accuracy‑related metrics. Pretrained models vary widely in performance, revealing differences in inherent representational bias prior to domain‑specific training.

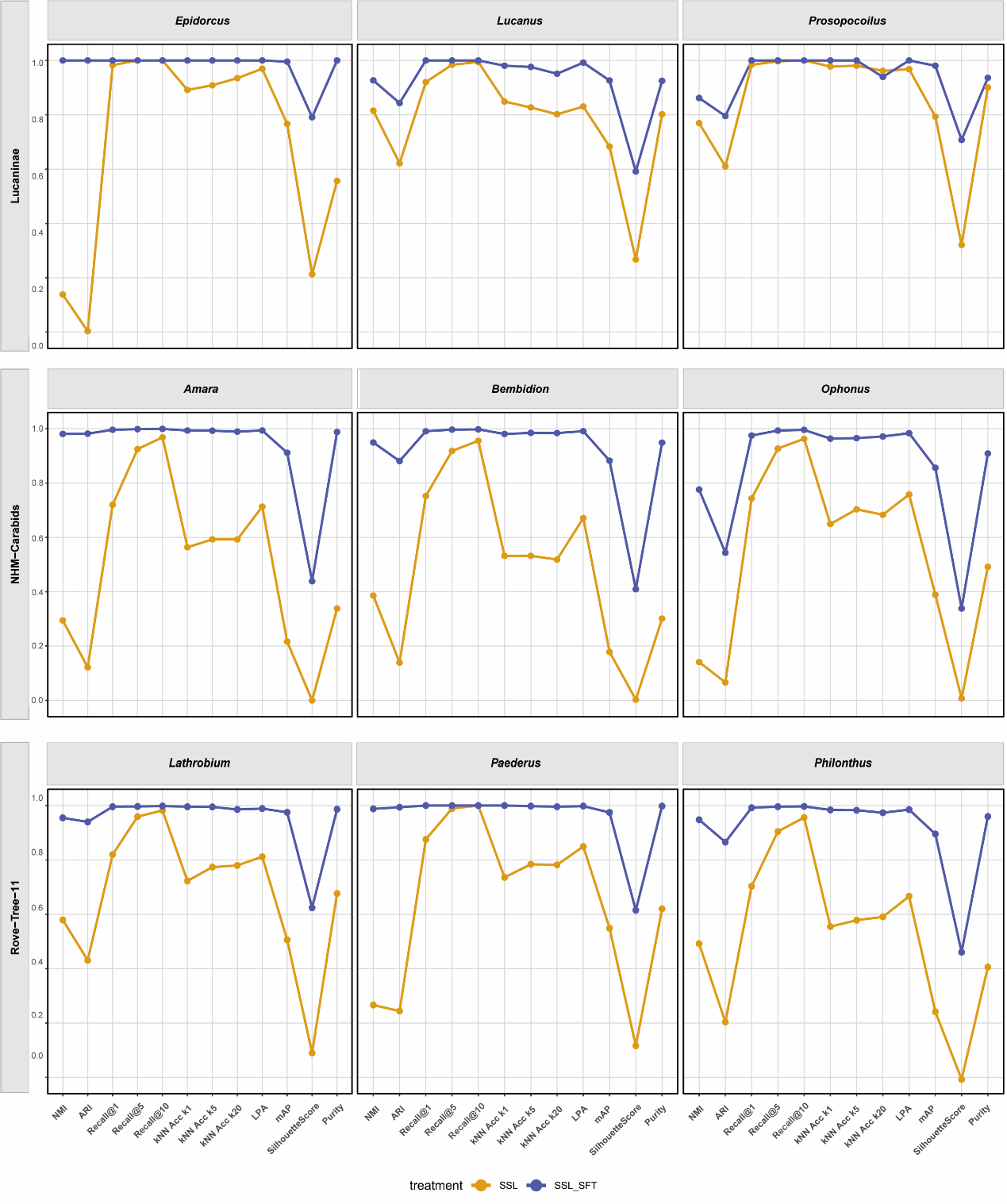

**Figure S28 | Genus‑level evaluation of morphOTU performance across nine closely related beetle groups**

Performance comparison of self‑supervised learning (SSL) and self‑supervised plus supervised fine‑tuning (SSL+SFT) across nine genus‑level subsets spanning three datasets: ZZH-Lucaninae (Epidorcus, Lucanus, Prosopocoilus), NHM-Carabids (Amara, Bembidion, Ophonus), and RowTree-1 (Lathrobium, Paederus, Philonthus). These genera represent groups of closely related and morphologically similar species that are often challenging in practical morphology‑based classification. Across all subsets, SSL+SFT (blue) consistently yields higher and more stable embedding performance than purely self‑supervised learning (orange), particularly under metrics sensitive to fine-scale phenotypic structure. The improved stability and separation achieved by SSL+SFT highlight OTU-Former's ability to resolve subtle morphological distinctions and historically problematic taxa, thereby supporting more reliable morphOTU formation in difficult species groups.

### **Supplementary Scripts**

### **clean_figs.py**

This script automatically cleans, standardizes, and resizes image files before saving them to a target directory. Its main operations are:

**1. Input Handling**

- Load all images from the input directory: temp2
- Only process files with extensions: .png, .jpg, .jpeg

**2. Filename Cleaning**

Each source filename is normalized as follows:

- Replace spaces → _
- Replace parentheses ()（） → _
- Remove characters: =, ', ,, .
- Strip leading/trailing underscores
- If the resulting name is empty → rename to "untitled"
- Standardize extension:
  - .jpg or .jpeg → .jpg
  - All other valid inputs → .png

**3. Ensuring Unique Filenames**

- If the cleaned name already exists in the output directory (images), append an index suffix: _1, _2, _3, …
- Maintain a set of used prefixes to avoid collisions.

**4. Image Resizing**

- Inspect the image’s short edge length.
- If short edge ≤ 512 px → keep original resolution.
- If short edge > 512 px → resize proportionally so that
  short edge = 512 px, using high‑quality LANCZOS interpolation.

**5. Color Mode Normalization**

- When saving as JPG, convert images to RGB if their mode is not RGB/L.

**6. Saving Processed Images**

- Save cleaned and resized images into the output directory: images
- Create the directory automatically if it does not already exist.

### **detect_bounding_box.py**

This script performs automatic object detection, segmentation, and cropping of biological specimen images. It identifies the largest object in each image, extracts its bounding box, optionally applies padding, and saves both the cropped image and its bounding‑box coordinates.

**1. Input Handling**

- Load all images from the specified input directory.
- Supported types: .png, .jpg, .jpeg, .bmp, .tiff, .tif
- Multi‑threaded processing enabled via ThreadPoolExecutor.

**2. Segmentation Methods**

The script supports three segmentation options:

1. Otsu thresholding
   - Convert to grayscale → Gaussian blur → Otsu binarization
   - Morphological dilation + erosion to refine mask
2. GrabCut segmentation
   - Initialize GrabCut with a rectangle covering most of the image
   - Extract foreground mask
   - Robust but may fail on some images
3. LAB-based adaptive thresholding (Default)
   - Convert image to LAB color space
   - Estimate background brightness from image corners
   - Apply inverse threshold on the L channel
   - Morphological opening to clean noise

**3. Object Detection**

- Find all contours in the segmentation mask.
- Select the largest contour as the target object.
- Compute an axis‑aligned bounding box using cv2.boundingRect.

**4. Bounding Box Expansion (Optional)**

Using a padding_ratio parameter:

- 1.0 = no padding (object fits the crop tightly)
- Lower values (e.g., 0.9) → add extra border around the object
- The padded box is clamped to image boundaries to avoid overflow.

**5. Crop Extraction**

- Crop the original image using the (optionally expanded) bounding box.
- If cropping fails (empty region), skip the file.

**6. Saving Outputs**

For each input image:

Cropped Image

- Saved using the specified output format: jpg / png / tif
- JPG images saved with quality = 95

Bounding Box Coordinates

- Saved as a text file (.txt) with:
- x y w h

in the coordinate frame of the processed image
(x, y = top-left corner; w, h = width and height).

**7. Command-line Options**

The script exposes the following arguments:

- --input_images_dir: folder containing input images
- --output_images_dir: destination folder for cropped outputs
- --method: segmentation method (Otsu, GrabCut, Lab, default: Lab)
- --out_image_format: output image format (jpg, png, tif)
- --padding_ratio: bounding box expansion factor (default: 1.0)
- --threads: number of parallel threads (default: 12)

**8. Additional Notes**

- Any segmentation failure or empty crop is skipped with a warning.
- Rotation logic for object alignment exists in the source code but is currently disabled.
- The script outputs both processed crops and corresponding .txt bounding box files for downstream tasks.

### **download_TreeOfLife-200M.py**

This script downloads images from the TreeOfLife‑200M dataset according to a user‑specified taxonomic filter (e.g., class/order/family/genus/species). It supports streaming from HuggingFace, local parquet processing, and multi‑threaded downloading while generating structured metadata for each image.

**1. Taxonomic Filtering**

The script filters dataset records by a specified taxonomic level and name:

- Supported levels: kingdom, phylum, class, order, family, genus, species
- Species filtering requires *"Genus species"* format.
- Records are matched via:
  - Direct field match (e.g., record['order'] == 'Coleoptera')
  - Fallback match within the taxonomic_path string

**2. Data Loading Options**

The script can obtain dataset metadata in three modes:

**(A) Local parquet file (recommended)**

- Processes large parquet files in streaming fashion (row‑group by row‑group) using PyArrow.
- Minimizes RAM usage.

**(B) Streaming from HuggingFace**

- No local parquet needed; downloads metadata record‑by‑record.
- Useful for distributed or low‑storage environments.

**(C) Automatic parquet shard download**

- Downloads all shard files from HuggingFace into a local directory (parquet_chunks/).
- Then processes them in streaming mode.

**3. Record Streaming and Filtering**

Records are read in batches (RowGroups or chunked batches), and only matching records pass to the downloader. Features:

- Chunk sizes adjustable (default: 10k rows)
- Multithreaded batch handling
- Batch aggregator (create_record_batcher) for efficient parallel downloading

**4. Image Downloading**

For each matched record:

1. Extract image URL and UUID
2. Generate a taxonomically structured filename

Format:

3. class_order_family_genus_species-uuid.ext
4. Download using HTTP streaming with robust error handling
5. Detect content type to confirm the response is an image
6. Save image into:
7. output_dir/images/
8. Log failed downloads separately (CSV) with error messages

Concurrent downloading uses up to --workers threads.

**5. Metadata Storage**

Two CSV files are produced:

- metadata.csv — successful downloads
- metadata.fail.csv — failed downloads, with error messages

Metadata fields include:

uuid, source_url, class, order, family, genus, species, scientific_name, img_type, publisher, etc.

Additionally, a file tracking processed UUIDs is maintained:

processed_uuids.txt

to avoid re‑downloading the same image.

**6. Filename Generation**

Filenames are constructed from available taxonomic fields:

- Spaces converted to underscores
- Missing levels skipped
- UUID appended to ensure uniqueness
- Non‑image extensions normalized to .jpg unless recognized

**7. Integrity Checking (Optional)**

Running with --validate:

- Scans all downloaded files
- Checks binary headers for JPEG/PNG/GIF signatures
- Reports number of corrupted files

**8. Command-line Arguments**

Key parameters include:

- --level: taxonomic level (e.g., class/order/family)
- --name: taxon name (e.g., “Insecta”, “Coleoptera”)
- --max-records: limit number of matched records
- --workers: concurrent download threads
- --output-dir: output folder
- --parquet: local parquet file for faster processing
- --stream: use HuggingFace streaming
- --log-level + --log-file: diagnostic logging options
- --validate: check integrity of downloaded images

**9. Summary of Output**

The script produces:

- Cleanly named image files in output_dir/images/
- metadata.csv (successful downloads)
- metadata.fail.csv (failed downloads + reasons)
- processed_uuids.txt (avoids duplicate downloads)
- Detailed logs (optional)

### **embeddings_tree.py**

This script computes pairwise embedding distances, summarizes their distribution, and constructs phylogeny‑style trees (UPGMA and/or Neighbor‑Joining) from image embeddings. It also supports bootstrap resampling to estimate clade support and produces partition scans, dendrograms, Newick trees, and cluster tables.

**1. Input & Configuration**

- Input: a CSV file where the first column is image ID and all remaining columns are numeric embeddings.
- CLI options allow:
  - Distance metric: cosine (default) or euclidean
  - Enable/disable UPGMA and Neighbor‑Joining (NJ) tree construction
  - Number of bootstrap replicates
  - Subsampling ratio for bootstrap feature resampling
  - Threshold ranges for cluster partition scans
  - Output directory and threading options

**2. Embedding Preparation**

Steps applied to the embeddings:

1. Load CSV, validate all feature columns are numeric
2. Extract embedding matrix X and corresponding IDs
3. Detect zero‑norm vectors and replace with a valid fallback
4. (Optional) L2 normalization
5. Compute pairwise distances using scipy.spatial.distance.pdist
6. Validate and symmetrize the distance matrix

Both condensed and square‑form distance matrices are used downstream.

**3. Distance Statistics & Visualization**

The script generates summary statistics of pairwise distances:

- Randomly samples up to max_distance_pairs distances
- Plots:
  - Histogram of distances
  - Log‑scaled histogram (positive distances only)
  - CDF plot
- Outputs are saved as PDF files in:
- out_dir/distance/

**4. Partition Scans (Threshold Sweep)**

The script evaluates cluster formation across a grid of distance cutoffs:

- For UPGMA:
  AgglomerativeClustering (linkage=average, precomputed distances)
- For NJ:
  Performs thresholding on linkage distances from NJ tree
- Produces:
  - CSV table: threshold vs. #clusters
  - Plot showing how clustering changes with cutoff

Partition definitions for each threshold are exported as tables listing cluster membership.

**5. UPGMA Workflow**

If --UPGMA is enabled:

1. Compute hierarchical clustering (scipy.cluster.hierarchy.linkage, average linkage)
2. Export:
   - Linkage partition tables
   - Newick tree
   - Partition scan results
3. Draw a partition + dendrogram panel:
   - Sample labels (left)
   - Cluster bars across thresholds (middle)
   - UPGMA dendrogram (right)

Optional bootstrap support:

- Resamples embedding dimensions
- Recomputes linkage per bootstrap replicate
- Counts clade occurrences
- Outputs:
  - Support‑annotated Newick tree
  - Bootstrap tree collection (*.trees)

**6. Neighbor‑Joining (NJ) Workflow**

If --NJ is enabled:

1. Build NJ tree (skbio.tree.nj) from validated distance matrix
2. Optionally midpoint‑root the resulting tree
3. Export:
   - Newick tree
   - Partition scan
   - Partition tables
   - Partition+dendrogram visualization
4. Optional bootstrap:
   - Subsample embedding features
   - Build NJ tree per replicate
   - Compute clade support
   - Output support‑annotated NJ tree + bootstrap tree file

**7. Unified Partition + Tree Visualization**

The script generates high‑resolution PDF panels combining:

- Tip labels
- Partition blocks across cutoffs
- Dendrogram alignment
- Optional bootstrap support annotations

This provides a visualization linking distance‑based tree structure and cluster stability across thresholds.

**8. Output Structure**

Running the script produces the following structure:

ebnf

out_dir/

distance/

distance_hist_*.pdf

distance_cum_*.pdf

UPGMA/

*.csv	# partition scans

partitions/

partition_*.csv

partition_*.tips.csv

UPGMA_*.nwk

UPGMA_*.pdf

UPGMA_*_bootstrap.*	# optional

NJ/

*.csv

partitions/

...

NJ_*.nwk

NJ_*.pdf

NJ_*_bootstrap.*	# optional

**9. Main Workflow**

Running the script:

1. Loads embeddings
2. Computes distance matrices
3. Generates histograms
4. Computes trees (UPGMA, NJ, or both)
5. Runs partition scans
6. Generates plots, Newick trees, cluster tables
7. (Optional) Performs bootstrap replicates
8. Saves all results to out_dir

### **autogluon_training.py**

This script extracts image embeddings using either a timm backbone (ConvNeXtV2, ViT, etc.) or a fine‑tuned AutoGluon multimodal model. It supports feature extraction, optional fine‑tuning with focal loss, image quality checks, duplicate filename protection, UMAP visualization, and saving embeddings in a standardized CSV format.

**1. Input Handling**

The script processes images from a user‑specified directory:

- Supported extensions: jpg, jpeg, png, tif, tiff, bmp, webp
- Ensures filenames (without suffix) are globally unique
- Produces a DataFrame listing image paths and stems

**2. Embedding Backends**

Two embedding strategies are supported:

(A) Raw timm backbone (default)

- Loads the specified model from timm (e.g., convnextv2_femto)
- Removes classification head (num_classes = 0)
- Uses timm’s built‑in preprocessing pipeline
- Extracts embeddings in batches using GPU/CPU/MPS

(B) AutoGluon fine‑tuning

Triggered when either:

- --train_data is provided
- --load_AutogluonModel is provided

Features:

- Fine-tunes a backbone (timm‑based) with optional:
  - Focal loss for class imbalance
  - Custom gamma parameter
  - Augmentation control
- Saves processed training CSV and trained model checkpoint directory
- Can later reload trained models with --load_AutogluonModel

Embeddings from a trained model are obtained via:

predictor.extract_embedding(...)

**3. Device Selection**

Automatically selects the best available device:

- CUDA → GPU
- MPS → Apple Silicon
- CPU fallback

User can override via --device.

**4. Embedding Extraction**

For either timm or AutoGluon:

1. Build a dataset of images
2. Use DataLoader for efficient batching
3. Run forward pass to extract embeddings
4. Combine all batches into a single array
5. Save embeddings as a standardized CSV:
6. image, E0000, E0001, ...

**5. Training Pipeline (Optional)**

If the user supplies training data:

**Input format**

CSV with:

image, label, and a directory containing the referenced image files.

**Workflow**

- Load and resolve training paths
- Compute optional focal‑loss class weights
- Build AutoGluon hyperparameters
- Train for up to:
  - --max_epochs, or
  - --time_limit (hours)

Model outputs stored under:

out_dir/AutogluonModels/<base_model>/

**6. UMAP Visualization (Optional)**

The script can visualize:

- Training embeddings (--visualize_train_data)
- Test/inference embeddings mapped via a label CSV (--visualize_test_data)

Features:

- Automatic L2 normalization
- Configurable UMAP parameters (neighbors, metric, min_dist)
- Automatic label color palette selection (tab10/tab20/husl)
- Optional label count limiting (--visualize_class_number)
- Saves high‑resolution PDF of UMAP plot

Outputs include:

umap.train.pdf

umap.pdf

**7. Output Files**

Running the script can generate:

- embeddings.csv — embeddings for inference images
- embeddings.train.csv — embeddings for training images
- UMAP plots (optional)
- AutoGluon model directory (if trained)
- Processed training CSV for reproducibility

All saved under:

<out_dir>/

**8. Command-line Options (High-level)**

- Image input: --input_images_dir
- Backbone: --base_model
- Training data: --train_data, --train_images_dir
- Load model: --load_AutogluonModel
- Loss/augmentation: --Focal_Loss, --disable_augment, etc.
- Extraction parameters: --batch_size, --num_workers
- Visualization: UMAP-related parameters
- Device selection: --device
- Output directory: --out_dir

**9. Main Workflow**

1. Validate and load images
2. Load or train prediction model
3. Extract embeddings
4. Save CSV
5. Optionally perform UMAP visualizations
6. Finish with a summary message

### **GradCam_heatmap.py**

This script generates Class Activation Map (CAM) visualizations—including GradCAM, GradCAM++, ScoreCAM, EigenCAM, LayerCAM, and AblationCAM—for deep learning classification models. It supports both timm backbones and AutoGluon MultiModalPredictor models. Outputs include overlaid heatmaps, raw CAM arrays, and a summary CSV.

**1. Input & Label Handling**

- Requires a CSV file containing image and label columns.
- --input_images_dir specifies the directory where images are stored.
- Optional limit on processed images via --max_images.

**2. Model Loading Options**

Two model-loading pathways:

(A) AutoGluon MultiModalPredictor

- Load with --load_AutogluonModel /path/to/model
- Internally extracts the PyTorch model for CAM computation.

(B) timm Backbone

- Specify --base_model (e.g., convnextv2_femto, vit_base_patch16_224)
- Optional features:
  - --checkpoint_path to load custom weights
  - --no_pretrained to avoid pre-trained weights
  - --num_classes override
- Architecture inference:
  - --cnn or --vit, or automatic inference from model name

**3. CAM Methods**

Supported CAM algorithms:

gradcam

gradcampp

layercam

ablationcam

scorecam

eigencam

- Implemented via the pytorch-grad-cam library
- Properly handles CNN and ViT architectures (including token reshaping for ViT)

**4. Target Layer Selection**

CAM requires selecting a target feature layer.

Options:

- --target_layer_name BLOCK.NAME (e.g., blocks.11.norm1)
- Automatic defaults:
  - CNN → last Conv2d layer
  - ViT → last block's normalization layer

AblationCAM for ViT also injects AblationLayerVit() when needed.

**5. Preprocessing & Display Transforms**

- Uses timm’s data-config system (resolve_model_data_config) to build:
  - Preprocess transform — resize → center crop → normalize
  - Display transform — resize → center crop (for visualization)
- Ensures CAM overlays align with the model’s effective input size.

**6. CAM Computation Workflow**

For each image:

1. Load + preprocess image
2. Forward pass to obtain logits, prediction index, and probability
3. Compute CAM heatmap for the predicted class
4. Normalize heatmap to [0, 1]
5. Resize CAM to original image resolution
6. Blend with the original image (image_weight, default = 0.5)
7. Generate side‑by‑side figure:
   - Left: raw image
   - Right: CAM overlay
8. Save outputs in:
9. out_dir/figures/
10. out_dir/arrays/

Optional: --save_npy writes normalized CAM arrays as .npy.

**7. Output Summary**

The script saves a cam_summary.csv, containing for each image:

- image filename
- ground‑truth label
- predicted class index
- predicted class probability
- path to CAM figure
- path to .npy file (optional)

**8. Device Selection**

Supports:

- CUDA (--device cuda)
- Apple MPS (--device mps)
- CPU fallback

Automatically warns and falls back when selected device unavailable.

**9. Logging & Verbosity**

- --log_level controls verbosity: DEBUG / INFO / WARNING / ERROR
- Periodic progress updates every 10 images
- Logs missing images or errors but continues processing

**10. Command‑Line Options (High-level)**

- Model selection:
  - --load_AutogluonModel
  - --base_model
  - --cnn or --vit
- CAM method:
  - --cam
  - --target_layer_name
- Device:
  - --device
- Output:
  - --out_dir
  - --fig_format
  - --save_npy
- Performance:
  - --cam_batch_size
  - --max_images

**11. Main Workflow**

1. Parse arguments
2. Load label CSV
3. Load model (AutoGluon or timm)
4. Build transforms
5. Determine target layer & initialize CAM extractor
6. Prepare output directories
7. Process each image and generate a heatmap
8. Write summary CSV
9. Finish with a log message

### **SSL.py**

This script provides a full pipeline for self-supervised learning (SSL), ArcFace metric finetuning, and embedding extraction, integrated with comprehensive evaluation metrics, UMAP visualization, attention‑pool finetuning, checkpoint management, and multi-crop ViT feature learning.

It is modularized into three major modes:

- --mode pretrain: iBOT‑style SSL pretraining
- --mode finetune: ArcFace metric-learning finetuning
- --mode extract: Embedding extraction + metrics + visualization

**1. Self-Supervised Pretraining (iBOT-style)**

Activated via:

--mode pretrain

1.1 Multi‑crop SSL training

- Two global crops + several local crops
- Augmentations include:
  - RandomResizedCrop
  - 180° rotation
  - horizontal & vertical flips
  - Gaussian blur & Color jitter
  - grayscale perturbation
- ViT backbone from timm, dynamically size‑aligned

1.2 Student–Teacher EMA Framework

- Student: full ViT with projector
- Teacher: EMA of student (momentum schedule)
- Loss components:
  - Global loss: similarity between student × teacher global views
  - Local loss: local‑to‑global contrastive terms
  - Masked‑token loss: reconstruction of teacher patch tokens
- Temperature warmup scheduling for both networks

1.3 Learning rate & temperature schedules

- Cosine LR decay
- Student & teacher temperature curves
- Momentum curve for EMA updates

1.4 Instant Metrics Logging (Lightweight)

Logged every N steps:

- losses (total/global/local/mask)
- gradient norm
- feature standard deviation
- embedding norm
- CLS token norm
- teacher center norm
- cosine similarity (student vs teacher)
- learning rate

Saved in: logs/instant_metrics.pretrain.csv

1.5 Comprehensive Metrics (Every few epochs)

Computes full evaluation metrics on train/test CSVs (optional):

- NMI, ARI
- Recall@1/5/10
- kNN‑Acc@1/5/20
- Linear Probing Accuracy (optional)
- Silhouette Score
- mAP (retrieval)
- Purity
- ArcFace‑specific metrics (if needed)

Written to: logs/metrics.pretrain.csv

1.6 Checkpointing

- Saves SSL_epoch_xxxx.pth
- Maintains SSL_latest.pth
- Deletes old checkpoints, keeping the last N

**2. ArcFace Metric-Learning Finetuning**

Activated via:

--mode finetune

2.1 Loading SSL pretrained weights

- Automatically adapts key names between EVA/ViT/ConvNeXt variants
- Freezes first ~70% of ViT blocks
- Replaces projector with a metric head:
- Linear → ReLU → Linear (embedding_dim)

2.2 ArcFace loss

Using pytorch-metric-learning:

- Margin = 28.6
- Scale = 64
- Embedding dimension defined by --metric_embed_dim
- Per-class trainable ArcFace weights

2.3 Instant metrics (lightweight)

Logged to: logs/instant_metrics.finetune.csv

2.4 Comprehensive metrics (optional)

Same as SSL mode, but saved in: logs/metrics.arcface.csv

2.5 Checkpoint management

- Saves arcface_epoch_xxxx.pth
- Maintains arcface_latest.pth

**3. Embedding Extraction Mode**

Activated via:

--mode extract

Provides multiple extraction modes:

3.1 Extraction sources

- SSL or ArcFace checkpoint
- Optional attention‑pool finetuning
- CSV‑based extraction for labeled datasets
- Folder‑level extraction for unlabeled images

3.2 Token extraction modes

--token_mode cls

--token_mode patch-topk

--token_mode attention-pool

(A) CLS token

- Use the projector or raw CLS features

(B) Patch‑TopK

- Extract top‑K patch embeddings (based on L2 norm)
- Apply PCA down to 256/512/1024 dimensions as needed

(C) Attention-pool

- Lightweight / multihead / gated pooling
- Supports automatic finetuning of attention query vector (few epochs)
- Saves finetuned checkpoint for reuse

3.3 CSV Extraction Output

Produces:

embeddings.train.csv, embeddings.test.csv, cam_summary.csv (if applicable)

logs/metrics.extract.csv

UMAP plots (optional)

**3.4 Folder Extraction Output**

When no CSV is provided:

- Extract embeddings for all images in the folder
- Compute unsupervised metrics (Silhouette)
- Save:
- embeddings.csv
- metrics.extract.csv
- images.processed.csv

**4. Metrics Implemented**

The script includes a comprehensive suite of evaluation metrics:

Clustering

- NMI
- ARI
- Purity

Retrieval

- Recall@K
- Mean Average Precision (mAP)

Classification

- 5‑fold Linear Probing Accuracy
- kNN Accuracy@K

Embedding Quality

- Silhouette Score
- Feature norm / CLS norm
- Gradient norm during training
- Teacher center norm

ArcFace‑specific

- Intra‑class variance
- Inter‑class cluster distance
- Embedding norm mean/std

**5. Visualization**

Multiple UMAP pipelines support:

- Train / test separation
- Top-N label filtering
- Automatic color‑palette adjustment
- Supports large class counts (HUSL palette)

Outputs saved under: logs/umap.*.pdf

**6. Checkpoint Adaptation Utilities**

Converts pretrained keys between:

- EVA <→ Standard ViT
- Swin <→ Standard
- ConvNeXt <→ Standard

Ensures compatibility across different backbones.

**7. Extra Features**

- PCA dimensionality reduction with safety checks
- Top‑K patch embedding extraction
- Automatic patch-size alignment
- Multi-crop dataset for SSL
- Unified instant and enhanced logging systems
- TeeLogger: mirrors stdout to log file while removing tqdm artifacts
- Auto‑detect latest checkpoints for finetune/extract modes

**8. Main Script Workflow**

Depending on mode:

PRETRAIN

1. Build dataset & augmentations
2. Initialize student & teacher models
3. Train for N epochs
4. Log metrics (instant + comprehensive)
5. Save checkpoints

FINETUNE

1. Load SSL checkpoint
2. Build ArcFace model
3. Train with metric loss
4. Save checkpoints + metrics

EXTRACT

1. Load checkpoint
2. Extract embeddings (cls / patch-topk / attention-pool)
3. Compute metrics
4. Save embeddings + UMAP plots

### **split_dataset.py**

This script performs dataset splitting for classification tasks. It supports two regimes:

- Ratio-based splitting (percentage-level control)
- Count-based splitting (absolute sample-count control)

Both modes allow separation into:

- train set (known classes)
- test.known set (known classes for closed-set evaluation)
- test.unknown set (unknown classes for open-set evaluation)

Additionally, the script exports class distribution reports for reproducibility.

**1. Input & Setup**

The script expects a CSV file containing:

image,label

Key initial steps:

- Load complete dataset
- Compute global per-class counts
- Save class statistics under:

out_dir/class_count/class.count

**2. Two Splitting Modes**

A. Ratio Mode (--mode ratio)

Used for standard train/test splits based on ratios.

**2.1 Unknown-class selection (Open-set)**

- Controlled by:
- --unknown_test_classes_ratio
- Randomly shuffles class order
- Accumulates entire classes until the desired *ratio of total samples* is reached
- Extracts these as test.unknown.csv

**2.2 Known-class split (Closed-set)**

- Remaining samples form the known subset
- Stratified sampling per-class using:
- train = group.sample(frac = 1 − known_test_classes_ratio)
- test_known = complement
- Saves:
  - train.csv
  - test.known.csv
  - Count files under class_count/

B. Count Mode (--mode count)

Used for fine-grained control of sample numbers.

**2.3 Unknown-class selection by count**

- Controlled by:
- --unknown_test_classes_count
- Accumulates full classes until reaching the required count
- Saves as **test.unknown.csv**

**2.4 Known test split by count**

- Controlled by:
- --known_test_classes_count
- Iterates class-by-class and takes samples sequentially until the total number matches the target
- Saved to test.known.csv

**2.5 Train set filtering via per-class min/max**

Using only remaining samples:

- --min_count_per_class: classes with fewer than this number are excluded
- --max_count_per_class: caps the number of training samples per class
- Saves:
- train.csv
- class.train.count

**3. Output Structure**

All output files saved under:

stylus

out_dir/

train.csv

test.known.csv (if applicable)

test.unknown.csv (if applicable)

class_count/

class.count

class.train.count

class.test.known.count

class.test.unknown.count

**4. Additional Features**

- Full reproducibility through --seed
- Strict validation that ratio-mode parameters cannot be mixed with count-mode parameters
- Automatic cleaning and ordering of per-class statistics
- Informative printouts showing the number of samples and resulting splits

**5. Main Workflow**

1. Parse arguments
2. Load raw CSV
3. Save global class statistics
4. Depending on mode (ratio or count), perform:
   - Unknown-class selection
   - Known-class splitting
   - Train-set construction
5. Save final CSV files
6. Save class distribution reports
7. Print summary
